# Inferring mammalian tissue-specific regulatory conservation by predicting tissue-specific differences in open chromatin

**DOI:** 10.1101/2020.12.04.410795

**Authors:** Irene M. Kaplow, Daniel E. Schäffer, Morgan E. Wirthlin, Alyssa J. Lawler, Ashley R. Brown, Michael Kleyman, Andreas R. Pfenning

**Author notes:** Corresponding authors: Irene M. Kaplow; Andreas R. Pfenning.

## Abstract

**Background:** Evolutionary conservation is an invaluable tool for inferring functional significance in the genome, including regions that are crucial across many species and those that have undergone convergent evolution. Computational methods to test for sequence conservation are dominated by algorithms that examine the ability of one or more nucleotides to align across large evolutionary distances. While these nucleotide alignment-based approaches have proven powerful for protein-coding genes and some non-coding elements, they fail to capture conservation at many enhancers, distal regulatory elements that control spatio-temporal patterns of gene expression. The function of enhancers is governed by a complex, often tissue- and cell type-specific, code that links combinations of transcription factor binding sites and other regulation-related sequence patterns to regulatory activity. Thus, function of orthologous enhancer regions can be conserved across large evolutionary distances, even when nucleotide turnover is high.

**Results:** We present a new machine learning-based approach for evaluating enhancer conservation that leverages the combinatorial sequence code of enhancer activity rather than relying on the alignment of individual nucleotides. We first train a convolutional neural network model that is able to predict tissue-specific open chromatin, a proxy for enhancer activity, across mammals. Then, we apply that model to distinguish instances where the genome sequence would predict conserved function versus a loss regulatory activity in that tissue. We present criteria for systematically evaluating model performance for this task and use them to demonstrate that our models accurately predict tissue-specific conservation and divergence in open chromatin between primate and rodent species, vastly out-performing leading nucleotide alignment-based approaches. We then apply our models to predict open chromatin at orthologs of brain and liver open chromatin regions across hundreds of mammals and find that brain enhancers associated with neuron activity and liver enhancers associated with liver regeneration have a stronger tendency than the general population to have predicted lineage-specific open chromatin.

**Conclusion:** The framework presented here provides a mechanism to annotate tissue-specific regulatory function across hundreds of genomes and to study enhancer evolution using predicted regulatory differences rather than nucleotide-level conservation measurements.

## BACKGROUND

The study of conservation has had a tremendous impact in multiple areas of mammalian biology. When a new genome is sequenced, conservation is applied to provide high-quality annotations of candidate exons, introns, promoters, and other likely functional genomic regions [1]. Regions of the human genome conserved across other primates or mammals show a stronger enrichment for disease-associated loci than any other evaluated category of regions [2]. Conversely, regions of the human genome that display accelerated evolution have been implicated in human-specific adaptation [3]. In endangered species, molecular conservation has been applied predict which regions of low heterozygosity may be impacting fitness [4]. Conservation has also been applied to find regions of the genome associated with the evolution of complex phenotype across mammals and vertebrates more broadly, including the loss of limb function [5], loss of eyesight [6], and longevity [7]. The powers of these studies are still growing, with many consortia, including the Vertebrate Genomes Project [8], the Genome 10K Project [9], the Bat 1K Project [10], and the Zoonomia Project [4], sequencing, assembling, and aligning [11] genomes from hundreds of mammals, including endangered species and species that live in remote parts of the world. Using these data, we can investigate conservation by comparing the DNA sequences of species whose most recent common ancestors lived tens of millions of years ago.

The methods used to infer conservation across species, including those applied to many of the challenges described above, typically rely on nucleotide-level constraint. They generally begin by aligning the nucleotides of multiple genomes together [12, 13]. Building off of those alignments, PhyloP calculates nucleotide-level constraint relative a neutral model of evolution, which can be aggregated across broader regions to identify signatures of conservation or acceleration [3, 13, 14]. To study the diversity of mammalian phenotypes, nucleotide alignments are often modeled in the context of a tree structure to look for signatures of positive or negative selection [15–17]. Once nucleotide-level selection has been inferred, additional techniques have been applied to link those patterns of selection to convergent evolution, instances where a specific phenotype has evolved independently in multiple lineages [6, 17, 18].

As new genomic resources have become available and computational techniques have advanced, it has become clear that a large component of phenotypic evolution is mediated by differences in *cis*-regulatory elements, many of which are enhancers that control gene expression [19–21]. Within the human population, enhancers show a strong enrichment for disease-associated genetic variants [22, 23]. Across species, nucleotide-level selection in enhancers has been associated the loss of eyesight, hindlimbs, and external testes [6, 18, 24, 25]. Numerous other complex phenotypes have been linked to gene expression differences across species, including vocal learning [26], domestication [27, 28], longevity [29–31], brain size [32], echolocation [33], and monogamy [34]. While nucleotide-level selection in enhancers is being applied to study the evolution of some of these phenotypes [35], recent studies of enhancers across and species suggest a model of evolution in which nucleotide-level conservation of enhancers can be low in spite of enhancers maintaining their function [36, 37].

Much of our knowledge of enhancers comes from regulatory genomics measurements that are associated with enhancer activity, especially the Assay for Transposase-Accessible Chromatin using Sequencing (ATAC-Seq) and the DNase hypersensitivity assay for open chromatin and chromatin immunoprecipitation sequencing (ChIP-Seq) for the histone modifications H3K27ac and H3K4me1 [38–41]. Studies involving these assays have demonstrated that enhancers, relative to genes, are substantially more tissue- or cell type-specific and generally less conserved across species [42–45]. Within a given cell type or tissue, a combinatorial code of transcription factor (TF) binding motifs and other sequence patterns determines the ultimate regulatory activity [46–48]. In a striking example of this, a recent study found that an Islet enhancer’s developmental function remains remarkably conserved across the 700 million years of evolution between mammals and sponges by maintaining a similar set of TF motifs despite negligible detectable conservation at the nucleotide level [36]. This understanding is further supported by studies of TF binding across species that display a large turnover in individual binding sites [49–51], even though gene expression is often highly conserved, and genes with highly conserved expression often have conservation in the TFs whose motifs occur within their candidate enhancers [52]. Therefore, to study conservation of enhancers, new methods are required that link genome sequence differences at candidate enhancers to differences in enhancer function.

Advances in the application of machine learning techniques to regulatory genomics open up the possibility that conservation could by assayed at the level of the regulatory code, rather that the alignment of individual nucleotides. In one recent study, support vector machines (SVMs) and convolutional neural networks (CNNs) were able to predict which 3kb windows have the enhancer-associated histone modification H3K27ac in brain, liver, and limb tissue of human, macaque, and mouse [53]. Importantly, the study found that models trained in one mammal achieved high accuracy in another mammal in the same clade and in another mammal in a different clade, suggesting that the regulatory code in all three of these tissues is highly conserved across mammals [54]. Another study obtained a similar result for regions associated with H3K27ac [55], and two other studies have obtained similar results using another proxy for enhancer activity – open chromatin regions (OCRs) [56, 57]. One of these studies found that training CNNs on OCRs from multiple mammals had better performance than training CNNs on OCRs from a single mammal, albeit using 131,072bp sequences as input. The boost in power from incorporating multiple species generalized to predicting TF binding strength from ChIP-seq data and gene expression from RNA-seq data [56]. An additional study found that a combined CNN-recurrent neural network [58, 59] trained on sequences underlying 500bp OCRs from melanoma cell lines in one species can accurately predict melanoma cell line open chromatin in other species at a wide range of genetic distances from the training species, including in parts of the genome with low sequence conservation between the training and evaluation species [57].

While these studies represent major advances in cross-species enhancer prediction, they have yet to comprehensively demonstrate an ability to identify sequence differences between species that are associated with differences in regulatory genomic measurements of enhancer activity, which is crucial for their application as a conservation metric (**Table 1**). In fact, an additional study trained SVMs to predict liver enhancers using dinucleotide-shuffled candidate enhancers as negatives. While the overall performance was good, human enhancers whose orthologs are active in Old World monkeys but not New World monkeys were predicted to have consistent activity across all primates, showing that models with good overall performance do not always work well on enhancer orthologs whose activity differs between species [60].

**Table 1:** Comparison of Evaluation Criteria Used to Evaluate Models in Candidate Enhancer Activity Conservation Prediction Papers.

| Evaluation Criterium | Chen *et al.*, 2018 | Huh *et al.*, 2018 | Kelley, 2020 | Minnoye *et al.*, 2020 | This Paper |
| --- | --- | --- | --- | --- | --- |
| Evaluation on Genomic Regions not Used in Training | ✓ | ✓ | ✓ | ✓ | ✓ |
| Evaluation on Species not Used in Training | ✓ | ✓ | ✓ | ✗ | ✓ |
| Comparison of Model Trained on One Species to Model Trained on Multiple Species | ✗ | ✓ | ✓ | ✗ | ✓ |
| Evaluation of Lineage-Specific Enhancer Accuracy | ✗ | ✗ | ✗ | ✗ | ✓ |
| Evaluation of Tissue-Specific Enhancer Accuracy | ✗ | ✗ | ✗ | ✗ | ✓ |
| Evaluation of Phylogeny-Matching Correlations | ✗ | ✗ | ✗ | ✗ | ✓ |
| Comparison to Conservation Scores | ✗ | ✗ | ✗ | ✗ | ✓ |
| Identification of Biologically Relevant Enhancers with Correctly Predicted Lineage-Specificity | ✗ | ✗ | ✗ | ✓ | ✓ |

To predict and evaluate candidate *cis*-regulatory element differences across species, we chose to focus on OCRs, which are only a proxy of regulatory activity, but whose high resolution has the potential to better identify specific genome sequence differences associated with putative regulatory activity. We leveraged new, controlled open chromatin experiments conducted by our laboratory [61] and trained a set of new models to predict OCR differences across species (**Figure 1**). We evaluated model performance using new criteria that we developed for this task, which focus on the ability to predict similarities and differences across species and tissues rather than the large number of regions that are consistently open or closed (**Supplemental Figure 1a**). We also developed a novel method for associating previously identified candidate enhancer sets with predicted lineage-specific open chromatin and lack of open chromatin, in which we clustered OCRs based on their predicted open chromatin and identified clusters overlapping candidate enhancer sets more than expected by chance. Our methods for evaluating approaches to OCR ortholog open chromatin status prediction and for identifying predicted lineage-specific OCRs associated with candidate enhancer sets can be applied to any tissue or cell type with open chromatin data from multiple species. This allows for reliable annotation of open chromatin across the hundreds of new genomes that are being sequenced, providing a valuable resource for research communities studying gene regulation in non-traditional model organisms, especially where direct measurements are not feasible. We anticipate that this work will encourage researchers to develop and properly evaluate new models for predicting OCR ortholog open chromatin across species and help reveal potential functional roles of lineage-specific enhancers, enabling us to uncover transcriptional regulatory mechanisms underlying the evolution of mammalian phenotypic diversity.

**Figure 1:**
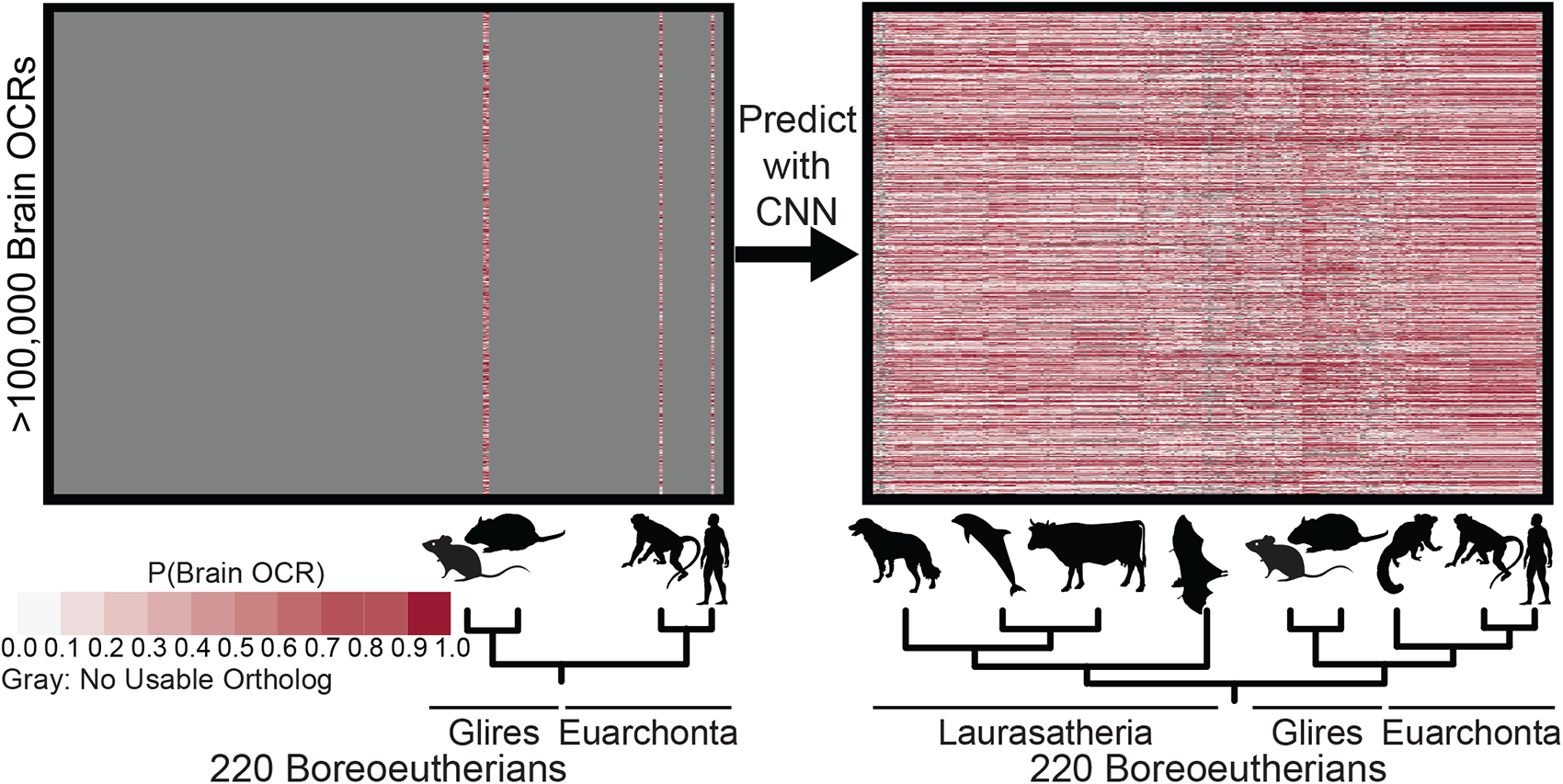
OCR Ortholog Open Chromatin Status Prediction Framework Overview. We trained a convolutional neural network (CNN) for predicting brain open chromatin using sequences underlying brain open chromatin region (OCR) orthologs in a small number of species and used the CNN to predict brain OCR ortholog open chromatin status across all of the species in the Zoonomia Project. We used the sequences underlying the orthologs for which we have brain open chromatin data to train a CNN for predicting open chromatin. Then, we used the CNN to predict the probability of brain open chromatin for all brain OCR orthologs; predictions are illustrated on the right.

## RESULTS

### Dataset Construction for Evaluating Approaches for OCR Ortholog Open Chromatin Prediction

To demonstrate the ability to predict OCR differences across species, we created a dataset of brain OCRs and their orthologs across dozens of species. To do this, we gathered open chromatin data generated by ATAC-seq [41, 62] or DNase hypersensitivity [63] from two brain regions – cortex and striatum – in four species: *Homo sapiens* [38, 64, 65], *Macaca Mulatta* [61], *Mus musculus* [66], and *Rattus norvegicus* [61]. We then defined our OCRs to be the 250bp in each direction of summits of non-exonic cortex open chromatin peaks that (1) overlap striatum open chromatin peaks, (2) are less than 1kb (to exclude super-enhancers), and (3) are at least 20kb from the nearest transcription start site (TSS) so that so they would not overlap promoters. We also identified the orthologs of each OCR in each of the other species collected by the Zoonomia Project [4, 67] (**Figure 1**). Rather than focus on longer regions, which have been shown to work well in some previous studies [54, 56], we used 500bp regions because of the ability shorter regions provide to focus on the impact of local sequence differences, because of the relative ease of obtaining orthologs of shorter regions in genomes with short scaffolds, and because predictions of enhancer activity for shorter regions are easier to experimentally validate.

We compared performance for two measures of conservation scores – PhastCons [13] and PhyloP [14] – and machine learning models trained using five different negative sets that are similar to those used for related tasks: (1) flanking regions [68], (2) OCRs from other tissues [56, 57], (3) about ten times as many G/C- and repeat-matched regions as positives [54], (4) about twice as many G/C- and repeat-matched regions as positives, and (5) ten dinucleotide-shuffled versions of each positive [69] (**Supplemental Figure 1b**). We additionally created a sixth, novel negative set to force the model to learn signatures of OCRs whose orthologs’ open chromatin status differs between species – sequences with closed chromatin in brain of a given species whose orthologs in another species are brain OCRs (we called this “non-OCR orthologs of OCRs,” which are the white regions in left part of **Figure 1**) – and included this negative set in our comparison (**Supplemental Figure 1b**). For a modeling approach, we chose CNNs [53, 70] because they can model complex combinatorial relationships between sequence patterns, and changes in a single TF motif often do not cause changes in open chromatin [48]; because they do not require an explicit featurization of the data, and many sequence patterns involved in brain open chromatin remain to be discovered; and because they can make predictions quickly relative to SVMs, the other leading approach for related tasks [54]. We did the comparison using conservation scores generated using mouse reference-based alignments [12, 71] and models trained on only mouse sequences so that we could evaluate their performance on both closely and distantly related species not used for training the machine learning models [54].

### Optimizing and Evaluating Methods for Achieving Lineage-Specific OCR Accuracy

We evaluated the machine learning model trained on each negative set on its corresponding test set. We found that all models worked well, with every model achieving an AUC > 0.85 and an AUPRC > 0.7 (**Supplemental Figure 1c**). This performance is especially impressive given that the ratios of the number of negatives to the number of positives ranged from approximately 1.2:1 (non-OCR orthologs of OCRs) to approximately 20:1 (OCRs in other tissues). The best-performing model was the model with the negative set consisting of dinucleotide-shuffled brain OCRs (**Supplemental Figure 1c**). However, in this comparison, each model was evaluated on a different negative set, so this evaluation may not be indicative of how useful each model would be in answering questions about gene expression evolution.

We therefore also evaluated each model’s and the conservation scores’ lineage-specific OCR accuracy (**Supplemental Figure 1a**), which we did by evaluating models and computing conservation scores for the OCR orthologs whose brain open chromatin status differs between species, an evaluation not done in any previous studies (**Table 1**). We did this by evaluating whether a model trained in one species could make accurate predictions of brain open chromatin status on sequences obtained from another species [54]. However, unlike other previous work that evaluated overall performance [54], we evaluated performance on the subset of regions in another species whose brain open chromatin status differs from that of their orthologs in the training species, as this is necessary for showing that models can accurately predict conservation – differences and similarities in brain open chromatin status between species. We therefore evaluated our models trained in mouse on macaque brain OCRs whose mouse orthologs are closed and on macaque brain closed chromatin regions whose mouse orthologs are open in brain and compared our results to conservation scores. We found that conservation scores tended to be slightly lower for both of these sets of regions than they for OCRs whose orthologs in both mouse and macaque are open, but they could not clearly distinguish between whether an OCR’s ortholog was open in mouse or in macaque (**Figure 2a**). On the other hand, all models achieved decent performance (AUC > 0.65, AUPRC > 0.55) (**Supplemental Figure 1g**), though the model trained on the dinucleotide-shuffled brain OCR negatives predicted that almost half of the macaque non-OCRs whose mouse orthologs are OCRs are open in brain (**Figure 2a**), causing it to have the worst performance (**Supplemental Figure 1g**). We also evaluated our models on human (**Supplemental Figure 1h, Supplemental Figure 2**) and rat (**Supplemental Figure 1i, Supplemental Figure 3a**) regions with differing brain open chromatin statuses from their mouse orthologs and found that the models generally did not work as well for such regions but still obtained decent performance and that the relative performance of conservation scores and different negative sets was similar to what it was for the macaque regions. We did additional evaluations to further show that overall model performance is not indicative of lineage-specific OCR accuracy, that re-calibrating the models with the positive training set and the training set from our novel negative set did not substantially change the relative performances of the models trained on different negative sets, and that conservation scores are significantly inferior to our best machine learning models at predicting open chromatin conservation across lineages (**Supplemental Notes, Supplemental Tables 1-9, Supplemental Figure 4**). These results demonstrate the necessity of evaluating models on OCRs whose orthologs’ open chromatin statuses differ between species.

**Figure 2:**
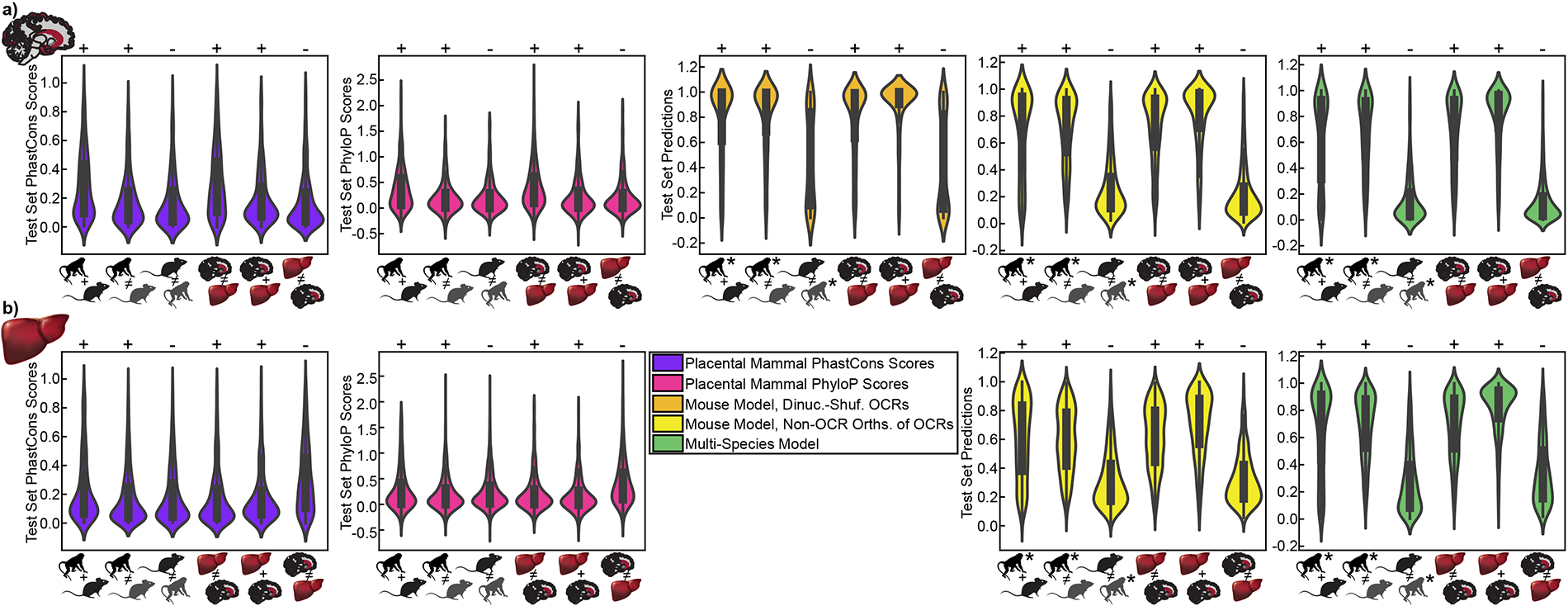
Violin Plots for Lineage-Specific and Tissue-Specific OCR Accuracy Evaluation in Macaque. **a)** Comparison of PhastCons [13] and PhyloP [14] scores to three different machine learning models’ predictions for brain OCRs with conserved open chromatin across mouse and macaque, macaque brain OCRs whose mouse orthologs are closed in brain, macaque brain non-OCRs whose mouse orthologs are open in brain, macaque brain OCRs that are closed in liver, macaque brain OCRs that are open in liver (centered on brain peak summits), and macaque liver OCRs that are closed in brain. **b)** Comparison of PhastCons [13] and PhyloP [14] scores to two different machine learning models’ predictions for liver OCRs with conserved open chromatin across mouse and macaque, macaque liver OCRs whose mouse orthologs are closed in liver, macaque liver non-OCRs whose mouse orthologs are open in liver, macaque liver OCRs that are closed in brain, macaque liver OCRs that are open in brain (centered on liver peak summits), and macaque brain OCRs that are closed in liver. +’s indicate that values should be large, and -’s indicate that values should be small. Conservation scores were generated from the mm10-based placental mammals alignment [12, 71] and averaged over 500bp centered on peak summits, where mouse peak summits were used for OCRs conserved between mouse and macaque and OCRs in mouse whose macaque orthologs are closed, and mouse orthologs of macaque peak summits were used for other evaluations. All machine learning model predictions were made using macaque sequences, where the macaque sequences for OCRs conserved between mouse and macaque and OCRs in mouse whose macaque orthologs are closed were centered on macaque orthologs of mouse peak summits, and macaque peak summits were used for other evaluations. Animal silhouettes were obtained from PhyloPic [132]. *’s indicate the species from which sequences were obtained for making predictions. Dinuc.-shuf. stands for dinucleotide-shuffled, and orths. stands for orthologs.

### Best Overall Model Performance Does Not Guarantee Tissue-Specific OCR Accuracy

In addition to evaluating conservation score and machine learning model performance on OCR orthologs whose open chromatin status differs between species, we also determined if conservation scores and our machine learning models achieved high tissue-specific OCR accuracy (**Supplemental Figure 1a**). While previous studies evaluated model performance in different tissues or cell lines [54, 56, 57] and showed that models trained in the evaluation tissue tend to be more accurate than models trained in other tissues [54], no study has directly evaluated model performance on tissue-specific candidate enhancers or candidate non-enhancers (**Table 1**). To determine if our models learned sequence patterns associated with only brain-specific open chromatin, we evaluated the conservation scores and our models’ predictions for the subset of brain OCRs that do not overlap liver OCRs and the subset of brain OCRs that do overlap liver OCRs. Test set predictions from all models for both of these subsets of the positive set were usually close to one (**Figure 2a, Supplemental Figures 2-3, Supplemental Figure 5a**). To determine if our models learned only sequence patterns that are indicative of general open chromatin, we also evaluated our models’ predictions for the liver OCRs that do not overlap brain OCRs. We compared this to the predictions on the negative set. We found that predictions on both of these sets tended to be close to zero. However, the liver, non-brain open chromatin status predictions from the model trained with dinucleotide-shuffled OCR negatives tended to be more evenly distributed between zero and one than the liver, non-brain open chromatin status predictions for the models trained with the other negative sets (**Figure 2a, Supplemental Figures 2-3, Supplemental Figure 5a**). In addition, unlike the machine learning model predictions, the conservation scores for brain, non-liver OCRs; OCRs in brain and liver; and liver, non-brain OCRs are similar to each other (**Figure 2a, Supplemental Figures 2-3**), demonstrating that conservation scores, in contrast to most of our machine learning models, provide little information about the tissue-specificity of open chromatin. We also did a comparison of the performances of models trained on different negative sets in which we limited the positive set to brain OCRs that overlap liver OCRs and defined the negative set as liver OCRs that do not overlap brain OCRs.

We found that all models worked well (AUC > 0.75, AUPRC > 0.6) on mouse as well as on macaque and rat, which were not used in training, with the model trained on dinucleotide-shuffled brain OCR negatives providing the worst performance (**Supplemental Figure 5b**). The difference in performance may be a result of differences between the TF motifs learned by the models (**Supplemental Notes, Supplemental Figure 6**). In addition, we found that the models that tended to work better on positives were more accurate for shared brain and liver OCRs than for liver, non-brain OCRs and that the models that tended to work better on brain closed chromatin orthologs of brain OCRs were more accurate for liver, non-brain OCRs than for shared brain and liver OCRs (**Supplemental Tables 10-11, Supplemental Figure 5c**). Calibration with the positive training set and the training set from our novel negative set had a similar effect to calibration for clade-specific open and closed chromatin regions (**Supplemental Tables 10-12, Supplemental Figure 5c**). These results show that conservation scores cannot be used directly to understand the tissue-specificity of open chromatin and that models with good overall performance are not always able to comprehensively learn open chromatin sequence patterns that are specific to the tissue in which they were trained.

### Predictions from Models of OCR Ortholog Open Chromatin Status Have Phylogeny-Matching Correlations

We also determined whether our models’ predictions have phylogeny-matching correlations in a way that does not require open chromatin data from multiple species. To do this, we obtained the orthologs of the mouse brain OCRs in all of the fifty-six Glires species in the Zoonomia Project [4, 67], used our machine learning models to predict the brain open chromatin statuses of these orthologs, computed the mean brain open chromatin statuses across all brain OCR orthologs in each species, and computed the correlation between mean predicted brain open chromatin status and evolutionary distance from mouse. Although there are OCR orthologs, such as species-specific OCRs and OCRs with convergently evolved open chromatin [6], whose open chromatin conservation across species is not associated with phylogenetic distance, we think that such OCRs in most tissues are a minority due to the principle of evolutionary parsimony and a previous study of enhancer activity across multiple species [44]. As anticipated, all models showed a strong negative correlation between mean predicted brain open chromatin status and divergence from mouse (**Supplemental Figure 7a**). Nevertheless, there is still more open chromatin at these brain OCR orthologs than would be expected from brain non-OCRs, even in the most distantly related Glires species, because all mean predictions are greater than the mean predictions for the negative test sets (**Supplemental Figure 7a**). We also expected there to be a strong positive correlation between the standard deviation of open chromatin status and divergence from mouse because most brain OCR orthologs in species closely related to mouse are active in brain, while the brain open chromatin status of brain OCR orthologs in species that are more distantly related should vary. We found this expected positive correlation for the predictions from machine learning models trained on all negative sets (**Supplemental Figure 7b**).

### Machine Learning Models Trained Using Data from Multiple Species Can Accurately Predict OCR Orthologs’ Open Chromatin Statuses

Based on other research showing that training models with data from multiple species can improve OCR prediction accuracy [56], we trained additional machine learning models using open chromatin data from multiple species. For each of brain and liver, we used the open chromatin data from all of the species that we had collected as positives (four species for brain, three species for liver) and the orthologs of all these OCRs in the other species for which we had data that did not overlap brain or liver open chromatin, respectively, as negatives. Before training multi-species models, we trained mouse-only models for liver and found that they achieved good performance for all of our criteria (**Figure 2b, Supplemental Notes, Supplemental Figure 8, Supplemental Table 13**). We then trained brain and liver multi-species models and found that they achieved high lineage-specific and tissue-specific OCR accuracy, where performance was generally better than the performance for any of the models trained on only mouse sequences (**Figure 2**, **Figures 3a-b**). In addition, we determined if the multi-species brain and liver models’ predictions had phylogeny-matching correlations by using them to predict the OCR ortholog open chromatin status of mouse brain and liver OCRs, respectively, across Glires and found strong negative correlations between divergence from mouse and mean OCR ortholog open chromatin status predictions (**Figures 3c-d**). We also found strong positive correlations between divergence from mouse and standard deviations of OCR ortholog open chromatin status predictions (**Supplemental Figures 9a-b**). For both these models and our mouse-only models, our phylogeny-matching correlation results could not be fully explained by genome quality (**Supplemental Notes, Supplemental Figure 10, Supplemental Tables 14-15**). Finally, we evaluated our mouse-only and multi-species liver models on Laurasiatheria-specific liver OCRs and liver non-OCRs [72], as no Laurasiatheria were used in training either model, and found that the multi-species liver model had better performance than the mouse-only liver model (**Figure 3e**). Like our mouse-only models, our multi-species brain and liver models also learned TF motifs for TFs that are known to be involved in brain and liver, respectively (**Supplemental Notes, Supplemental Figures 9c-d**), and their predictions were significantly more associated with open chromatin conservation than conservation scores were (**Figure 2**, **Supplemental Notes, Supplemental Table 16**).

**Figure 3:**
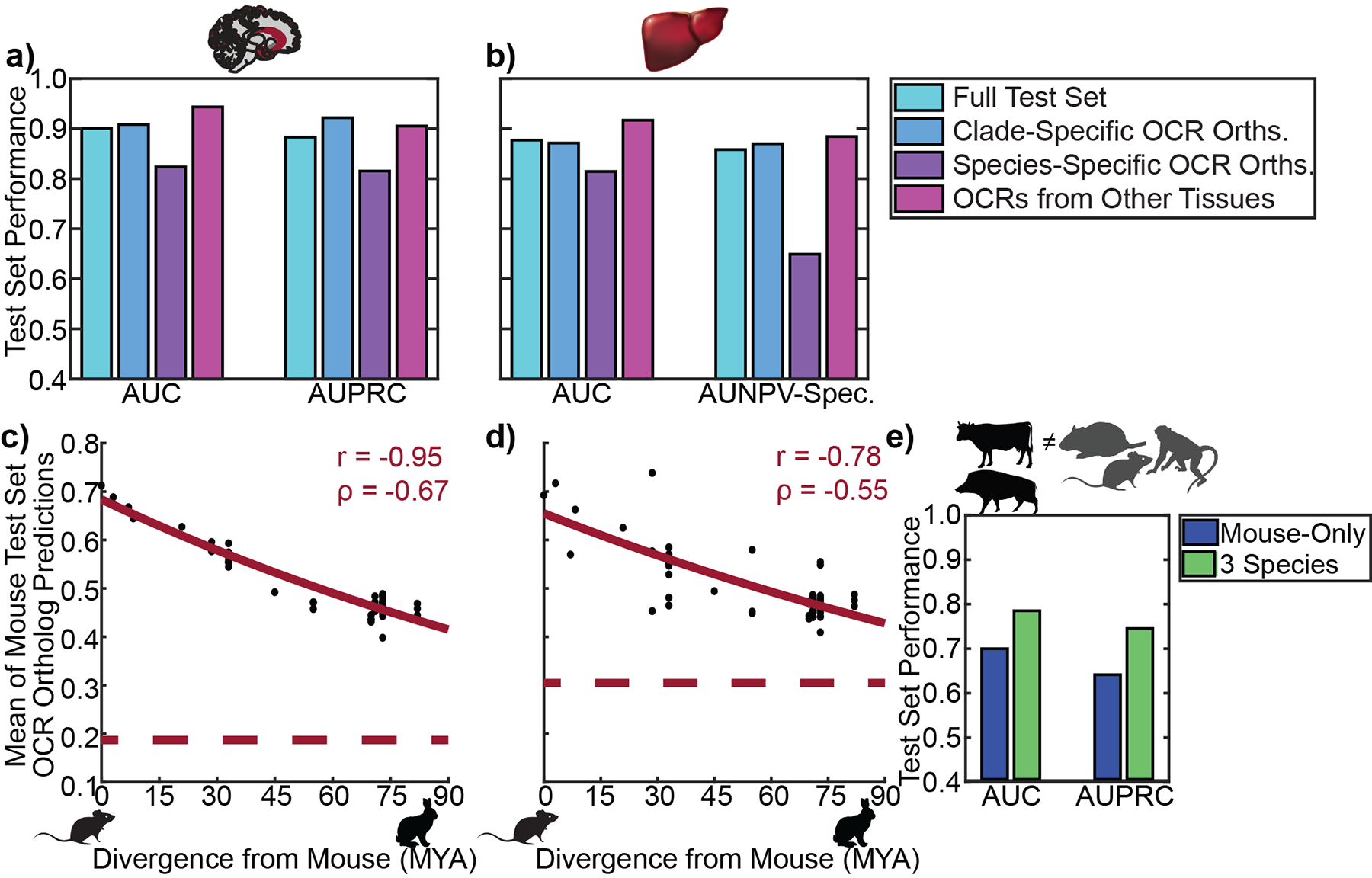
Multi-Species Model Performance. **a)** Performance of multi-species brain model on full test set, subset of test set consisting of clade-specific brain OCRs and non-OCRs, subset of test set consisting of species-specific brain OCRs and non-OCRs, and subset of positive test set consisting of OCRs shared between brain and liver as well as subset of liver OCRs from test set genomic regions that do not overlap brain OCRs. **b)** Performance of multi-species liver model on full test set, subset of test set consisting of clade-specific liver OCRs and non-OCRs, subset of test set consisting of species-specific liver OCRs and non-OCRs, and subset of positive test set consisting of OCRs shared between liver and brain as well as subset of brain OCRs from test set genomic regions that do not overlap liver OCRs. We reported area under the negative predictive value (NPV)-specificity (Spec.) curve instead of the AUPRC because these test sets have more positives than negatives. **c)** Divergence from mouse versus mean multi-species brain model predictions across mouse brain OCR orthologs in Glires. **d)** Divergence from mouse versus mean multi-species liver model predictions across mouse liver OCR orthologs in Glires. **e)** Performance of mouse-only liver model versus multi-species liver model on Laurasiatheria-specific liver enhancers and liver non-enhancers whose mouse orthologs are in the test set. Animal silhouettes were obtained from PhyloPic [132]. AUC stands for area under the receiver operating characteristic curve, AUPRC stands for area under the precision-recall curve, and MYA stands for millions of years ago. The red curves are the best fit exponential function of the form y = ae^bx^. The red dotted lines are the average prediction across test set negatives.

Some of the OCR orthologs for which our multi-species brain model correctly predicted brain open chromatin conservation in spite of low mean sequence conservation or for which our models correctly predicted lack of brain open chromatin conservation in spite of high mean sequence conservation are near genes that have been shown to play important roles in the brain. For example, there is a region on mouse chromosome 2 – part of our test set – that has low mean sequence conservation according to PhastCons [13] and PhyloP [14] but high brain experimentally identified and predicted open chromatin conservation between mouse and macaque (**Figure 4a**) and whose mouse and macaque orthologs are located near the gene *Stx16*. This gene is involved in vesicle trafficking in most tissues, including the brain [73], and may play a role in Alzheimer’s disease [74]; in fact, its role in axon regeneration is conserved between mammals and *C. elegans* [75]. Although this region near *Stx16* has generally low sequence conservation, running TomTom [76] on the 22bp sequence with high conservation revealed a subsequence that is similar to the *Fos* motif, which is also found in the macaque ortholog. Since our machine learning model used sequence similarity to the *Fos* motif in making predictions (**Supplemental Figure 9c**), the machine learning model was likely able to automatically determine that it should use this sequence in making its prediction. In addition, there is a region on mouse chromosome 2 that has high mean sequence conservation but low experimentally identified and predicted open chromatin conservation between mouse and macaque (**Figure 4b**) and whose mouse and macaque orthologs are located near the gene *Lnpk*. This gene is an endoplasmic reticulum junction stabilizer [77] that has been shown to play a role in brain and limb development [78], and mutations in this gene have been associated with neurodevelopmental disorders [79]. It is possible that this region of the genome near *Lnpk* has a high degree of sequence conservation because it has functions in other tissues. These results demonstrate the potential benefits of using OCR ortholog open chromatin status predictions instead of mean conservation scores for studying the evolution of the expression of important genes in a tissue of interest.

**Figure 4:**
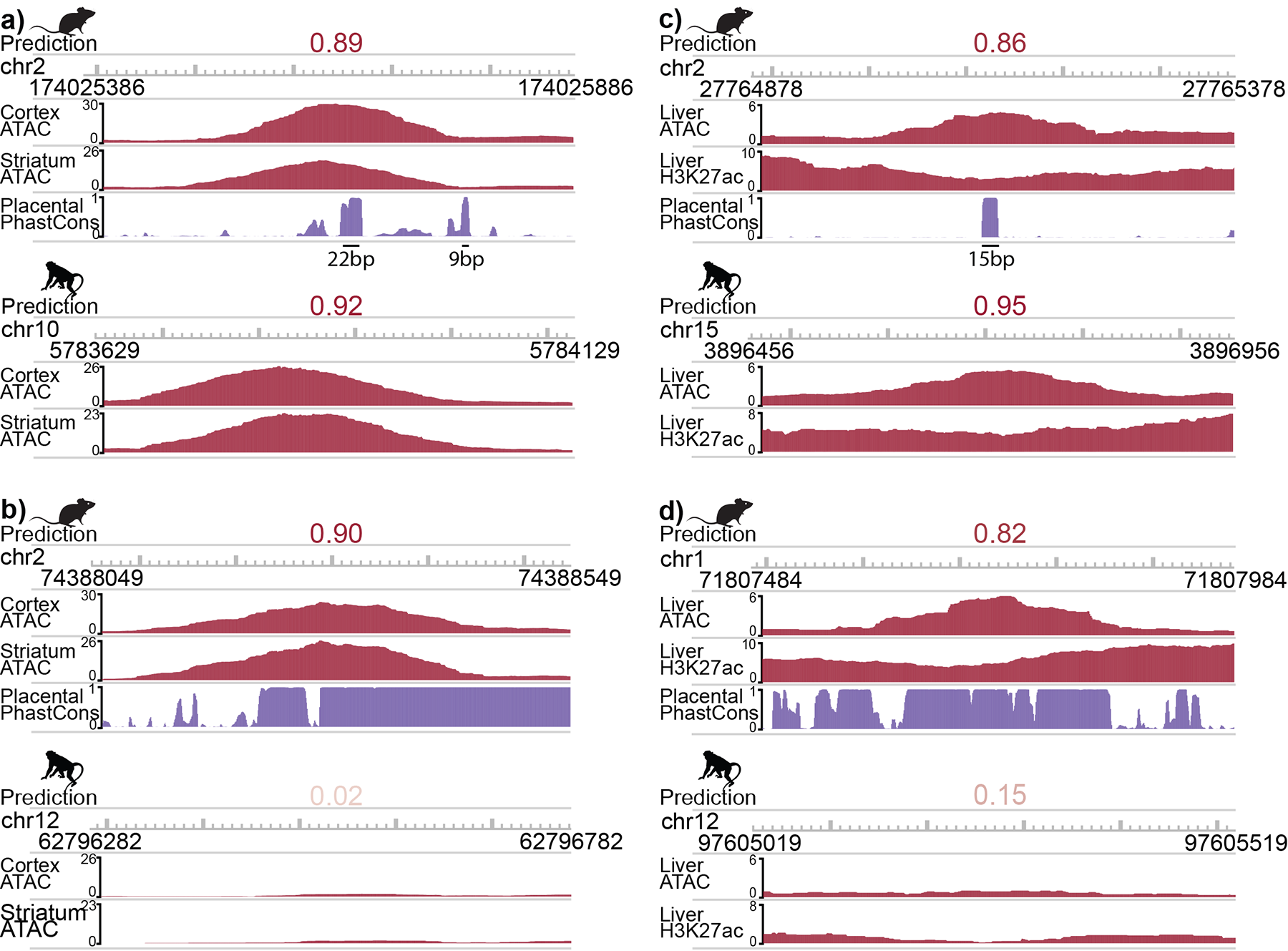
Examples of Mean Conservation Score and Open Chromatin Status Prediction versus Open Chromatin Conservation. **a)** 7-week-old mouse cortex and striatum and macaque orofacial motor cortex (“Cortex”) and putamen (“Striatum”) open chromatin signal for a mouse brain OCR that is 50,328 bp away from the *Stx16* transcription start site (TSS). Experimentally identified and predicted brain open chromatin statuses are conserved even though mean mouse PhastCons score is low. **b)** 7-week-old mouse cortex and striatum and macaque orofacial motor cortex (“Cortex”) and putamen (“Striatum”) open chromatin signal for a mouse brain OCR that is 144,474 bp away from the *Lnpk* TSS. Experimentally identified and predicted brain open chromatin statuses are not conserved even though mean mouse PhastCons score is high. **c)** Our mouse liver and macaque liver open chromatin signal for a mouse liver OCR that is 24,814 bp away from the *Rxra* TSS. Experimentally identified and predicted liver open chromatin statuses are conserved even though mean mouse PhastCons score is low. **d)** Our mouse liver and macaque liver open chromatin signal for a mouse liver OCR that is 154,404 bp away from the *Fn1* TSS. Experimentally identified and predicted liver open chromatin statuses are not conserved even though mean mouse PhastCons score is high. Animal silhouettes were obtained from PhyloPic [132]. Regions are mouse cortex open chromatin peak summits +/-250bp and their macaque orthologs, signals are from pooled reads across biological replicates, and liver H3K27ac ChIP-seq data comes from E-MTAB-2633 [37].

In addition, some of the accurate multi-species model liver open chromatin conservation predictions disagreed with the mean conservation scores. For instance, there is a region on mouse chromosome 2 that has high experimentally identified and predicted liver open chromatin conservation but low sequence conservation (**Figure 4c**) and whose mouse and macaque orthologs are located near *Rxra*. This gene is a TF involved in regulating lipid metabolism [80–82], TF-MoDISco identified a sequence similar to its motif as being important in our liver models (**Supplemental Figure 8f, Supplemental Figure 9d**), and its liver expression is stable across fifteen mammals [83]. Although this region near *Rxra* has generally low sequence conservation, the 15bp segment with high conservation is similar to the motif for *Ctcf* according to TomTom [76], and that motif is also found in the macaque ortholog. Since our machine learning model used sequence similarity to the *Ctcf* motif in making predictions (**Supplemental Figure 9d**), the machine learning model was likely able to automatically determine that it should use this sequence in making its prediction. There is also a region on mouse chromosome 1, which is part of our test set, whose mouse ortholog is an OCR and macaque ortholog is not an OCR according to our data and predictions in spite of being highly conserved (**Figure 4d**) and whose mouse and macaque orthologs are near *Fn1*. This gene has been implicated in liver fibrosis [84–86], and a multi-species liver RNA-seq study found that it has higher expression in mouse liver relative to livers of other mammals and birds and lower expression in primate livers relative to livers of other mammals and birds [87]. For both of these OCRs, the H3K27ac signal conservation in the same regions [37] is similar to the open chromatin status conservation, suggesting that the open chromatin status conservation is indicative of enhancer activity conservation (**Figures 4c-d**). These results suggest that using predicted OCR ortholog open chromatin status instead of conservation has the potential to be beneficial for understanding gene expression evolution in multiple tissues.

### Lineage-Specific Brain and Liver OCRs Are Associated with Neuron and Liver Functions

Since our models can accurately predict lineage-specific OCRs, we evaluated whether lineage-specific brain and liver OCRs were associated with brain and liver functions. First, we used publicly available RNA-seq data from the livers of thirteen placental mammals with high-quality genomes [83] to identify genes with Rodent-specific expression; we obtained 15 genes with false discovery rate (FDR) < 0.05 [88, 89]. We then used our predictions to identify liver OCRs with predicted Rodent-specific activity. One of the genes with Rodent-specific expression that is near OCRs with predicted liver-specific open chromatin is Atp9a (**Supplemental Figure 11**). This gene has high expression in mouse liver according to Mouse Genome Informatics [90], it has little or no RNA or protein expression in human liver according to all of the studies referenced in the Human Protein Atlas [91, 92], and its protein is expressed in the rat canicular membrane [93], a part of hepatocytes involved in bile production. This suggests that OCRs with predicted lineage-specific open chromatin may be regulating genes with lineage-specific expression.

To further investigate the roles of predicted lineage-specific OCRs, we clustered each of the brain and liver OCRs, where the features were the predicted activity in each of the Boreoeutheria from the Zoonomia Project [4]. We first used k-means clustering to cluster the OCRs into thousands of small clusters and then used affinity propagation cluster to cluster the smaller clusters into larger clusters. We selected affinity propagation clustering because we did not know how many clusters to expect, and affinity propagation clustering automatically determines the number of clusters. For each of brain and liver, we obtained a little over one hundred clusters with different patterns of predicted open chromatin across species (brain clusters: https://github.com/pfenninglab/OCROrthologPrediction/clusters/brain, liver clusters: https://github.com/pfenninglab/OCROrthologPrediction/liver).

We then determined whether each brain cluster that was open in mouse overlapped mouse candidate enhancers associated with neuron firing [94] more than expected by chance (**Supplemental Table 17**) and each brain cluster that was open in human overlapped human candidate enhancers associated with neuron activity [95] more than expected by chance (**Supplemental Table 18**). Interestingly, the candidate enhancers from each of these sets intersected clusters with predicted lineage-specific open chromatin or predicted lineage-specific lack of open chromatin more than expected by chance. Specifically, mouse neuron bicuculline (Bic)-specific candidate enhancers, where Bic induces neuron firing, were enriched for overlapping two predicted Murinae-specific brain open chromatin clusters – cluster 43 (**Figure 5**) and cluster 27 (**Supplemental Figure 12a**). We think that these results are unlikely to be explained by the number of usable orthologs or conservation because mouse brain OCRs overlapping Bic-specific candidate enhancers do not have significantly fewer usable orthologs or lower conservation according to PhastCons [13] or PhyloP [14] than mouse brain OCRs in general. In contrast, mouse activity-invariant candidate enhancers were enriched for overlapping the predicted open chromatin in all species cluster (cluster 1), a noisy cluster without a clear pattern of predicted open chromatin (cluster 37), a noisy predicted Yangochioptera-specific brain non-open chromatin cluster (cluster 81), and a noisy predicted Primate-specific brain non-open chromatin cluster (cluster 88). Likewise, human candidate enhancers from GABAergic neurons made from hiPSCs from four genotypes that had increased activity two hours after exposure to potassium chloride (KCl), where KCl induces neuron activity, were enriched for overlapping a predicted Carnivora-, Perissodactyla- and Euarchonta- specific brain open chromatin cluster (cluster 74, **Figure 5**), a predicted Hystricognathi-specific brain non-open chromatin cluster (cluster 11, **Supplemental Figure 12b**), and a predicted Muroidea- and Pecora-specific brain non-open chromatin (cluster 48, **Supplemental Figure 12b**). These results are unlikely to be explained the number of usable orthologs or conservation because human brain OCRs overlapping this set of GABAergic neuron candidate enhancers tended to have more usable orthologs and higher conservation than human brain OCRs overall according to both PhastCons [13] and PhyloP [13]. In contrast, candidate enhancers from the same source that had decreased activity two hours after exposure to KCl were enriched for overlapping the predicted open chromatin in all species cluster (cluster 1) and a predicted Ruminantia-specific brain non-open chromatin cluster (cluster 82). These results suggest that there may be a relationship between enhancer response to neuron activity and whether the enhancer’s activity tends to be specific to the lineage in which it was identified.

**Figure 5:**
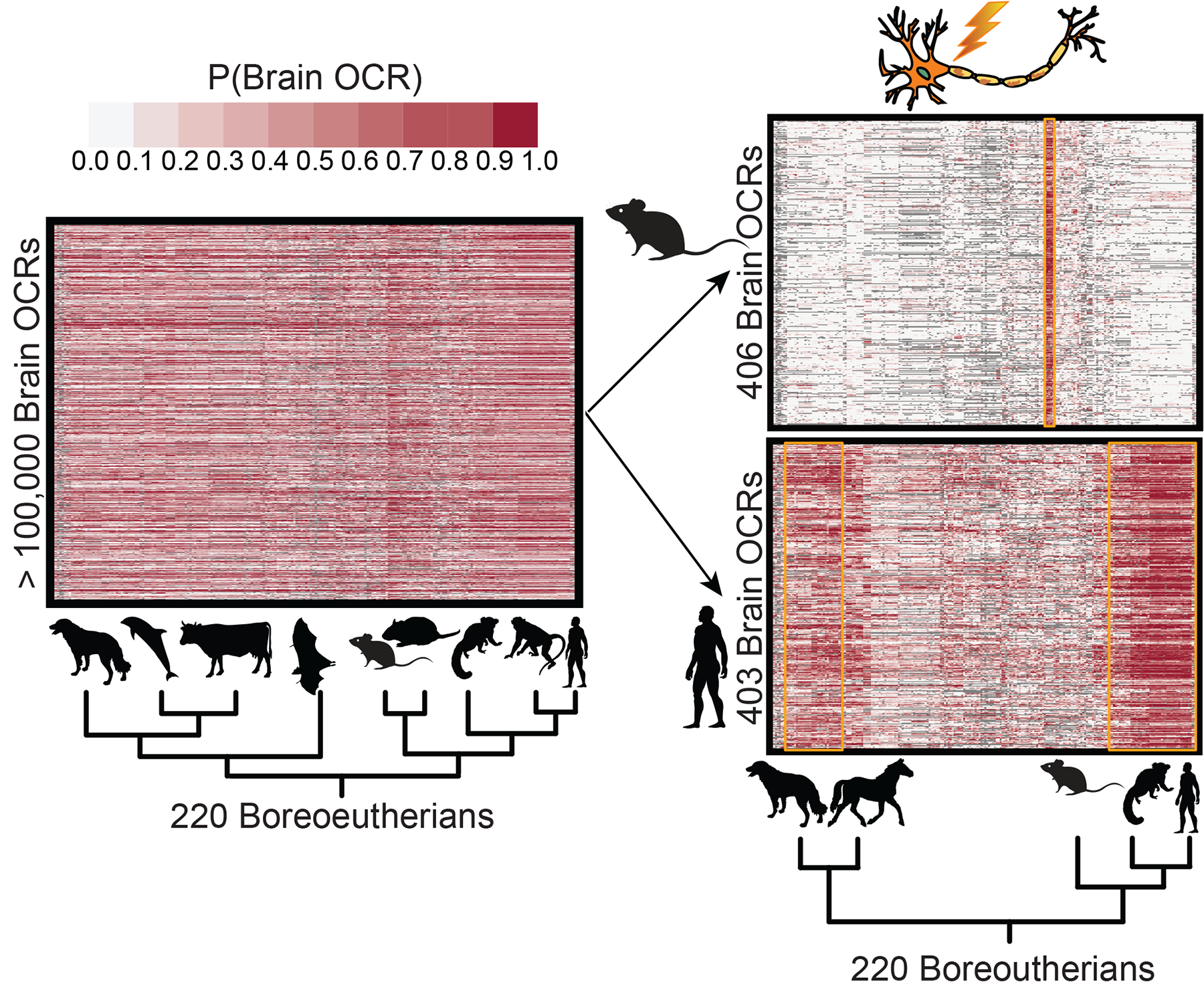
Examples of Brain OCR Clusters with Predicted Lineage-Specific Open Chromatin Associated with Neuron Activity. We clustered the brain OCRs, where the features were the brain predictions in each Boreoeutheria species from Zoonomia, and then identified clusters whose regions had significant overlap with regions associated with mouse neuron firing and human neuron activity. Mouse neuron firing enhancers had significant overlap with a predicted Murinae-specific OCR cluster (cluster 43), and human neuron activity enhancers had significant overlap with a predicted predicted Euarchonta and Carnivora-specific non-OCR cluster (cluster 74). Animal silhouettes were obtained from PhyloPic [132].

We also investigated whether each liver cluster that was active in mouse overlapped mouse candidate enhancers associated with liver regeneration [96] more than expected by chance (**Supplemental Table 19**). We found that candidate liver enhancers that have increased activity four weeks after hepatocyte repopulation relative to the control were enriched for overlapping a cluster with predicted Murinae-specific open chromatin – cluster 29 – as well as two clusters with predicted Muroidea-specific open chromatin – cluster 36 and cluster 100 (**Supplemental Figure 12c**). We think that these results are unlikely to be explained by the number of usable orthologs or conservation because OCRs overlapping this candidate enhancer set do not have significantly fewer usable orthologs or lower conservation according to PhastCons [13] or PhyloP [14] than liver OCRs overall. We also found that candidate liver enhancers that have increased activity one week after hepatocyte repopulation relative to the control were enriched for overlapping a cluster with noisy predicted Primate-specific liver non-open chromatin (cluster 83). In contrast to these findings, liver candidate enhancers with decreased activity four weeks after hepatocyte repopulation relative to the control were not enriched for overlapping any clusters, and liver candidate enhancers with decreased activity four weeks after hepatocyte repopulation relative to one week after hepatocyte repopulation were enriched for overlapping a cluster without a clear pattern of predicted open chromatin (cluster 39). These results suggest that there may be a relationship between liver regeneration in Murinae and Murinae-specific open chromatin.

## DISCUSSION

### Developing a Tissue-Specific Metric of Regulatory Conservation

While there are well-established methods for quantifying nucleotide-level conservation based on a sequence alignment [13, 14, 97], these methods have limited utility for the many enhancers with tissue-specific activity [42, 43] or low sequence conservation in spite of high functional conservation [36]. Yet quantifying enhancers’ conservation can provide insight into their functional relevance, such as the identification of convergent evolution in gene regulation underlying complex traits [20, 26, 29]. Here, we developed a machine learning-based approach to measure tissue-specific enhancer conservation. First, we trained CNNs to predict OCR orthologs’ open chromatin statuses in the tissue from which the OCRs were obtained. Then, we applied our models to predict the conservation of over 100,000 OCRs from each of brain and liver in over 200 mammals.

Our approach vastly outperformed nucleotide alignment-based methods of conservation PhastCons [13] and PhyloP [14] at predicting lineage- and tissue-specific open chromatin status. As expected, we found many examples where nucleotide-level conservation is low but the predicted open chromatin in a tissue of interest is conserved [36]. Conversely, we also identified cases where the nucleotide-level conservation is high, but the few differences between species disrupt open chromatin in a tissue of interest. We attribute the success of our method to our CNNs’ abilities to learn a conserved regulatory code linking genome sequence to tissue-specific open chromatin [54].

Our method builds upon recently published approaches that constructed machine learning models that learned conserved tissue-specific regulatory codes across species [54–57]. Yet a model’s ability to predict regulatory genomics features across multiple species does not guarantee that the model can accurately predict conservation or divergence. For example, a study predicting liver open chromatin using SVMs showed overall high accuracy but was not able to accurately predict cases where sequence differences between primates were associated with regulatory differences [60]. To demonstrate our ability to predict regulatory conservation, we developed a new, systematic set of evaluation criteria that evaluates if lineage- and tissue-specific differences can be predicted (**Table 1**). In the future, our criteria could be used to evaluate a wide range of extensions to our approach (**Supplemental Notes**).

### Limitations of Our Approach

Although our models excel in many criteria for their stated purpose of predicting lineage- and tissue-specific open chromatin, our models’ set-up also has inherent limitations that prevent them from fully fulfilling their purposes. For example, many OCRs’ open chromatin statuses are influenced by factors beyond the 250 base pairs in each direction of their summits, and the fixed-size inputs required by CNNs prevents us from modeling some of the long-range interactions that may influence open chromatin [98] (**Supplemental Notes**). In addition, since we predict open chromatin status of only OCR orthologs, we cannot identify OCRs whose orthologs are not OCRs in any species for which we have open chromatin data (**Supplemental Notes**). We also treat open chromatin status as binary, where, in reality, in bulk open chromatin data, open chromatin is a continuous signal (**Supplemental Notes**). Finally, our approach requires high-quality open chromatin from at least two species and is likely to have stronger performance when more are included. Cases where the datasets are not matched for sex, age, time of day, or other factors could influence model performance (**Supplemental Notes**).

### Applications of Inferring Conservation of Tissue-Specific Regulatory Activity

In addition to being accurate, our models’ predictions revealed that many OCRs may have lineage-specific patterns of open or closed chromatin and that these patterns may be functional. We identified a predicted Rodent-specific liver OCR near Atp9a, a gene with Rodent-specific expression in the liver that may play a role in bile production in rats [93]. In addition, we developed a novel approach to deciphering the putative lineage-specificity of candidate enhancers identified in other studies in which we clustered our OCRs using predicted activity as features and identified clusters with more overlap with these candidate enhancers than expected by chance. Using this approach, we found that mouse candidate enhancers activated during neuron firing [94] are enriched for overlapping clusters with predicted Murinae-specific open chromatin, and human candidate enhancers activated during neuron activity [95] are enriched for overlapping clusters with predicted Euarchonta-specific gains or Muroidea-specific losses of open chromatin, suggesting that there may be Muroidea-specific enhancer signatures of Muroidea neuron activity. Although no study, to our knowledge, has evaluated this for large numbers of species, a few studies have compared neuron activity between mice and one or two Euarchonta and found striking differences, including lower spiking frequencies in fast-spiking mouse cortical neurons relative to human and rhesus macaque [99] and lack of expression of h-Channels in mouse excitatory neurons relative to human [100]. We also found that candidate enhancers activated during liver regeneration [96] are enriched for overlapping predicted Murinae-specific open chromatin. We found clusters with predicted lineage-specific open chromatin for additional linages, predicted lineage-specific losses in open chromatin for additional lineages, and predicted convergent gains and losses in open chromatin; investigating the roles of the open chromatin regions in these clusters could be an exciting path towards identifying mechanisms underlying linage-specific differences between mammalian brains and livers.

Our approach has the potential to be applied to numerous other groups of species, cell types, and tissues. It does not require experimentally determined OCR data from more than a few species; it requires only genomes from a large number of species, which are being generated in unprecedented quantities [4, 9, 10]. In addition to having many advantages relative to nucleotide-level conservation scores, our use of 500bp sequences for training our OCR ortholog open chromatin status prediction models, as compared to other machine learning approaches that require sequences of 3kb [54] or over 100kb [56], enables us to make predictions in species whose assemblies have short scaffolds, including many of the species in the Zoonomia [4] and B10K datasets [101].

Our open chromatin conservation predictions can also be used in a forward genomics approach [102] to help identify the mechanisms underlying the evolution of the expression of genes of interest or of phenotypes that have evolved through gene expression. This can be done by identifying OCR orthologs whose changes in predicted open chromatin status correspond to changes in gene expression or phenotypes. Many multi-species gene expression datasets in tissues with data from an assay that can serve as a proxy for enhancers are publicly available [37, 83, 103, 104], and more will likely be generated in the near future [105]. Additional multi-species single-cell RNA-seq datasets are being generated from some of these tissues [106, 107] and will likely soon be supplemented by single-cell ATAC-seq. Many of these tissues have been associated with phenotypes that have evolved through gene expression [26, 29], so we can use the gene expression data along with predictions from models like ours to gain insights how these phenotypes evolved. Such insights may also reveal mechanisms underlying diseases associated with these phenotypes [57].

## CONCLUSIONS

The lineage and tissue-specificity of many enhancers limits our ability to quantify the conservation of enhancers through nucleotide-level conservation scores. Therefore, rather than focusing on identifying cases where natural selection is acting on individual nucleotides, we used open chromatin data from tissues of interest to train CNNs for predicting open chromatin and showed that they are able to identify cases where natural selection operates to maintain the combination of TF binding sites that are needed to regulate gene expression. We evaluated the success of our CNNs as well as nucleotide conservation-based and other machine learning-based methods using new criteria that we designed explicitly for evaluating open chromatin conservation prediction. We then used CNNs to predict brain and liver open chromatin conservation across mammals and found that candidate enhancers associated with neuron firing and liver regeneration tended to overlap regions of predicted lineage-specific open or closed chromatin. Our approaches to quantifying enhancer conservation and evaluating methods for this task can be applied to any tissue or cell type with enhancer data available from multiple species. As this is, to our knowledge, the first study to predict open chromatin conservation across more than a few species, we anticipate that our work will serve as a foundation for identifying OCRs whose open chromatin conservation is involved in gene expression evolution, providing insights into transcriptional regulatory mechanisms underlying phenotypes that have evolved through gene expression and their associated diseases.

## METHODS

### Constructing Positive Sets

Data generation and processing is described in **Supplemental Methods**. Mouse experiments were approved by Carnegie Mellon University’s Institutional Animal Care and Use Committee. To obtain OCRs in each species, we intersected the IDR “optimal set” reproducible peaks from each of the brain regions and datasets for brain and each of the liver datasets for liver and defined OCRs to be the intersected peaks that are likely to be enhancers. Specifically, for each species, we selected one set of reproducible open chromatin peaks to be the “base peaks,” used bedtools intersect with the -wa and -u options to intersect it with each of the other reproducible peak sets in series, and then used bedtools closestBed with the -t first and -d options to identify the “base peaks” that overlapped at least one peak from each other set that were over 20kb from the nearest protein-coding TSS (not promoters), at most 1kb long (not super-enhancers), and non-exonic [108]. The base peaks for human brain were the IDR “optimal set” from NeuN+ cells in the primary motor cortex from GSE96949 [65], for the macaque brain were the IDR “optimal set” from the orofacial motor cortex from data we previously generated [61], for the macaque liver were the IDR reproducible peaks across self-pseudo replicates from the first macaque liver replicate from data we previously generated [61], for mouse brain were the IDR “optimal set” from the cortex from the seven-week-old mouse from data we previously generated [66], for mouse liver were the IDR “optimal set” from our mouse liver ATAC-seq dataset, for the rat brain were the IDR “optimal set” from the primary motor cortex from data we previously generated [61], and for the rat liver were the IDR “optimal set” from the second and third rat liver replicates from data we previously generated [61]. To determine the distance from the nearest protein-coding TSS, we used the GENCODE protein-coding TSS’s for human (version 27) and mouse (version M15) [109, 110], the union of the RefSeq rheMac8 protein-coding TSS’s [1] and the human GENCODE protein-coding TSSs mapped to rheMac8 using liftOver [111] for macaque, the union of the RefSeq rn6 protein-coding TSS’s [1] and the mouse GENCODE protein-coding TSSs mapped to rn6 using liftOver [111] for rat, the union of the RefSeq Btau_5.0.1 TSS’s [1] and the human GENCODE protein-coding TSS’s mapped to Btau_5.0.1 with halLiftover [112] on the version 1 Zoonomia Cactus alignment [11] for cow, and the union of the susScr11 TSS’s mapped to susScr3 with liftOver [111] and the human GENCODE protein-coding TSS’s mapped to susScr3 with halLiftover [112] on the version 1 Zoonomia Cactus alignment [11] for pig. To identify non-exonic peaks, we used bedtools [108] subtract with option -A to identify peaks that did not overlap protein-coding exons, where human protein-coding exons were obtained from GENCODE (version 27), mouse protein-coding exons were obtained from GENCODE (version M15) [109, 110], macaque protein-coding exons were obtained from RefSeq for rheMac8, and rat protein-coding exons were obtained from RefSeq for rn6 [1], cow protein-coding exons were obtained from RefSeq for Btau_5.0.1 [1], and pig protein-coding exons were obtained from RefSeq for susScr11 [1] and then mapped to susScr3 with liftOver [111]. We defined the peak summit of an OCR to be the peak summit of the corresponding base peak, and we constructed positive examples by taking likely enhancer peaks summits +/-250bp and their reverse complements. We centered peaks on their summits because previous work has shown that there is a concentration of TF motifs at peak summits [113–115]. By requiring OCRs to be reproducible open chromatin peaks according to IDR, intersecting OCRs across multiple datasets, and filtering OCRs in a conservative way, we limited the number of false positive OCRs being used to train our machine learning models.

### Constructing Negative Set with Non-OCR Orthologs of OCRs

We obtained the non-OCR orthologs of OCRs by obtaining the orthologs of OCRs in each species in each other species for which we had data from the same tissue and filtering the orthologs to include only those that did not overlap open chromatin in the same tissue. For example, mouse chr4:127435564-127436049 does not overlap any OCRs in mouse cortex or striatum, but its human ortholog, chr1:34684126-34684689, is an OCR in both cortex and striatum, so this mouse region was a member of our novel negative set. To ensure that we would have enough negatives for training the model, we created a less conservative set of OCRs, which we called “loose OCRs.” For each species, tissue combination, the loose OCRs are the base peaks that are non-exonic, at least 20kb from a TSS, and at most 1kb long (same criteria used in constructing the positive set) and intersect at least 1 peak from the pooled reads across replicates from each of other datasets that are used for the species, tissue combination; we obtained these loose OCRs using bedtools [108]. We defined the peak summit of a loose OCR to be the peak summit of the corresponding base peak.

We identified orthologs of our loose OCRs in each other species with open chromatin data from the same tissue using halLiftover [112] followed by HALPER [113]. We used these tools instead of liftOver [111] because they map regions using Cactus alignments [116], which, unlike the pairwise alignment liftOver chains, contain many species and account for a wide range of structural rearrangements, including inversions. We ran halLiftover with default parameters on our loose OCRs and their peak summits using the Zoonomia version 1 Cactus alignment [11]. We then constructed contiguous loose OCR orthologs from the outputs of halLiftover by running HALPER with parameters -max_frac 2.0, -min_len 50, and – protect_dist 5. Finally, we used bedtools subract with the -A option [108] to remove the loose OCR orthologs in each species that overlapped peaks from the pooled reads across replicates from any of the datasets from the same tissue in that species.

For the models trained only on mouse sequences, we used only the mouse orthologs of loose OCRs in non-mouse species. For the multi-species brain models, we used the mouse, human, macaque, and rat orthologs of loose brain OCRs from each of the other species, and for the multi-species liver models, we used the mouse, macaque, and rat orthologs of loose liver OCRs from each of the other species. When constructing negatives of the non-OCR orthologs of OCRs, we used sequence underlying the ortholog of the base peak summit +/-250bp and that sequence’s reverse complement. Our negative:positive training set ratios were approximately 1.16:1 for the brain model trained on only mouse sequences, 0.704:1 for the liver model trained on only mouse sequences, 1.49:1 for the multi-species brain model, and 0.822:1 for the multi-species liver model. Construction of additional negative sets is described in **Supplemental Methods**.

### Training Machine Learning Models

Construction of training, validation, and test sets is described in **Supplemental Methods**. We used CNNs [70] for our machine learning model because they have achieved state-of-the-art performance in related tasks [58, 117, 118]; they can learn complex combinatorial relationships between sequences, which we know can play an important role in enhancer activity [48]; and they do not require an explicit featurization of the data, enabling them to learn yet-to-be-discovered sequence patterns that are important for enhancer activity. Our inputs were one-hot-encoded DNA sequences [69], and our outputs were the probability of the sequence being an OCR in the tissue for which the model was trained. We used smaller architectures than those used by many previous studies for related tasks [56, 58, 117, 118] because these previous studies were training multi-task models, where our models were single-task models and therefore likely needed to learn substantially fewer sequence signatures. We first tuned hyper-parameters for the brain model trained on only mouse sequences with our novel negative set by comparing the validation set performances of various models. We did not do an exhaustive hyper-parameter search for any model because our goal was to evaluate the feasibility of training models that had satisfactory performance for all of our evaluation criteria and not to train optimally performing machine learning models. Our final architecture was five convolutional layers with 300 filters of width 7 and stride 1 in each, followed by a max-pooling layer with width and stride 26, followed by a fully connected layer with 300 units, followed by a fully-connected layer that went into a sigmoid. All convolutional layers had dropout 0.2 and L2 regularization 0.00001. The model was trained using stochastic gradient decent with learning rate 0.001, Nesterov momentum 0.99, and batch size 100, and each class was assigned a weight equal to the fraction of the other class in the training set. The model was trained using the training set until there were three consecutive epochs with no improvement in recall at eighty percent precision (or, if there were more positives than negatives, no improvement in specificity at eighty percent NPV) on the validation set. Before training, weights were initialized to be those from a pre-trained neural network with the same hyper-parameters and the negative set randomly down-sampled to be the size of the positive set (or a positive set randomly down-sampled to be the size of the negative set if the positive set was larger). The weights for the pre-training were initialized using Keras’s He normal initializer [119, 120].

We then tuned hyper-parameters for the CNNs for the other negative sets, the CNNs for the liver data, and the CNNs for the multi-species models. We began with the architecture described above and adjusted the number of convolutional filters per layer and the learning rate, ultimately selecting the values that provided the best performance on the validation set. For the model with flanking regions negatives, we used 250 convolutional filters per layer and learning rate 0.001. For the model with the OCRs from other tissues negatives, we used 250 convolutional filters per layer and learning rate 0.0005. For the model with the larger number of random G/C- and repeat-matched negatives, the multi-species brain model, and both liver models, we used 350 convolutional filters per layer and learning rate 0.001. For the models with the smaller number of random G/C- and repeat-matched negatives and the dinucleotide-shuffled negatives, we used 300 convolutional filters per layer and learning rate 0.001. All models were implemented and trained using Keras [120, 121] version 1.2.2 with the Theano backend [122] and evaluated using Scikit-learn [123] and PRROC [124].

### Evaluating Machine Learning Models

#### Evaluating Models’ Lineage-Specific OCR Accuracy

##### OCR Orthologs with Differing Open Chromatin Statuses between Two Species

To obtain OCR orthologs that are open in one species but not in another, we used as positives the sequences and reverse complements of sequences underlying OCRs from one species whose orthologs in the other species have closed chromatin and as negatives sequences and reverse complements of sequences underlying non-OCRs whose orthologs in at least one other species are OCRs. More specifically, for evaluating mouse OCRs whose open chromatin status differs in other species, we used halLiftover [112] followed by HALPER [113] with the same parameters we used previously to identify mouse OCR orthologs in human, macaque, and rat for brain and macaque and rat for liver that did not overlap any of the peaks from the pooled replicates from any of the datasets that we used for identifying OCRs. This gave us 1,570 positive test set examples for evaluating brain models and 3,738 positive test set examples for evaluating liver models. For evaluating mouse non-OCRs whose open chromatin status differs in other species, we used the same approach that we took to construct our novel negative set except that we identified non-OCR orthologs of OCRs instead of loose OCRs. This gave us 2,062 negative test set examples for evaluating brain models and 4,080 negative test set examples for evaluating liver models (**Supplemental Figure 1d, Supplemental Figure 8b**). For evaluating mouse OCRs whose rat orthologs are not OCRs, we identified mouse OCRs whose rat orthologs do not overlap pooled peaks across replicates for any rat OCRs from any of the datasets used to define OCRs; this gave us 674 positive test set examples for evaluating brain models and 2,482 positive test set examples for evaluating liver models. For evaluating mouse closed chromatin regions whose orthologs are open in rat, we used the subset of our novel negative set that came from rat OCR orthologs (not rat loose OCR orthologs); this gave us 890 negative test set examples for evaluating brain models and 2,050 negative test set examples for evaluating liver models (**Supplemental Figure 1e, Supplemental Figure 8b**). For evaluating macaque OCRs whose open chromatin status differs in mouse, we identified macaque OCRs whose mouse orthologs do not overlap peaks from the pooled replicates of any of the datasets we used for identifying mouse OCRs; this gave us 734 positive test set examples for evaluating brain models and 2,384 positive test set examples for evaluating liver models. For identifying macaque non-OCRs whose OCR status differs in mouse, we identified mouse OCR orthologs in macaque that do not overlap peaks from the pooled replicates from any of the datasets we used for identifying macaque OCRs; this gave us 788 negative test set examples for evaluating brain models and 2,228 negative test set examples for evaluating liver models (**Supplemental Figure 1g, Supplemental Figure 8b**). We obtained human and rat OCRs and non-OCRs whose OCR status is different in mouse using same process that we used for macaque. We obtained 416 positive and 896 negative test set examples for evaluating brain models (**Supplemental Figure 1h**). For evaluating rat OCRs and non-OCRs whose mouse orthologs have different OCR statuses, we obtained 990 positive and 676 negative test set examples for evaluating brain models and 2,050 positive and 2,482 negative test set examples for evaluating liver models (**Supplemental Figure 1i, Supplemental Figure 8b**).

##### Clade- and Species-Specific OCRs

We defined a clade-specific OCR in a tissue as an OCR whose ortholog is open in that tissue in every species within a clade for which we have data and closed in every other species for which we have data, and we defined a species-specific OCR as an OCR whose ortholog is open a species for which we have data and is closed in the most closely related species for which we have data. Laurasiatheria data was used only for comparing mouse-only to multi-species liver models (**Figure 3e**). More specifically, we identified clade-active OCRs in each clade – OCRs in a “base species” whose ortholog in the other species in that clade (if there was another) overlaps an open chromatin peak from the pooled reads across replicates in all of the datasets we used in that tissue from that species. We did not require the OCR ortholog in the non-base species to overlap a reproducible open chromatin peak so that we could have at least one hundred test set examples for each evaluation. We chose the “base species” to be the species in each clade with the highest-quality genomes – mouse for Glires for brain and liver, human for Euarchonta for brain, macaque for Euarchonta for liver, and cow for Laurasiatheria for liver. We then identified the subset of clade-active peaks from the base species whose orthologs in all species in the other clade do not overlap any open chromatin peaks from the pooled replicates from any of the datasets we used to identify OCRs; these were our clade-specific OCRs. To obtain clade-specific non-OCRs for a clade, we identified orthologs of clade-active OCRs from the other clade in the base species in the clade whose orthologs in all species in the clade did not overlap open chromatin peaks from pooled replicates in any of the datasets we used for identifying OCRs. The sequences of clade-specific OCRs and non-OCRs used for evaluating the models were those from the base species and their reverse complements. This gave us 230 Glires-specific test set brain OCRs, 134 Glires-specific test set brain non-OCRs (**Supplemental Figure 1f, Supplemental Figure 4a, Figure 3a**), 134 Euarchonta-specific test set brain OCRs, 230 Euarchonta-specific test set brain non-OCRs (**Supplemental Figure 1j, Supplemental Figure 4b, Figure 3a**), 1,024 Glires-specific test set liver OCRs, 1,826 Glires-specific test set liver non-OCRs (**Supplemental Figure 8b,** **Figure 3b**), 1,826 Euarchonta-specific test set liver OCRs, 1,024 Euarchonta-specific test set liver non-OCRs (**Supplemental Figure 8b,** **Figure 3b**), 77 Laurasiatheria-specific test set liver OCRs, and 86 Laurasiatheria-specific test set liver non-OCRs (**Figure 3e**). When evaluating the multi-species models, we combined the clade-specific OCRs and non-OCRs from Euarchonta and Glires.

When evaluating species-specific OCRs and non-OCRs, we identified mouse OCRs and non-OCRs whose orthologs’ open chromatin status differs in rat, rat OCRs and non-OCRs whose orthologs’ open chromatin status differs in mouse, human OCRs and non-OCRs whose orthologs’ open chromatin status differs in macaque, and macaque OCRs and non-OCRs whose orthologs’ open chromatin status differs in human. We obtained 66 human-specific brain OCRs, 188 human-specific brain non-OCRs, 188 macaque-specific brain OCRs, and 66 macaque-specific brain non-OCRs (**Figure 3a**). We did not include macaque-specific OCRs and non-OCRs when evaluating the multi-species liver model because we did not have liver open chromatin from any other Euarchonta species. We combined the species-specific OCRs and non-OCRs from different species when evaluating the multi-species models (**Figures 3a-b**).

#### Evaluating Models’ Tissue-Specific OCR Accuracy

To evaluate the performance of models trained in one tissue on OCRs from another tissue, we defined our positives to be OCRs that are shared between the two tissues (shows our models were not learning only the sequences involved in tissues-specific open chromatin), and we defined our negatives to be OCRs in the evaluation tissue that do not overlap OCRs in the training tissue (shows our models were not learning only sequences involved in general open chromatin). More specifically, we used bedtools intersectBed with options -wa and -u [108] to identify OCRs from our training tissue that overlap OCRs from the evaluation tissue. For brain models, we obtained 1,040 positives for mouse, 846 positives for macaque, and 1,770 positives for rat (**Supplemental Figure 5**). For liver models, we obtained 2,012 positives for mouse, 946 positives for macaque, and 1,130 positives for rat (**Supplemental Figure 8c**). We used bedtools subtractBed with option -A [108] to identify liver OCRs that do not overlap any open chromatin from pooled replicates in any of the datasets that were used to generate the brain OCRs and brain OCRs that do not overlap open chromatin from pooled replicates in any of the datasets that were used to generate liver OCRs. For brain, we obtained 3,382 negatives for mouse, 1,898 negatives for macaque, and 3,518 negatives for rat (**Supplemental Figure 5**). For liver, we obtained 2,212 negatives for mouse, 1,428 negatives for macaque, and 2,942 negatives for rat (**Supplemental Figure 8c)**. We combined the data from all three species for evaluating the multi-species models (**Figures 3a-b**).

For the negative set comparison, we also compared the distributions of test set predictions for the brain OCRs that do not overlap liver OCRs, the brain OCRs that overlap liver OCRs, the liver OCRs that do not overlap brain OCRs, and the negative set. We defined these groups of OCRs as we did for other evaluations, and we used predictions for sequences and their reverse complements. We compared the distributions for the brain OCRs that overlap liver OCRs to the liver OCRs that do not overlap brain OCRs using a Wilcoxon rank-sum test and multiplied the p-values by 6 to do a Bonferroni correction.

#### Evaluating if Models’ Predictions Had Phylogeny-Matching Correlations

To evaluate the relationship between OCR ortholog open chromatin status and phylogenetic distance, we identified test set mouse OCR orthologs in all of the fifty-six Glires species from Zoonomia, predicted the open chromatin statuses of those orthologs, and computed the correlation between those predictions and the species’ phylogenetic divergences from mouse. This provides us with an approximate measure of how predicted OCR ortholog open chromatin statuses change over evolution. We identified the test set mouse OCR orthologs and OCR summit orthologs in Glires using halLiftover [112] with the Zoonomia version 1 Cactus alignment [11, 116]; we used brain OCRs for evaluating brain OCR models and liver OCRs for evaluating liver OCR models. We next constructed contiguous orthologs from the outputs of halLiftover using HALPER [113] with parameters -max_frac 2.0, -min_len 50, and -protect_dist 5. We constructed inputs for our models from the contiguous OCR orthologs by using bedtools fastaFromBed [108] with fasta files downloaded from NCBI [4, 125] and the UCSC Genome Browser [126] to obtain the sequences underlying their summit orthologs +/-250bp. We constructed the reverse complements of sequences, used our models to predict each sequence and its reverse complement’s open chromatin status, and averaged the predictions between the forward and reverse strands. We then removed all predictions from OCRs with orthologs in less than one quarter of species. After that, for each model, we computed the mean OCR ortholog open chromatin status prediction and the standard deviation of predictions across all remaining OCR orthologs in each species. We finally computed the Pearson and Spearman correlations between these means and standard deviations of predictions and the millions of years since divergence from mouse, which we obtained from TimeTree [127]. We did this for brain OCR orthologs using brain models trained on mouse sequences from each negative set and the multi-species brain model as well as for liver OCR orthologs using the liver model trained on only mouse sequences and the multi-species liver model.

### Comparing Predictions to Mean Conservation Scores

We compared the predictions to mean conservation scores by identifying OCR orthologs with conserved and non-conserved open chromatin status between species, computing the mean conservation scores of those OCR orthologs, and comparing those scores to the predicted open chromatin status of those OCR orthologs. We defined an OCR ortholog with conserved open chromatin status between mouse and another species as a mouse OCR whose ortholog in the other species overlaps an OCR in the same tissue. For mouse brain test set OCRs, we identified 441 OCRs with conserved open chromatin status in macaque, 195 OCRs with conserved open chromatin status in human, and 670 OCRs with conserved open chromatin status in rat. For mouse liver test set OCRs, we identified 689 OCRs with conserved open chromatin status in macaque and 580 OCRs with conserved open chromatin status in rat. We defined an OCR ortholog with non-conserved open chromatin status between mouse and another species as a mouse OCR whose ortholog in another species does not overlap any OCR from the pooled replicates from any dataset used in defining an OCR in that tissue in that species. For mouse brain test set OCRs, we identified 394 OCR orthologs with non-conserved open chromatin status in macaque, 448 OCR orthologs with non-conserved open chromatin status in human, and 338 OCR orthologs with non-conserved open chromatin status in rat. For mouse liver test set OCRs, we identified 1,114 OCR orthologs with non-conserved open chromatin status in macaque and 1,241 OCR orthologs with non-conserved open chromatin status in rat. We think that the differences in numbers of OCR orthologs with conserved and non-conserved open chromatin status between species is due not only to differences in evolutionary relatedness but also to differences between species in numbers of datasets used to define OCRs and differences in sequencing depths of those datasets [38, 61, 65].

We used our models to predict the OCR ortholog open chromatin status for the open chromatin status-conserved and open chromatin status non-conserved OCR orthologs in the non-mouse species and compared it to the conservations scores of the mouse OCRs. We computed mean conservation scores of the mouse OCRs by calculating the mean PhastCons [13] and PhyloP [14] scores at the peak summits +/-250bp. We evaluated if the distributions of the predictions and each type of conservation score differed between the open chromatin status-conserved and open chromatin status non-conserved orthologs using a Wilcoxon rank-sum test. We did a Bonferroni correction by multiplying all p-values by 20 (2 conservation score comparisons and 2 model predictions comparisons – models trained on only mouse sequences and multi-species models – for 5 species, tissue pairs).

We then evaluated whether the predictions were more effective than the mean conservation scores at differentiating between open chromatin status-conserved and open chromatin status non-conserved OCR orthologs. We first averaged the predictions of the sequence underlying the non-mouse OCR ortholog’s summit +/-250bp and its reverse complement so that each OCR ortholog would have a single prediction value. We next combined our open chromatin status-conserved and open chromatin status non-conserved OCR orthologs and ranked them according to each of PhastCons score, PhyloP score, and OCR ortholog open chromatin status prediction. We then did a Wilcoxon sign-rank test to compare the ranking distributions of the open chromatin status-conserved OCR orthologs between the OCR ortholog open chromatin status predictions and each type of conservation score. We also did this for the ranking distributions of the open chromatin status non-conserved OCR orthologs. We did this for predictions made by the models trained using only mouse sequences and by the multi-species models. Finally, we did a Bonferroni correction by multiplying all p-values by 40 (2 conservation score comparisons for each of open chromatin status-conserved and open chromatin status non-conserved OCR orthologs in 2 tissues for 5 species, tissue pairs). Processing of liver H3K27ac ChIP-seq data for comparing predicted open chromatin conservation and conservation scores to conservation of H3K27ac ChIP-seq is described in **Supplemental Methods**.

### Clustering OCRs

Identifying genes with Rodent-specific expression OCRs with predicted Rodent-specific open chromatin is described in **Supplemental Methods**. To generate OCR clusters for OCRs from a tissue, we mapped the OCRs from each species across all of the Boreoeutheria from Zoonomia [4] except for *Manis tricuspis*, filtered the OCRs, and clustered the OCRs. Specifically, we mapped OCRs from each species with OCRs using halLiftover with the Zoonomia version 1 Cactus [11] and constructed contiguous orthologs from halLiftover outputs using HALPER [113] with settings -max_frac 2.0, -min_len 50, and -protect_dist 1. 5. We then used the model corresponding to the tissue for which the OCR was generated to make predictions for the OCR orthologs and OCR orthologs’ reverse complements in all of the Boreoeutheria that we mapped to except for *Galeopterus variegatus*, *Hippopotamus amphibius*, *Monodon monoceros*, and *Platanista gangetica*. For each OCR ortholog, we set the prediction to be the average between the prediction for the ortholog and the prediction for its reverse complement.

We filtered OCRs to ensure that we did not have redundant OCRs and to ensure that we had usable OCR orthologs in enough species to use predictions in each species as features for clusters. First, for brain, we removed all human OCRs whose mouse ortholog overlapped a mouse brain OCR, all macaque OCRs whose mouse ortholog overlapped a mouse brain OCR or whose human ortholog overlapped a human brain OCR, and all rat OCRs whose mouse ortholog overlapped a mouse brain OCR, whose human ortholog overlapped a human brain OCR, or whose macaque ortholog overlapped a macaque brain OCR. Likewise, for liver, we removed all macaque OCRs whose mouse ortholog overlapped a mouse liver OCR and all rat OCRs whose mouse ortholog overlapped a mouse liver OCR or whose macaque ortholog overlapped a macaque liver OCR. After that, we removed all remaining OCRs that did not have a usable ortholog in at least half of all species or at least one quarter of each of Euarchonta, Glires, and Laurasiatheria.

We clustered the remaining OCRs using the prediction of each species’s OCR ortholog’s open chromatin status as a feature and treating species without usable orthologs as missing data. We first used k-means clustering with k = 9,000 (brain) or 12,000 (liver) to cluster the OCRs into small clusters; cluster centroid values for each species were defined as the mean of all OCR ortholog activity predictions for that species in the cluster. We then used affinity propagation clustering [128], implemented in scikit-learn [123], with preference = -0.6 to cluster the outputs of the k-means clustering, which we defined as the centroids of the small clusters, into larger clusters. In both clustering steps, we defined the distance between two OCRs as 1 minus the cosine similarity between the vectors of enhancer activity predictions in the species for which both had usable orthologs. We ultimately obtained 102 brain clusters and 103 liver clusters.

### Identifying Enhancer Sets with More Overlap with Clusters than Expected by Chance

To identify enhancer sets with more overlap with a cluster than expected by chance, we first obtained the enhancer sets from the supplemental information of the relevant manuscripts. For the mouse neuron firing candidate enhancers, we used the regions in Tables 11, S12 and S13 from a recent study of enhancers activated during mouse neuron firing [94] and used liftOver [111] to map the coordinates from mm9 to mm10. For the human GABAergic neuron activity candidate enhancers, we used the regions in Supp Data 12 from a recent study of enhancers activated during human neuron activation [95] and used liftOver to map the coordinates from hg19 to hg38 [111]. For the mouse liver regeneration candidate enhancers, we used various subsets of regions from Supplemental Table 1 from a recent study of enhancers activated during liver regeneration [96]; for each category of regions in **Supplemental Table 19**, we required the FDR to be less than 0.05 and the log2 fold-change to be greater than 1 (for ↑) or less than 1 (for ↓). We then used LOLA [129] to run a hypergeometric test to evaluate the statistical significance of the overlap of each enhancer set with each cluster from the relevant tissue that is open in the relevant species; our query was our enhancer set, our database was our list of bed files with the relevant cluster locations, our universe was our list of OCRs in the relevant species, tissue combination, and we set “redefineUserSets” to TRUE. We did a Bonferroni correction by multiplying all p-values by 391, which was the total number of tests.

We wanted to determine if our enrichment for overlaps with clusters with lineage-specific open chromatin or lack of open chromatin could be partially explained by OCRs overlapping enhancer sets having fewer usable orthologs or lower conservation. To evaluate this, we used bedtools [108] to identify the OCRs from each relevant tissue, species combination overlapping each enhancer set. We next computed the number of usable orthologs, the average PhastCons [13] score for the peak summit +/-250bp, and the average PhyloP [14] score for the peak summit +/-250bp for all OCRs from the relevant tissue, species combination as well as for the OCRs overlapping each enhancer set. Then, for each enhancer set, we used a Wilcoxon rank-sum test to compare the numbers of usable orthologs and the PhastCons and PhyloP scores between the OCRs overlapping the enhancer set and the full set of OCRs. We considered the difference to not be statistically significant if the nominal p-value was greater than or equal to 0.05.

## DECLARATIONS

### Ethics approval and consent to participate

All human data is publicly available, and all other animal data is either publicly available or was collected in experiments approved by Carnegie Mellon University’s Institutional Animal Care and Use Committee.

### Consent for publication

Not applicable.

### Availability of data and materials

Human DNase hypersensitivity data analyzed in this study was downloaded from the ENCODE portal (https://www.encodeproject.org/) [38, 64], and human ATAC-seq data analyzed in this study was downloaded from Gene Expression Omnibus accession GSE96949 [65]. Macaque and rat ATAC-seq data from data we previously generated [61] and mouse liver ATAC-seq data in this manuscript can be accessed in Gene Expression Omnibus accession GSE159815 with token gzkbqwyobvmvfsd. Mouse brain data analyzed in this study can be accessed in Gene Expression Omnibus accession GSE161374 with token cropkwsgnnyxhgh. Publicly available mouse liver ATAC-seq data was downloaded from China National Gene Bank accession CNP0000198 [130]. Other mouse ATAC-seq data was downloaded from China National Gene Bank accession CNP0000198 [130] and the ENCODE portal (https://www.encodeproject.org/) [131]. Cow and pig ATAC-seq data was obtained from the authors of [72]. Mouse and macaque H3K27ac ChIP-seq data was downloaded from ArrayExpress accession E-MTAB-2633 [37]. Code and cluster images can be found at https://github.com/pfenninglab/OCROrthologPrediction, and additional code and cluster information can be obtained from the authors upon request.

### Competing interests

The authors declare that they have no competing interests.

### Funding

The Carnegie Mellon University Computational Biology Department Lane Fellowship supported I.M.K. The Alfred P. Sloan Foundation Research Fellowship supported I.M.K., M.E.W., and A.R.P. The NIH NIDA DP1DA046585 supported I.M.K., D.E.S., M.E.W., A.J.L., A.R.B., M.K., and A.R.P. The NSF Graduate Research Fellowship Program under grants DGE1252522 and DGE1745016 supported A.J.L. The Carnegie Mellon Neuroscience Institute Presidential Fellowship supported M.K.

### Authors’ contributions

I.M.K. and A.R.P. designed the study with assistance from M.E.W. A.J.L. collected the mouse liver ATAC-seq data with assistance from A.R.B. I.M.K. processed the open chromatin and histone modification data with assistance from M.E.W. I.M.K. did the machine learning model training, machine learning model evaluation, and other computational method implementation except for the clustering. I.M.K. and A.R.P. designed the machine learning model evaluation criteria with assistance from M.K. and M.E.W. I.M.K. and D.E.S. developed the clustering method with assistance from M.K. and A.R.P., and D.E.S. implemented the clustering method. I.M.K. wrote the manuscript with assistance from D.E.S. and A.J.L. All authors reviewed and helped revise the manuscript.

## Acknowledgements

We would like to thank the other members of the Pfenning Lab for useful discussions and suggestions and J. Ma and Y. Zhang for feedback on the manuscript. We would also like to thank the members of the Paten Lab and the Zoonomia Project for assistance with using Cactus multi-species alignments and providing us with early access to these alignments. This work used the Extreme Science and Engineering Discovery Environment (XSEDE), through the Pittsburgh Supercomputing Center Bridges Compute Cluster, which is supported by National Science Foundation grant number TG-MCB190067.

## Notes

https://github.com/pfenninglab/OCROrthologPrediction

### Competing Interest Statement

The authors have declared no competing interest.

### Summary of Updates

We now compare machine learning models to conservation scores in their ability to achieve both lineage-specific open chromatin accuracy and tissue-specific open chromatin accuracy.

