## SupplementalText for "Inferring mammalian tissue-specific regulatory conservation by predicting tissue-specific differences in open chromatin"

**CONTENTS**

1. Supplemental Notes
2. Supplemental Materials and Methods
3. Supplemental Figure Captions
4. Supplemental Tables
5. Supplemental References

**SUPPLEMENTAL NOTES**

Additional Evaluations of Lineage-Specific OCR Accuracy for Machine Learning Models Trained on Only Mouse Sequences

In addition to evaluating performance on OCRs and non-OCRs in non-mouse species whose mouse ortholog has a different open chromatin status, we evaluated each model on the subset of mouse brain OCRs whose orthologs in at least one other species are closed (subset of positive set for all models) and the mouse brain closed chromatin regions whose orthologs in at least one other species are open in brain. Interestingly, although all models performed decently on these genomic regions (AUC > 0.65, AUPRC > 0.55), none of the models worked as well on these genomic regions as they did on the test set corresponding to the regions used in training them, and the best-performing model on its own test set – the model trained with dinucleotide-shuffled negatives – was the worst-performing model on these regions (**Supplemental Figure 1d**). We obtained similar results for the subset of mouse brain open and closed chromatin regions whose rat orthologs have the opposite brain open chromatin status (**Supplemental Figure 1e**), showing that some of these models are capable of accurately predicting differences in brain open chromatin between closely related species. These findings reveal a bias in some existing methods for accurately predicting open chromatin conservation and show that these biases can be mitigated by using other negative sets.

Having demonstrated the ability to train a model in one species and predict in another, we evaluated the ability of the models to accurately predict clade-specific brain open and closed chromatin regions – regions whose brain open chromatin status is shared across species in one clade for which we have data but not shared by any species in another clade for which we have data (**Supplemental Figure 1a**). We focused our analyses on the two major clades for which we had experimentally determined open chromatin data (**Figure 1**): Glires (the clade comprising all rodents and lagomorphs, including mouse and rat) and Euarchonta (the clade comprising all primates — including human and macaque — and their closest relatives, colugos and tree shrews). We found that all models obtained decent performance for Glires-specific open and closed chromatin regions (AUC > 0.75, AUNPV-Specificity > 0.7), with the model trained on the dinucleotide-shuffled brain OCR negatives providing the worst performance and the models trained with our negative set and the larger G/C- and repeat-matched negative set providing the best performance (**Supplemental Figure 1f**). Because we trained our models on sequences from mouse (clade: Glires), we also evaluated how well our models work on clade-specific brain open and closed chromatin regions from Euarchonta species, as this clade was not used in training. We found that the models obtained slightly worse performance on Euarchonta-specific brain open and closed chromatin regions than they did on Glires-specific brain open and closed chromatin regions, with similar relative performances for the models trained on different negative sets (**Supplemental Figure 1j**). Since many phenotypes are clade-specific, these results demonstrate the necessity of evaluating models for OCR ortholog open chromatin prediction on clade-specific open and closed chromatin regions for clades used and not used in training.

<u>Re-Calibrating Models with Our Negative Set Usually Does Not Substantially Improve Performance</u>

Since, for many applications, we need to make a binary classification as to whether a region is open in brain, we also investigated how well-calibrated our models are. We found that models trained on some negative sets — including flanking regions, OCRs in other tissues, the smaller G/C- and repeat-matched set, and dinucleotide-shuffled brain OCRs — tended to do better on clade-specific OCRs than on clade-specific closed chromatin regions. On the other hand, the models trained with the larger G/C- and repeat-matched set and our novel negative set tended to do better on clade-specific closed chromatin regions than on clade-specific OCRs. We tried re-calibrating all of the models with the positive training set and the training set from our novel negative set. For the models trained on all negative sets except for ours, this led to an increase in specificity and a decrease in sensitivity. In general, the increase in specificity was similar to the decrease in sensitivity (**Supplemental Tables 1-6**), but, for the smaller G/C- and repeat-matched region negative set, the increase in specificity was substantially larger than the decrease in sensitivity (**Supplemental Figure 2**). Thus, while some models were poorly calibrated, re-calibrating models with our negative set usually had limited utility.

<u>Machine Learning Models Predict OCR Orthologs' Open Chromatin Status Significantly More Accurately than Mean Conservation Scores</u>

To do quantify the extent to which our machine learning model trained with our novel negative set can predict differences in open chromatin conservation relative to conservation scores, we identified test set mouse brain and liver OCRs whose macaque orthologs do and do not overlap OCRs in brain and liver, respectively, and computed the mean conservation scores of these OCRs [1, 2] as well as the predictions on test set macaque orthologs of the machine learning models trained with our novel negative set in the corresponding tissue. We found that mean conservation scores and model predictions tended to be higher for the macaque orthologs for which open chromatin status was conserved than those for which open chromatin status was not conserved (**Supplemental Tables 7-8**). For each tissue, we then ranked the macaque OCR orthologs based on their mean conservation scores and their model predictions, with the highest rank corresponding to the highest score or open chromatin status prediction. For the open chromatin status-conserved OCRs in each tissue, we used a Wilcoxon signed-rank test to evaluate whether these OCRs tended to have higher ranks for our predictions than they do for mean conservation scores; we found that the ranks were significantly higher for our predictions (brain predictions vs. PhastCons scores: $3.69 \times 10^{-4}$, brain predictions vs. PhyloP scores: $9.30 \times 10^{-6}$, liver predictions vs. PhastCons scores: $6.25 \times 10^{-17}$, liver predictions vs. PhyloP scores: $1.35 \times 10^{-22}$). For each tissue, we also used a Wilcoxon signed-rank test to evaluate whether the OCR orthologs without open chromatin tended to have lower ranks for our predictions than they do for mean conservation scores, and we found that the ranks were significantly lower for our predictions (brain predictions vs. PhastCons scores: 6.88 x $10^{-4}$, brain predictions vs. PhyloP scores: 1.85 x $10^{-7}$, liver predictions vs. PhastCons scores: 3.63 x $10^{-11}$, liver predictions vs. PhyloP scores: 3.42 x $10^{-18}$). We repeated this for human and rat orthologs of mouse brain OCRs with conserved and non-conserved open chromatin statuses and for rat orthologs of mouse liver OCRs with conserved and non-conserved OCR statuses and obtained similar results (**Supplemental Tables 7-9**). This shows that using machine learning models for predicting open chromatin status conservation of OCR orthologs can be more accurate than using mean sequence conservation scores.

Machine Learning Models Learned Motifs of Brain Transcription Factors

To determine what sequence patterns our models were prioritizing, we ran DeepLIFT with the rescale rule [3] followed by TF-MoDISco [4] on the positive examples from the validation set from each model and compared the results to known motifs. All of the models seemed to have learned motifs of TFs that are known to play important roles in the brain, including Ctcf [5, 6], Fos [7-9], Egr2 [10, 11], and Rfx4 [12-14] (**Supplemental Figure 6**). All of the models except for those trained with flanking region negatives (**Supplemental Figure 6a**) and those trained with dinucleotide-shuffled brain OCR negatives (**Supplemental Figure 6e**) seemed to also have learned the motif of Mef2c, a TF with multiple roles in the brain [15-17] (**Supplemental Figure 6**). The model with OCRs in other tissues as negatives seemed to have learned the depletion of motifs of multiple TFs whose human orthologs are not expressed in the brain, including Hnf4g, Nr5a1, Elf3, and Foxd2, and the model with model with the larger number of G/C- and repeat-matched negatives seemed to have learned a depletion of the motif for Nr2f6, which has very low expression in the brain [18, 19] (**Supplemental Figures 6b-c**). The model with the dinucleotide-shuffled brain OCR negatives seemed to have learned the motif for Bcl6 [20, 21], but this motif consists almost exclusively of G's, so it might be indicative of many consecutive G's being more common in brain OCRs than in shuffled brain OCRs (**Supplemental Figure 6e**). The model trained with our novel negative set also seemed to have learned the motif for Dbp, which has been implicated in circadian rhythms [22, 23]; two slightly different Rfx motifs (also learned by the model with the smaller number of G/C- and repeat-matched negatives), which is not surprising because multiple Rfx TFs play important roles in the brain [12, 14, 24]; and a depletion of the motif for Thra (**Supplemental Figures 6d, f**). It is possible that these apparent differences in motifs learned by the models caused their differences in performance.

<u>Approach to Evaluating Machine Learning Models for OCR Ortholog Open Chromatin Status Prediction Can Be Applied to Any Tissue or Cell Type with Open Chromatin Data from Multiple Species</u>

Although we prototyped our approach to evaluating machine learning models for predicting open chromatin status of OCR orthologs in the brain, this approach can be applied to any tissue or cell type with open chromatin data from multiple species. We therefore applied it to another tissue, the liver, and found that our novel approach to negative set construction also worked well for most metrics. To do this, we first generated a new mouse liver open chromatin dataset and found that it was high-quality (**Supplemental Figure 8a**, TSS enrichment for replicate 1 = 17.27, TSS enrichment for replicate 2 = 16.31, rescue ratio = 1.02, self-consistency ratio = 1.26). We then defined our positive set for liver as the 250bp in each direction of peak summits of our mouse liver ATAC-seq peaks that overlapped liver ATAC-seq peaks from CNP0000198 [25]. We obtained negatives by mapping rat and macaque liver ATAC-seq data from the Pfenning Lab [26] to mouse and identifying the mouse orthologs that did not overlap mouse liver ATAC-seq peaks. We found that the model achieved high lineage-specific and tissue-specific OCR accuracy (AUC > 0.7, AUPRC > 0.65, **Supplemental Figures 8b-c**). We also determined if our predictions had phylogeny-matching correlations by obtaining orthologs of the mouse liver OCRs in all of the Glires from the Zoonomia project [27, 28] and predicting their open chromatin statuses. As with brain, we found a strong negative correlation between the predicted mean liver OCR ortholog open chromatin status in

Glires and those species' divergence from mouse (**Supplemental Figure 8d**) and a strong positive correlation between standard deviation of predicted liver OCR ortholog open chromatin status in Glires and those species' divergence from mouse (**Supplemental Figure 8e**). In addition, we interpreted the model using DeepLIFT with the rescale rule [3] followed by TF-MoDISco [4] and found that the model seemed to have learned motifs of multiple known liver TFs, including Ctcf [5, 29], Ppara [30-32], and Cebpa [33, 34], as well as a depletion of the motif for Wt1, which is not expressed in liver [18, 19] (**Supplemental Figure 8f**). Finally, for each species with open chromatin data, we identified OCRs overlapping H3K27ac ChIP-seq peaks from https://www.ebi.ac.uk/research/flicek/publications/FOG15 [35] and compared the predicted activity of OCR orthologs in each other species with H3K27ac ChIP-seq in the orthologs with H3K27ac ChIP-seq to those in the orthologs without H3K27ac ChIP-seq. We found that the predictions in OCR orthologs for which H3K27ac ChIP-seq was conserved tended to be higher than the predictions in OCR orthologs for which H3K27ac ChIP-seq was not conserved (**Supplemental Table 13**). This illustrates that our novel approach to constructing a negative set for open chromatin status prediction of OCR orthologs works well in multiple tissues.

<u>Open Chromatin Predictions Do Not Seem to Be Associated with Genome Quality</u>

Since the Zoonomia genomes vary in quality, we evaluated whether our open chromatin status predictions are associated with genome quality [27, 28, 36]. We computed the correlation between mean predicted brain open chromatin status across the mouse brain OCR orthologs in each Glires species and scaffold and contig N50's. We found a weak Pearson correlation and even weaker Spearman correlation between the scaffold and contig N50's and the mean predicted brain open chromatin status (**Supplemental Figure 10a**). We repeated this process in for the liver predictions of the mouse liver OCR orthologs and obtained similar results (**Supplemental Figure 10b**). To demonstrate that mean predicted mouse OCR ortholog open chromatin status has a stronger relationship with divergence from mouse than it does with genome quality, we created generalized linear models for mean predicted mouse OCR ortholog open chromatin status with covariates for divergence from mouse and scaffold or contig N50. The coefficients for divergence from mouse were all statistically significantly different from zero and larger in magnitude than the coefficients for scaffold or contig N50, and the coefficients for scaffold or contig N50 were never statistically significantly different from zero (**Supplemental Table 14**). These results suggest that lower-quality genomes are not strongly associated with lower OCR ortholog open chromatin status predictions.

To further evaluate the relationship between genome quality and our predictions, we investigated whether the extent to which OCR ortholog open chromatin status predictions vary within a species is associated with genome quality. To do this, we computed the correlation between standard deviation of predicted brain open chromatin status across the mouse brain OCR orthologs in each of the Glires and scaffold and contig N50's. We found a weak negative Pearson correlation and even weaker negative Spearman correlation between the scaffold N50's and the standard deviation of predicted brain open chromatin status; for contig N50, the Pearson correlation was weak and negative, while the Spearman correlation was weak and positive (**Supplemental Figure 10c**). We repeated this process for the liver open chromatin status predictions of the mouse liver OCR orthologs and obtained similar results except that the Spearman correlation between the contig N50 and standard deviation of predicted open chromatin status was weak and negative (**Supplemental Figure 10d**). To demonstrate that standard deviation of predicted mouse OCR ortholog open chromatin status had a stronger relationship with divergence from mouse than it did with genome quality, we created generalized linear models for standard deviation of predicted mouse OCR ortholog open chromatin status with covariates for divergence from mouse and scaffold or contig N50. The coefficients for divergence from mouse were all statistically significantly different from zero and larger in magnitude than the coefficients for scaffold or contig N50 (**Supplemental**

**Table 15**). These results further demonstrate that genome quality does not substantially influence our OCR ortholog open chromatin status predictions.

Multi-Species Models Learn Additional Brain and Liver TF Motifs

We found that our multi-species brain and liver models seemed to have learned motifs of brain and liver TFs, respectively. When interpreting the multi-species brain model, in addition to the motifs that we found for the model trained on only mouse sequences, we found a depletion of the motifs for Nr1i3 and Pit1, both of which are not expressed in human brain (**Supplemental Figure 9c**) [18, 19]. When interpreting the multi-species liver model, in addition to the motifs that we found for the model trained on mouse sequences, we also found the motifs for additional TFs that are known to be involved in the liver, including Hnf4a [37-39], Foxk1 [40, 41], Ets2 [42, 43], Sp1 [44, 45], Onecut1 [46, 47], Bcl6 [48, 49], and Nfe2l2 [50, 51], as well as a depletion of the motif for Dbx1, which is not expressed in human liver [18, 19], and a depletion of the motif for Zfp637 (**Supplemental Figure 9d**). These results suggest that our multi-species models are learning the importance of relevant sequence features to their tasks and not only learning general patterns of genome sequence content.

Multi-Species Machine Learning Models Make Significantly More Accurate Predictions than Mean Conservation Scores

We compared the test set predictions of our multi-species models to those made by mean conservation scores. First, we found that our model predictions for non-mouse orthologs of mouse brain and liver OCRs whose OCR status is conserved tended to be higher than for non-mouse orthologs of mouse brain and liver OCRs whose OCR status is not conserved (**Supplemental Tables 7-8**). Then, for each tissue, we ranked the macaque OCR orthologs based on their model predictions, with the highest rank corresponding to the highest score or open chromatin status prediction. For the open chormatin status-conserved OCRs, we evaluated whether these OCRs tended to have higher ranks for our multi-species model predictions than they did for mean conservation scores; we found that the ranks were significantly higher for our predictions (brain predictions vs. PhastCons scores: $7.80 \times 10^{-5}$, brain predictions vs. PhyloP scores: $1.53 \times 10^{-6}$, liver predictions vs. PhastCons scores: $3.68 \times 10^{-22}$, liver predictions vs. PhyloP scores: $1.25 \times 10^{-27}$). For each tissue, we also evaluated whether the OCR orthologs without open chromatin tended to have lower ranks for our multi-species model predictions than they do for mean conservation scores, and we found that the ranks were significantly lower for our predictions (brain predictions vs. PhastCons scores: $4.34 \times 10^{-5}$, brain predictions vs. PhyloP scores: $8.08 \times 10^{-9}$, liver predictions vs. PhastCons scores: $7.15 \times 10^{-14}$, liver predictions vs. PhyloP scores: $9.82 \times 10^{-22}$) (**Supplemental Tables 7-8, 16**).

Additional Limitations of Our Method

Despite the numerous advantages of predicting open chromatin status with short sequences, using shorter input sequences also has limitations. Some enhancers, such as super-enhancers, are much longer than 500 base pairs, and such enhancers have been shown to play important roles in the brain [52]. In addition, open chromatin status can be affected by long-range interactions with DNA sequences that are more than a few hundred base pairs away from open chromatin peak summits [53]. For example, one study showed that many variants associated with open chromatin occur at least a few hundred base pairs away from OCRs [54]. Encouragingly, our knowledge of 3D genome structure is advancing rapidly, so incorporating such information into machine learning models may be feasible in the near future. Furthermore, open chromatin status changes over evolutionary history can be affected by factors not influenced by local sequence, such as changes in TFs' protein structures that affect their ability to interact with DNA or other TFs [55], so any model with only DNA sequence underlying OCRs as input will not be able to predict every OCR ortholog open chromatin status difference between species.

In addition, training and evaluating any machine learning model using regulatory genomics data from multiple species is inherently limited because raising different species in the same type of controlled environment is infeasible, and this can make differentiating between lineage-specific OCRs and confounding factor-specific OCRs difficult. For example, the activity of many enhancers has been associated with aging [56, 57]. Although all of our data were from adults, the mouse [25, 58] and rat data were from younger adults, whereas the human and macaque data came from a combination of younger and older adults [26, 59, 60]. Part of our motivation for conservatively defining clade-specific OCRs was the desire to prevent Glires-specific OCRs from being young adult-specific OCRs. In addition, time of day and the amount of time since waking up has been shown to affect enhancer activity [61], and controlling for these factors is challenging when obtaining post-mortem human data or data from different animal colonies. Although our macaque and rat samples were collected approximately two hours after the animals woke up, time of day of collection relative to sleep cycle for the remaining samples used was either not described or not able to be controlled [25, 26, 58-60]. Thus, some individual OCR ortholog open chromatin status differences between species and tissues could be affected by the amount of time that the animal had been was awake, in addition to species and tissue differences. Furthermore, an animal's sex has been shown to be associated with the activity of both brain [62, 63] and liver [64] enhancers. Although all of our datasets with multiple biological replicates had both males and females, the number of male and female replicates differed between datasets. We hope that our conservative definitions of clade-specific, species-specific, and tissue-specific OCRs prevented these OCRs from being sex-specific OCRs. However, our conservative definitions of clade-specific, species-specific, and tissue-specific OCRs limited our power to compare different models on these OCRs.

Furthermore, while our CNN provided accurate predictions of open chromatin conservation, using a CNN for our machine learning model has limitations. CNNs require inputs of a fixed size; this prevented us from accounting for differences in peak length between OCRs and would make using CNNs in future work incorporating long-range interactions difficult. CNNs also require extensive hyper-parameter tuning, and their performance can be sensitive to the random seed used in initialization. It is possible that, with more extensive hyper-parameter tuning or a different random seed, we would have been able to train models with better performance for some of the negative sets whose models had poor performance for our criteria or to obtain models trained on only mouse sequences with comparably good performance to the multi-species models. While multiple Bayesian optimization methods exist for automating much of the hyper-parameter tuning process [65-67], these methods often require extensive compute time that is not available to many researchers. SVMs do not have CNNs' input size limits, have only a few hyper-parameters to tune, and have been shown to work well on related tasks [68-70], but their prediction time can be slow because their kernels need to be computed for every DNA sequence, which could make using SVMs for predicting open chromatin conservation of hundreds of thousands of OCRs in each of dozens of species intractable. In addition, CNNs continue to be less directly interpretable than methods with user-defined features that cannot account for complex combinatorial relationships between sequence patterns involved in open chromatin, even though many advances have been made to improve the interpretability of CNNs [3, 4, 71]. Interpreting models for open chromatin conservation prediction could reveal the mechanisms through which enhancer orthologs have lost activity over evolutionary history, such as losses in TF motifs and changes in DNA shape.

<u>Potential Extensions of Our Method</u>

There are many ways to extend our approach for open chromatin conservation prediction that have the potential to both improve our accuracy and expand the space in which we can make predictions. Since our models trained on multiple negative sets work well, training models on combinations of negatives, while requiring more training time, may improve performance. In addition, training a model on a few species with open chromatin data using genome-wide negatives to predict OCRs genome-wide in species without open chromatin data would require substantial additional training time but may improve accuracy and would enable us to predict open chromatin in regions whose orthologs are not open in any of the species for which we have data. While some machine learning models have been successfully trained to predict open chromatin genome-wide [72, 73], such models have not yet been applied to predicting open chromatin conservation across species. Likewise, training a model that includes TF protein sequences and, if available, TF expression, could enable models to learn when differences between species in TF sequence or expression might be associated with differences in open chromatin. Finally, modifying our models to predict continuous open chromatin signal across species would enable us to not only predict changes in the existence of OCRs but also in their strength. A previous study trained CNNs to predict continuous open chromatin signal across species [74], suggesting that accomplishing this task might be feasible, but such models' ability to accurately predict changes in open chromatin between species has yet to be systematically evaluated. In fact, any extension to our approach would need to be evaluated for its ability to predict lineage- and tissue-specific open chromatin, and, given that some of our models trained with widely-used negative sets such as dinucleotide-shuffled sequences did not meet all of our evaluation criteria, direct application of some existing methods to predicting open chromatin conservation may not initially be successful.

**SUPPLEMENTAL MATERIALS AND METHODS**

Assaying Open Chromatin in Mouse Liver

We performed ATAC-seq experiments on two 10-week-old heterozygous Pvalb-2A-Cre mice (B6.Cg-Pvalb$^{tm1.1(cre)Aibs}$/J; Jackson Stock No: 012358) [75], one male (Replicate 1 in **Supplemental Figure 8a**) and one female (Replicate 2 in **Supplemental Figure 8a**). We euthanized the mice by isoflurane and decapitation. We quickly dissected fresh liver tissue and extracted nuclei by 30 strokes of Dounce homogenization with the loose pestle (0.005 in. clearance) in 5mL of cold lysis buffer [76]. We filtered the nuclei suspensions through a 70µm cell strainer, pelleted them by centrifugation at 2,000 x g for 10 minutes, resuspended them in water, and filtered them a final time through a 40µm cell strainer. We stained sample aliquots with DAPI (Invitrogen #D1206) and quantified nuclei concentrations using a manual hemocytometer under a fluorescent microscope. We then input approximately 50,000 nuclei into a 50µL ATAC-seq tagmentation reaction as described in [76] and [77]. We amplified the resulting libraries to 1/3 qPCR saturation, and fragment length distributions estimated by the Agilent TapeStation System showed high-quality ATAC-seq periodicity. We paired-end sequenced the samples on the Illumina NovaSeq 6000 System through Novogene services. We obtained 165,337,124 reads from the male mouse and 225,752,264 reads from the female mouse.

Identifying Brain and Liver OCRs

We used open chromatin data from four species: *Homo sapiens* [59, 60, 78], *Macaca Mulatta* [26], *Mus musculus* [25, 58], and *Rattus norvegicus* [26]. For human brain OCRs, we used NeuN+ primary motor cortex (4 biological replicates), putamen (4 biological replicates), and nucleus accumbens (1 biological replicate) ATAC-seq data from GSE96949 [59] and caudate and putamen DNase hypersensitivity data (2 biological replicates) from ENCODE [60]. For macaque brain OCRs, we used orofacial motor cortex (2 biological replicates), hand motor cortex (2 biological replicates), caudate (2 biological replicates), putamen (2 biological replicates), and nucleus accumbens (1 biological replicate) ATAC-seq data from our previous study [26]. For macaque liver OCRs, we used liver ATAC-seq data (1 biological replicate) from data we previously generated [26]. For mouse brain OCRs, we used cortex and striatum ATAC-seq data from seven-week-old and twelve-week-old mice from our previous study [58] (2 biological replicates each). For mouse liver OCRs, we used the mouse liver ATAC-seq data that we generated as well as mouse liver ATAC-seq data from CNP0000198 [25] (4 biological replicates). For rat brain OCRs, we used primary motor cortex (3 biological replicates) and striatum data (2 biological replicates) from our previous study [26]. For rat liver OCRs, we used liver ATAC-seq data (2 biological replicates) from our previous study [26]. For each dataset, we combined reads from technical replicates. In addition, we identified Laurasiatheria-specific liver OCRs and non-OCRs using cow and pig liver ATAC-seq data (2 biological replicates each) from the FAANG Consoritum [79].

We processed DNase hypersensitivity data by using the Kundaje Lab open chromatin pipeline [80] to map reads to hg38 [81], filter reads, call peaks, evaluate which peaks are reproducible, and remove peaks overlapping the ENCODE black list [82]. We used the default settings for the pipeline. We downloaded human brain DNase hypersensitivity data from the caudate nucleus and the putamen from the ENCODE portal [83]. Since the caudate nucleus and the putamen are both parts of the striatum but came from different people, we treated them as biological replicates. The final set of peaks was the larger set of the peaks that were reproducible according to the Irreproducible Discovery Rate (IDR) [84] across biological replicates and the peaks that were reproducible according to the IDR across pooled pseudo-replicates (the "optimal set").

We processed the mouse brain ATAC-seq data using the Kundaje Lab open chromatin pipeline [80] and the mouse liver, human, macaque, and rat ATAC-seq data as well as the cow and pig ATAC-seq data we used for identifying Laurasiatheria-specific OCRs and non-OCRs using the ENCODE ATAC-seq pipeline [85]. For the mouse brain ATAC-seq data, we began with the filtered bam files from data we previously generated [58] and used the default parameters for the remainder of the pipeline. For the other ATAC-seq data, we used the default parameters except for "atac.multimapping" : 0, "atac.cap_num_peak" : 300000, "atac.smooth_win" : 150, "atac.enable_idr" : true, and "atac.idr_thresh" : 0.1; these parameter modifications enabled the parameters for read filtering, peak calling, and calculating the IDR to be the same as those used for the mouse brain data. We mapped the human data to hg38 [81], the macaque data to rheMac8 [86], the mouse data to mm10 [87], the rat data to rn6 [88], the cow data to NCBI assembly Btau_5.0.1 [89], and the pig data to susScr3 [90]. For the mouse liver ATAC-seq data from CNP0000198 [25], we treated the two female and two male samples as four biological replicates. The final set of peaks for datasets with multiple biological replicates was the larger set of the peaks that were reproducible according to the IDR [84] across biological replicates and the peaks that were reproducible according to IDR across pooled pseudo-replicates (the “optimal set”); the final set of peaks for datasets with only 1 biological replicate was the peaks that were reproducible according to IDR across self-pseudo-replicates.

We then used the percentage of mapped reads, number of filtered reads, periodicity, TSS enrichment, number of IDR reproducible peaks, rescue ratio, and self-consistency ratio analyses generated by the pipelines [76, 91] to evaluate data quality. We found that most of the samples were high-quality. However, we excluded the second macaque nucleus accumbens biological replicate because it had only about sixteen million filtered reads and poor periodicity and because the two replicates had rescue ratio 4.01 and self-consistency ratio 2.04. We also excluded the second macaque liver replicate because it had only about two million filtered reads and poor periodicity and because the two replicates had self-consistency ratio 3.53. In addition, we excluded the first rat liver biological replicate it had only 35,593 reproducible peaks according to the IDR across self-pseudo-replicates in spite of having over sixty-eight million filtered reads. As a result, for macaque nucleus accumbens and liver, we used the peaks from the first biological replicate that were reproducible according to the IDR across self-pseudo-replicates, and, for rat liver, we used the “optimal set” from running the ENCODE ATAC-seq pipeline on only biological replicates 2 and 3.

Constructing Additional Negative Sets

*Flanking Regions*

We constructed the flanking region negative set by using bedtools [92, 93] to identify the subset of regions flanking mouse brain OCRs that are not OCRs. Specifically, we first identified the 500bp flanking regions of each of our OCRs +/- 500bp; we required a 500bp separation between flanks and OCRs to ensure that we would not have false positives in our negative set due to poorly defined peak boundaries. We then removed all flanking regions that overlapped any open chromatin peaks called from the pooled set of reads across biological replicates from cortex or striatum from either age [80, 85]; we used these peaks instead of the subset of such peaks that we defined as OCRs because non-reproducible peaks have the potential to be enhancers, and we wanted to limit the number of false negatives in our negative set. For each remaining flanking region, we used its underlying sequence and that sequence’s reverse complement. Thus, although there could be up to two negatives for every positive, our negative:positive training data ratio was approximately 1.65:1 (**Supplemental Table 20**).

*OCRs from Other Tissues*

We constructed the OCRs from other tissues negative set by identifying OCRs in non-cortex and non-striatum tissues that do not overlap our brain OCRs. We first used the ENCODE ATAC-seq pipeline [85] with the same parameters that we used for the brain samples to process all of the ATAC-seq data from tissues that do not overlap cortex and striatum from the mouse ENCODE post-natal samples [94] and from CNP0000198 [25]. We then used the same quality control metrics that we used for selecting datasets to include as OCRs to evaluate the quality of these datasets and removed those that were low-quality. The mouse ENCODE datasets that we included were from liver, intestine, and cerebellum. (We did not use this for our liver positive set because it came from an embryonic sample, and the other datasets were from adults.) The datasets from CNP0000198 [25] that we included were from female abdominal fat, female adrenal gland, female kidney, male kidney, female liver, male liver, female lung, male lung, female pancreas, male small intestine, male spleen, male stomach, female thymus, and male thymus. (For the purposes of creating this negative set, male and female samples were processed separately for all tissues, including liver.) We obtained the union of the IDR "optimal set" peaks across all of these datasets as well as our mouse liver data and used bedtools subtract with the -A option [92] to remove those peaks that overlapped open any chromatin peaks called from the pooled set of replicates from cortex or striatum from either age. For each filtered peak, we used the sequence underlying its summit +/- 250bp and that sequence's reverse complement. Our negative:positive training data ratio was approximately 19.78:1 (**Supplemental Table 20**).

*G/C- and Repeat-Matched Regions*

We identified G/C- and repeat-matched regions for our OCRs using a combination of R packages and bedtools [92]. We first created a repeat-masked mm10 genome by running forgeMaskedBSgenomeDataPkg from the BSgenome R package [95] on mm10 [87] with masks downloaded from the UCSC Genome Browser [96]. We then ran genNullSeqs from the gkmSVM R package [69, 97] on the sequences of the brain OCR peak summits +/- 250bp and our masked mouse genome with default parameters except for the following: length_match_tol=0.00, which ensures that all of our sequences are 500bp; nMaxTrials=100, which allows for more attempts to find G/C- and repeat-matched regions than the default; and xfold=10 for the larger G/C- and repeat-matched region negative set and =2 for the smaller G/C- and repeat-matched region negative set. Although we allowed for more trials, getNullSeqs found fewer G/C- and repeat-matched regions than we had requested. After generating these regions, we used bedtools subtract [92] with the -A option to remove any regions that overlapped any open chromatin peaks called from the pooled set of replicates from cortex or striatum from either age. For each filtered G/C- and repeat-matched region, we used its underlying sequence and that sequence's reverse complement. As a result, for the larger G/C- and repeat-matched negative set, the negative:positive training data ratio was approximately 8.15:1, and, for the smaller G/C- and repeat-matched negative set, the negative:positive training data ratio was approximately 1.64:1 (**Supplemental Table 20**).

*Dinucleotide-Shuffled OCRs*

We obtained dinucleotide shuffled OCRs by running the MEME suite's [98] fasta-shuffle-letters on the sequences of our brain OCR peak summits +/- 250bp. We used the default parameters except for -kmer 2, which enabled us to preserve dinucleotide frequencies, and -copies 10, which enabled us to generate a negative set that was ten times larger than our positive set. We used every shuffled sequence and its reverse complement. Thus, the negative:positive training data ratio was exactly 10:1 (**Supplemental Table 20**).

<u>Constructing Training, Validation, and Test Sets</u>

For the models trained using only mouse sequences, we divided the positives and negatives (except for the dinucleotide-shuffled OCRs) into training, validation, and test sets based on chromosomes to ensure that there would be no overlap between the sets. For the dinucleotide-shuffled OCRs negatives, we put them into the set that corresponded to the positive example from which they were constructed. Our training set consisted of regions on mm10 chromosomes 3-7, 10-19, and X. Our validation set that we used for developing our positive and negative set definitions (for example, validation set performance was used to determine that we should use orthologs of loose OCRs instead of OCRs for our novel negative set), early-stopping, and hyper-parameter tuning consisted of regions on mm10 chromosomes 8-9. Our test set consisted of mm10 chromosomes 1-2. All presented evaluations on mouse genomic regions, including those on types of regions not used in model training, were done on only regions from mm10 chromosomes 1-2.

For the models and model evaluations using sequences from non-mouse species, we divided sequences into training, validation, and test sets based on the chromosomes to which their mouse orthologs mapped. In other words, we mapped such regions to mm10 using halLiftover [99] with the Zoonomia version 1 Cactus alignment [27, 100] followed by HALPER [101] with parameters -max_frac 2.0, -min_len 50, and -protect_dist 5 and put them into the training set if their mm10 orthologs were on chromosomes 3-7, 10-19, or X; put them into the validation set if their mm10 orthologs were on chromosomes 8-9; put them into the test set if their mm10 orthologs were on chromosomes 1-2; and excluded them if their orthologs were elsewhere in mm10 or if they had no orthologs. Although many non-mouse regions were excluded from evaluation, because some OCRs have high sequence conservation, not accounting for the location of mouse orthologs when constructing training, validation, and test sets could lead to test set sequences that are almost identical to training set sequences [78, 102].

Calibrating Machine Learning Models

Because the machine learning models trained with some of the negative sets had high sensitivity and low specificity, we re-calibrated them with the training data from our novel negative set. More specifically, we first made predictions with the model we wanted to re-calibrate on the training data from the positive set and our novel negative set. We next trained a logistic regression to use the model's predictions as features to predict the real open chromatin status for these training examples. We then used the logistic regression to make predictions on the test set. The training and prediction was done using Scikit-learn [103].

Evaluating the Relationship between OCR Ortholog Open Chromatin Status and Genome Quality

To evaluate the relationship between predicted OCR ortholog open chromatin status and genome quality, we computed the correlation between the mean and standard deviation of predicted OCR ortholog open chromatin status and the Glires' genome assemblies' scaffold and contig N50's. We obtained the scaffold and contig N50's from NCBI [28, 36] and computed the log base ten of each of them. We computed the correlations for predictions from multi-species brain and liver models, using brain and liver OCR orthologs, respectively. We also determined the relative association of phylogenetic distance and genome quality with predictions by fitting generalized linear models of mean and standard deviation of predicted OCR ortholog open chromatin status as a combination of divergence from mouse and scaffold or contig N50. In addition to comparing the effect sizes for the generalized linear models, we also computed the p-values on the coefficients and multiplied them by four to do a Bonferroni correction.

Interpreting Deep Learning Models

We interpreted the deep learning models by computing the importance of every nucleotide in each true positive example in the validation set and then using these importance values to construct motifs. We computed the importance of every nucleotide in every true positive example in the validation set using deepLIFT, which calculates the extent to which each input contributes to the prediction relative to a reference [3]. We used the deepLIFT version 0.5.5-theano with the Rescale rule scores from the sequence layer with the target of the final convolutional layer, where our reference was a sequence of

N's. We also used an extension to deepLIFT, also with the Rescale rule, to compute the "hypothetical scores" for each nucleotide at each position for each sequence, which can be thought of as the preference of the model for observing each nucleotide at each position in the sequence [4].

We combined the scores and hypothetical scores using the TF-MoDISco method to construct "TF-MoDISco Motifs" [4]. TF-MoDISco first identifies frequently occurring sequence patterns with high deepLIFT scores within the sequences of each OCR (called seqlets), next computes a similarity matrix between all seqlets, and then uses the similarity matrix to cluster the seqlets into nonredundant motifs. We used the following settings for TF-MoDIsco: seqlet FDR threshold = 0.2; gapped k-mer settings for similarity computation k-mer length = 8, number of gaps = 1, and number of mismatches = 0; and final motif width = 50. We visualized our TF-MoDISco motifs from TF-MoDISco using the aggregated hypothetical scores of the seqlets supporting each motif. We created position frequency matrices from TF-MoDISco motifs by averaging the one-hot-encoded sequences at all of the seqlet coordinates belonging to the motifs and compared them to known motifs by running TomTom [104] on them with the *Mus musculus* motifs from CIS-BP [105].

<u>Comparing Liver Open Chromatin Predictions to H3K27ac ChIP-seq</u>:

To compare our predicted open chromatin conservation to H3K27ac ChIP-seq conservation, we first used halLiftover [99] with the Zoonomia version 1 Cactus alignment [27, 100] followed by HALPER [101] with settings -max_frac 2.0, -min_len 50, and -protect_dist 5 to identify orthologs of all mouse, rat, and macaque liver OCRs in all Zoonomia species except for *Manis tricuspis*, which was not in the Cactus alignment. We next used our multi-species liver model to predict the liver open chromatin statuses of the orthologs and orthologs' reverse complements in all of the species except for *Galeopterus variegatus*, *Hippopotamus amphibius*, *Monodon monoceros*, *Platanista gangetica*, and *Procavia capensis*, which we excluded due to challenges converting between chromosome naming conventions. Then, for each OCR ortholog, we set the prediction to be the average between the prediction for the ortholog and the prediction for its reverse complement. After that, we obtained the liver H3K27ac ChIP-seq regions in each species and those regions' orthologs in other species with liver H3K27ac ChIP-seq data as well as whether those orthologs overlapped H3K27ac ChIP-seq regions from https://www.ebi.ac.uk/research/flicek/publications/FOG15 [35]. We mapped the rat ChIP-seq regions from rn5 to rn6 and the macaque ChIP-seq regions from rheMac2 to rheMac8 using liftOver [106]. We finally filtered the liver OCRs by removing those that did not overlap H3K27ac ChIP-seq regions in the same species.

When evaluating the relationship between liver open chromatin predictions and liver H3K27ac ChIP-seq conservation, we considered all Boreoeutheria with liver H3K27ac ChIP-seq except for *Chlorocebus sabaeus* because the H3K27ac ChIP-seq reads were mapped to *Chlorocebus pygerythrus* instead of *Chlorocebus sabaeus* [35]. For each of mouse, rat, and macaque, we considered H3K27ac ChIP-seq to be conserved if there was a liver H3K27ac ChIP-seq region overlapping the ortholog and to be non-conserved if the ortholog did not have an overlapping H3K27ac ChIP-seq region; we did not include any species for which either the H3K27ac ChIP-seq data or our overlapping OCRs had no ortholog. Then, for each combination of species for which we had liver open chromatin data and species with liver H3K27ac ChIP-seq data, we compared the multi-species liver model predictions for the orthologs with conserved H3K27ac ChIP-seq to those for the orthologs with non-conserved H3K27ac ChIP-seq using a Wilcoxon rank-sum test; we did a Bonferroni correction by multiplying all p-values by twenty-nine, which was the number of tests we did. We also found that median of our predictions for the orthologs with conserved H3K27ac ChIP-seq was higher than the median of our predictions for the orthologs with non-conserved H3K27ac ChIP-seq.

#### Obtaining and Visualizing Signal Tracks

We obtained the signal tracks used in **Figure 4** using the pooled replicates fold-change bigwigs from the data processing pipelines. For the H3K27ac ChIP-seq data, we downloaded the mouse and macaque H3K27ac ChIP-seq data from E-MTAB-2633 [35] and reprocessed it using the AQUAS Transcription Factor and Histone ChIP-Seq processing pipeline [107] with default parameters, mapping reads to mm10 and rheMac8, respectively. We evaluated the data quality of each biological replicate based on the percentage of mapped reads, number of filtered reads, NSC, RSC, number of IDR reproducible peaks, rescue ratio, and self-consistency ratio analyses generated by the pipelines and found that all four biological replicates from each species were high-quality. We created visualizations for these figures using the New WashU Epigenome Browser [108].

#### Identifying Genes with Rodent-Specific Expression:

We obtained the median transcripts per million (TPM) of each human gene for each species from https://www.ebi.ac.uk/research/flicek/publications/FOG20 [109]. We next removed all genes that did not have a median TPM value in every placental mammal. We then used limma version 3.42.2 [110] to identify genes that have higher expression in rodents relative to other placental mammals. We used Benjamini-Hochberg to adjust p-values [111] and considered a gene to have rodent-specific expression if it had adjusted p-value < 0.05 and log2 fold-change < -1 (Larger fold change corresponded to higher expression in non-rodent placental mammals.).

#### Identifying OCRs with Predicted Rodent-Specific Open Chromatin:

To identify Rodent-specific open chromatin regions, we used the pre-filtered OCR ortholog predictions that we used when comparing our predictions to H3K27ac ChIP-seq. We filtered our predictions by using bedtools [92] to remove all rat OCR orthologs for rat OCRs for which the mouse ortholog overlapped a mouse liver OCR and all macaque OCR orthologs for macaque OCRs for which the mouse ortholog overlapped a mouse liver OCR or the rat ortholog overlapped a rat liver OCR. In addition, we removed all OCRs for which there was not a usable ortholog in at least 1 Rodent and at least 1 non-Rodent. Finally, for each remaining OCR, we ran a Wilcoxon rank-sum test to compare the predictions for its orthologs in Rodents to those for its orthologs in non-Rodents and did a Bonferroni correction by multiplying the p-values by the number of remaining OCRs. We considered an OCR to have predicted Rodent-specific open chromatin if the median activity in Rodents was greater than the median activity outside of Rodents and the Bonferroni-corrected p-value was less than 0.05.

### SUPPLEMENTAL FIGURE CAPTIONS

**Supplemental Figure 1: Additional Lineage-Specific OCR Accuracy Evaluations for Different Negative Sets**

**a)** To demonstrate that our models can accurately predict whether sequence differences between species are associated with open chromatin differences, in addition to the evaluations described in previous work [68], we evaluated our performance on (1) clade-specific losses in open chromatin for a clade not used in model training, (2) species-specific losses in open chromatin for a species not used in model training, and (3) tissue-specific open chromatin. Since such regions often comprise a minority of open chromatin region

(OCR) orthologs, models could obtain good overall performance while obtaining poor performance on such regions.

**b)** Illustration of different negative sets.

**c)** Performance of models trained on all negative sets on test sets from negative sets used in model training.

**d)** Performance of models trained on all negative sets on mouse brain test set OCRs whose orthologs in at least one other species are not brain OCRs and mouse brain test set non-OCRs whose orthologs in at least one other species are brain OCRs.

**e)** Test set performance of models trained on all negative sets on mouse brain OCRs whose orthologs in rat are not brain OCRs and mouse brain non-OCRs whose orthologs in rat are brain OCRs.

**f)** Test set performance of models trained on all negative sets on Glires-specific brain OCRs and Glires-specific brain non-OCRs.

**g)** Performance of models trained on all negative sets on macaque brain OCRs whose orthologs in mouse are not brain OCRs and are on test set chromosomes and macaque brain non-OCRs whose orthologs in mouse are brain OCRs and are on test set chromosomes.

**h)** Performance of models trained on all negative sets on human brain OCRs whose orthologs in mouse are not brain OCRs and are on test set chromosomes and human brain non-OCRs whose orthologs in mouse are brain OCRs and are on test set chromosomes.

**i)** Performance of models trained on all negative sets on rat brain OCRs whose orthologs in mouse are not brain OCRs and are on test set chromosomes and rat brain non-OCRs whose orthologs in mouse are brain OCRs and are on test set chromosomes.

**j)** Performance of models trained on all negative sets on Euarchonta-specific brain OCRs and Euarchonta-specific brain non-OCRs whose mouse orthologs are on test set chromosomes.

Animal silhouettes were obtained from PhyloPic [112]. AUC stands for area under the receiver operating characteristic curve, AUPRC stands for area under the precision-recall curve, Rep. stands for repeat, Dinuc.-Shuf. stands for dinucleotide-shuffled, and Orths. stands for orthologs. For evaluations with more positives than negatives, we reported the area under the negative predictive value (NPV)-specificity (Spec.) curve instead of the area under the precision-recall curve.

**Supplemental Figure 2: Violin Plots for Lineage-Specific and Tissue-Specific OCR Accuracy Evaluation in Human**

Comparison of PhastCons [2] and PhyloP [1] scores to three different machine learning models' predictions for brain OCRs with conserved open chromatin across mouse and human, human brain OCRs whose mouse orthologs are closed in brain, human brain non-OCRs whose mouse orthologs are open in brain, human brain OCRs that are closed in liver, human brain OCRs that are open in liver (centered on brain peak summits), and human liver OCRs that are closed in brain. +'s indicate that values should be large, and -'s indicate that values should be small. Conservation scores were generated from the mm10-based placental mammals alignment [113, 114] and averaged over 500bp centered on peak summits, where mouse peak summits were used for OCRs conserved between mouse and human and OCRs in mouse whose human orthologs are closed, and mouse orthologs of human peak summits were used for other evaluations. All machine learning model predictions were made using human sequences, where the human sequences for OCRs conserved between mouse and human and OCRs in mouse whose human orthologs are not OCRs were centered on human orthologs of mouse peak summits, and human peak summits were used for other evaluations. Animal silhouettes were obtained from PhyloPic [112]. *'s indicate the species from which sequences were obtained for making predictions. Dinuc.-shuf. stands for dinucleotide-shuffled, and Orths. stands for orthologs.

**Supplemental Figure 3: Violin Plots for Lineage-Specific and Tissue-Specific OCR Accuracy Evaluation in Rat**

**a)** Comparison of PhastCons [2] and PhyloP [1] scores to three different machine learning models' predictions for brain OCRs with conserved open chromatin across mouse and rat, rat brain OCRs whose mouse orthologs are closed in brain, rat brain non-OCRs whose mouse orthologs are open in brain, rat brain OCRs that are closed in liver, rat brain OCRs that are open in liver (centered on brain peak summits), and rat liver OCRs that are closed in brain.

**b)** Comparison of PhastCons [2] and PhyloP [1] scores to two different machine learning models' predictions for liver OCRs with conserved open chromatin across mouse and rat, rat liver OCRs whose mouse orthologs are closed in liver, rat liver non-OCRs whose mouse orthologs are open in liver, rat liver OCRs that are closed in brain, rat liver OCRs that are open in brain (centered on liver peak summits), and rat brain OCRs that are closed in liver.

+'s indicate that values should be large, and -'s indicate that values should be small. Conservation scores were generated from the mm10-based placental mammals alignment [113, 114] and averaged over 500bp centered on peak summits, where mouse peak summits were used for OCRs conserved between mouse and rat and OCRs in mouse whose rat orthologs are closed, and mouse orthologs of rat peak summits were used for other evaluations. All machine learning model predictions were made using rat sequences, where the rat sequences for OCRs conserved between mouse and rat and OCRs in mouse whose rat orthologs are closed were centered on rat orthologs of mouse peak summits, and rat peak summits were used for other evaluations. Animal silhouettes were obtained from PhyloPic [112]. *'s indicate the species from which sequences were obtained for making predictions. Dinuc.-shuf. stands for dinucleotide-shuffled, and Orths. stands for orthologs.

**Supplemental Figure 4: Performance of Brain Model Trained with Smaller G/C- and Repeat-Matched Negatives before and after Calibration**

**a)** Test set performance on Glires-specific brain open chromatin regions (OCRs) and Glires-specific brain non-OCRs before and after calibration with training set positives and non-OCR orthologs of OCR negatives. We reported the negative predictive value (NPV) instead of the precision because there are more positives than negatives in this evaluation.

**b)** Performance on Euarchonta-specific brain OCRs and Euarchonta-specific brain non-OCRs whose mouse orthologs are on test set chromosomes before and after calibration with training set positives and non-OCR orthologs of OCRs.

Animal silhouettes were obtained from PhyloPic [112].

**Supplemental Figure 5: Additional Tissue-Specific OCR Accuracy Evaluations – Performance of Brain Models on Liver OCRs**

**a)** We made test set predictions with machine learning models trained on different negative sets on brain open chromatin regions (OCRs) that do not overlap liver OCRs, brain OCRs that overlap liver OCRs, liver OCRs that do not overlap brain OCRs, and negative sets. p-Values were computed with a Wilcoxon rank-sum test and a Bonferroni correction across all negative sets.

**b)** We evaluated the performance of the brain models on the positives that are also OCRs in liver and on the negatives that are OCRs in liver but not OCRs in brain. We did this for such positives and negatives on the mouse test set chromosomes and in the macaque and rat whose orthologs are on mouse test set chromosomes.

**c)** We evaluated the performance of the brain model trained with the smaller G/C- and repeat-matched negatives on the positives that are also OCRs in liver and on negatives that are OCRs in liver but not OCRs in brain before and after calibration with the training set positives and non-OCR orthologs of OCRs. We did this for such positives and negatives on the mouse test set chromosomes and in the macaque and rat whose orthologs are on mouse test set chromosomes.

The moues silhouette was obtained from PhyloPic [112]. AUC stands for area under the receiver operating characteristic curve, AUPRC stands for area under the precision-recall curve, Rep. stands for repeat, and Orths. stands for orthologs.

**Supplemental Figure 6: TF-MoDISco Motifs from Brain Models Trained with Different Negative Sets**

Each table contains motifs from TF-MoDISco; transcription factors (TFs) whose motifs match the TF-MoDISco motifs with TomTom q-value < 0.05 ordered from most to least significant TomTom p-value [104], where red TFs are those whose motifs are considered important and gold TFs are those whose motifs' depletions are considered important; and number of supporting seqlets for each motif.

**a)** TF-MoDISco motifs for brain model with flanking region negative set.

**b)** TF-MoDISco motifs for brain model with open chromatin regions (OCRs) from other tissues negative set.

**c)** TF-MoDISco motifs for brain model with larger G/C- and repeat-matched regions negative set.

**d)** TF-MoDISco motifs for brain model with smaller G/C- and repeat-matched regions negative set.

**e)** TF-MoDISco motifs for brain model with dinucleotide-shuffled OCRs negative set.

**f)** TF-MoDISco motifs for brain model with non-OCR orthologs of OCRs negative set.

**Supplemental Figure 7: Phylogeny-Matching Correlations Evaluation on Different Negative Sets**

**a)** Divergence from mouse versus mean of the brain open chromatin region (OCR) ortholog open chromatin status predictions across each Glires species from the model trained on each negative set. The curves are the best fit exponential functions of the form $y = ae^{bx}$. The dotted lines are the average predictions across test set negatives.

**b)** Divergence from mouse versus standard deviation (Std. Dev.) of the brain OCR ortholog open chromatin status predictions across each of the Glires from the model trained on each negative set. The curves are the best fit exponential functions of the form $y = c/(1 + ae^{-bx})$.

Animal silhouettes were obtained from PhyloPic [112]. Rep. stands for repeat, Orths. stands for orthologs, and MYA stands for millions of years ago.

**Supplemental Figure 8: Performance of Mouse Liver Models**

**a)** Periodicity plots for each of the biological replicates from our new mouse liver ATAC-seq data.

**b)** Test set performance of mouse liver models on the full test set, subset whose ortholog in at least one other species has different liver open chromatin status, subset whose ortholog in rat has different liver open chromatin status, macaque liver open and closed chromatin regions whose mouse orthologs have different liver open chromatin statuses, rat liver open and closed chromatin regions whose mouse orthologs have different liver open chromatin statuses, subset for which the liver open or closed chromatin is Glires-specific, and Euarchonta-specific liver open chromatin regions (OCRs) and non-OCRs. For the full test set, macaque liver open and closed chromatin regions whose mouse orthologs have different liver open chromatin status, and Euarchonta-specific liver open and closed chromatin regions, we reported the area under the negative predictive value (NPV)-specificity (Spec.) curve because these evaluations had more positives than negatives.

**c)** Test set performance of mouse liver models on liver OCRs that overlap brain OCRs and on negatives that are brain OCRs that do not overlap liver OCRs from mouse, macaque, and rat.

**d)** Divergence from mouse versus mean predictions across mouse liver OCR orthologs in Glires. The curve is the best fit exponential function of the form $y = ae^{bx}$. The dotted line is the average prediction across test set negatives. MYA stands for millions of years ago.

**e)** Divergence from mouse versus standard deviation (Std. Dev.) of predictions across mouse liver OCR orthologs in Glires. The curve is the best fit exponential function of the form $y = c/(1 + ae^{-bx})$. MYA stands for millions of years ago.

**f)** TF-MoDISco motifs for the mouse liver model; transcription factors (TFs) whose motifs match the TF-MoDISco motifs with TomTom q-value < 0.05 ordered from most to least significant TomTom p-value [104], where red TFs are those whose motifs are considered important and gold TFs are those whose motifs' depletions are considered important; and number of supporting seqlets for each motif.

Animal silhouettes were obtained from PhyloPic [112]. AUC stands for area under the receiver operating characteristic curve, and AUPRC stands for area under the precision-recall curve.

**Supplemental Figure 9: Additional Evaluations from Multi-Species Brain and Liver Models**

**a)** Divergence from mouse versus standard deviation (Std. Dev.) of multi-species brain model predictions across mouse brain open chromatin region (OCR) orthologs in Glires. The red curve is the best fit exponential function of the form $y = c/(1 + ae^{-bx})$. MYA stands for millions of years ago.

**b)** Divergence from mouse versus Std. Dev. of multi-species liver model predictions across mouse liver OCR orthologs in Glires. The red curve is the best fit exponential function of the form $y = c/(1 + ae^{-bx})$. MYA stands for millions of years ago.

**c)** TF-MoDISco motifs for multi-species brain model; transcription factors (TFs) whose motifs match the TF-MoDISco motifs with TomTom q-value < 0.05 ordered from most to least significant TomTom p-value [104], where red TFs are those whose motifs are considered important and gold TFs are those whose motifs' depletions are considered important; and number of supporting seqlets for each motif.

**d)** TF-MoDISco motifs for multi-species brain model; TFs whose motifs match the TF-MoDISco motifs with TomTom q-value < 0.05 ordered from most to least significant TomTom p-value [104], where red TFs are those whose motifs are considered important and gold TFs are those whose motifs' depletions are considered important; and number of supporting seqlets for each motif.

Animal silhouettes were obtained from PhyloPic [112].

**Supplemental Figure 10: Genome Quality versus Open Chromatin Status Predictions in Glires**

**a)** log base ten of scaffold and contig N50's of each of the Glires versus mean brain open chromatin region (OCR) ortholog open chromatin status prediction across each of the Glires.

**b)** log base ten of scaffold and contig N50's of each of the Glires versus mean liver OCR ortholog open chromatin status prediction across each of the Glires.

**c)** log base ten of scaffold and contig N50's of each of the Glires versus standard deviation (Std. Dev.) of brain OCR ortholog open chromatin status predictions across each of the Glires.

**d)** log base ten of scaffold and contig N50's of each of the Glires versus Std. Dev. of liver OCR ortholog open chromatin status predictions across each of the Glires.

**Supplemental Figure 11: Rodent-Specific Liver OCR near Gene with Rodent-Specific Liver Expression**

Example of Rodent-specific liver open chromatin region (OCR) near Atp9a, a gene with Rodent-specific expression in liver that has been associated with bile production in rats. Most animal silhouettes were obtained from PhyloPic [112].

**Supplemental Figure 12: Additional Predicted Lineage-Specific OCR Clusters Associated with Neuron Firing, Neuron Activity, and Liver Regeneration**

**a)** Additional predicted Murinae-specific brain open chromatin region (OCR) cluster (cluster 27) with significant overlap with mouse enhancers associated with neuron firing.

**b)** Predicted Hystricognathi-specific brain non-OCR cluster (cluster 11) and Muroidea and Pecora-specific non-OCR cluster (cluster 48) with significant overlap with human enhancers associated with neuron activity.

**c)** We clustered the liver OCRs where the features were the liver predictions in each Boreoeutherian species from Zoonomia and then identified clusters whose regions had significant overlap with regions associated with mouse liver regeneration. These clusters were a Murinae-specific OCR cluster (cluster 29) and two Muroidea-specific OCR clusters (cluster 36 and cluster 100).

Animal silhouettes were obtained from PhyloPic [112].

### SUPPLEMENTAL TABLES

**Supplemental Table 1: Mouse Sequence Brain Model Sensitivity on Glires-Specific Brain OCRs and Non-OCRs before and after Calibration**

| Training Negative Set | Uncalibrated Model Sensitivity | Calibrated Model Sensitivity |
|---|---|---|
| Flanking Regions | 0.84 | 0.74 |
| OCRs in Other Tissues | 0.73 | 0.71 |
| Large G/C- and Repeat-Matched | 0.71 | 0.70 |
| Small G/C- and Repeat-Matched | 0.86 | 0.72 |
| Dinucleotide-Shuffled OCRs | 0.83 | 0.67 |
| Non-OCR Orths. of OCRs | 0.74 | 0.77 |

**Supplemental Table 2: Mouse Sequence Brain Model Specificity on Glires-Specific Brain OCRs and Non-OCRs before and after Calibration**

| Training Negative Set | Uncalibrated Model Specificity | Calibrated Model Specificity |
|---|---|---|
| Flanking Regions | 0.81 | 0.90 |
| OCRs in Other Tissues | 0.79 | 0.82 |
| Large G/C- and Repeat-Matched | 0.95 | 0.96 |
| Small G/C- and Repeat-Matched | 0.74 | 0.88 |
| Dinucleotide-Shuffled OCRs | 0.61 | 0.79 |
| Non-OCR Orths. of OCRs | 0.92 | 0.92 |

**Supplemental Table 3: Mouse Sequence Brain Model Precision on Glires-Specific Brain OCRs and Non-OCRs before and after Calibration**

| Training Negative Set | Uncalibrated Model Precision | Calibrated Model Precision |
|---|---|---|
| Flanking Regions | 0.89 | 0.92 |
| OCRs in Other Tissues | 0.86 | 0.87 |
| Large G/C- and Repeat-Matched | 0.96 | 0.97 |
| Small G/C- and Repeat-Matched | 0.85 | 0.91 |
| Dinucleotide-Shuffled OCRs | 0.79 | 0.85 |
| Non-OCR Orths. of OCRs | 0.94 | 0.94 |

**Supplemental Table 4: Mouse Sequence Brain Model Sensitivity on Euarchonta-Specific Brain OCRs and Non-OCRs before and after Calibration**

| Training Negative Set | Uncalibrated Model Sensitivity | Calibrated Model Sensitivity |
|---|---|---|
| Flanking Regions | 0.74 | 0.65 |
| OCRs in Other Tissues | 0.75 | 0.72 |
| Large G/C- and Repeat-Matched | 0.60 | 0.57 |
| Small G/C- and Repeat-Matched | 0.84 | 0.69 |
| Dinucleotide-Shuffled OCRs | 0.68 | 0.51 |
| Non-OCR Orths. of OCRs | 0.60 | 0.61 |

**Supplemental Table 5: Mouse Sequence Brain Model Specificity on Euarchonta-Specific Brain OCRs and Non-OCRs before and after Calibration**

| Training Negative Set | Uncalibrated Model Specificity | Calibrated Model Specificity |
|---|---|---|
| Flanking Regions | 0.64 | 0.80 |
| OCRs in Other Tissues | 0.66 | 0.68 |
| Large G/C- and Repeat-Matched | 0.84 | 0.86 |
| Small G/C- and Repeat-Matched | 0.55 | 0.73 |
| Dinucleotide-Shuffled OCRs | 0.54 | 0.72 |
| Non-OCR Orths. of OCRs | 0.85 | 0.83 |

**Supplemental Table 6: Mouse Sequence Brain Model Precision on Euarchonta-Specific Brain OCRs and Non-OCRs before and after Calibration**

| Training Negative Set | Uncalibrated Model Precision | Calibrated Model Precision |
|---|---|---|
| Flanking Regions | 0.55 | 0.65 |
| OCRs in Other Tissues | 0.56 | 0.57 |
| Large G/C- and Repeat-Matched | 0.69 | 0.70 |
| Small G/C- and Repeat-Matched | 0.52 | 0.60 |
| Dinucleotide-Shuffled OCRs | 0.46 | 0.51 |
| Non-OCR Orths. of OCRs | 0.70 | 0.68 |

**Supplemental Table 7: PhastCons, PhyloP, and Predictions for Mouse Brain OCRs with Conserved versus Non-Conserved Open Chromatin Status**

| Species with Mouse OCR Orthologs | PhastCons Scores | PhyloP Scores | Predictions from Brain Model Trained on Mouse | Predictions from Multi-Species Brain Model |
|---|---|---|---|---|
| Macaque | $4.82 \times 10^{-18}$ | $4.12 \times 10^{-10}$ | $8.06 \times 10^{-129}$ | $2.39 \times 10^{-143}$ |
| Human | $6.13 \times 10^{-9}$ | $1.05 \times 10^{-6}$ | $4.19 \times 10^{-68}$ | $4.59 \times 10^{-80}$ |
| Rat | $1.50 \times 10^{-16}$ | $7.18 \times 10^{-14}$ | $6.73 \times 10^{-123}$ | $7.14 \times 10^{-149}$ |

**Supplemental Table 8: PhastCons, PhyloP, and Predictions for Mouse Liver OCRs with Conserved versus Non-Conserved Open Chromatin Status**

| Species with Mouse OCR Orthologs | PhastCons Scores | PhyloP Scores | Predictions from Liver Model Trained on Mouse | Predictions from Multi-Species Liver Model |
|---|---|---|---|---|
| Macaque | $1.69 \times 10^{-5}$ | > 1 | $1.34 \times 10^{-182}$ | $1.75 \times 10^{-228}$ |
| Rat | $1.30 \times 10^{-4}$ | > 1 | $4.46 \times 10^{-157}$ | $4.19 \times 10^{-213}$ |

**Supplemental Table 9: OCR Predictions by Mouse Sequence Models on Other Species' Orthologs versus Conservation Scores**

| Tissue | Species | Conservation Score Type | Open Chromatin Conserved | Open Chromatin Not Conserved |
|---|---|---|---|---|
| Brain | Human | PhastCons | $5.81 \times 10^{-4}$ | $4.26 \times 10^{-1}$ |
| Brain | Human | PhyloP | $7.09 \times 10^{-5}$ | $9.77 \times 10^{-2}$ |
| Brain | Rat | PhastCons | $1.24 \times 10^{-2}$ | $4.80 \times 10^{-3}$ |
| Brain | Rat | PhyloP | $9.50 \times 10^{-3}$ | $3.60 \times 10^{-8}$ |
| Liver | Rat | PhastCons | $5.85 \times 10^{-6}$ | $9.30 \times 10^{-2}$ |
| Liver | Rat | PhyloP | $6.98 \times 10^{-7}$ | $2.54 \times 10^{-2}$ |

**Supplemental Table 10: Mouse Sequence Brain Model Sensitivity on Liver OCRs before and after Calibration**

| Training Negative Set | Uncalibrated Model Sensitivity | Calibrated Model Sensitivity |
|---|---|---|
| Flanking Regions | 0.90 | 0.85 |
| OCRs in Other Tissues | 0.71 | 0.69 |
| Large G/C- and Repeat-Matched | 0.80 | 0.79 |
| Small G/C- and Repeat-Matched | 0.91 | 0.83 |
| Dinucleotide-Shuffled OCRs | 0.86 | 0.79 |
| Non-OCR Orths. of OCRs | 0.85 | 0.86 |

**Supplemental Table 11: Mouse Sequence Brain Model Specificity on Liver OCRs before and after Calibration**

| Training Negative Set | Uncalibrated Model Specificity | Calibrated Model Specificity |
|---|---|---|
| Flanking Regions | 0.79 | 0.89 |
| OCRs in Other Tissues | 0.97 | 0.98 |
| Large G/C- and Repeat-Matched | 0.95 | 0.95 |
| Small G/C- and Repeat-Matched | 0.75 | 0.89 |
| Dinucleotide-Shuffled OCRs | 0.53 | 0.69 |

| | | |
|---|---|---|
| Non-OCR Orths. of OCRs | 0.93 | 0.92 |

**Supplemental Table 12: Mouse Sequence Brain Model Precision on Liver OCRs before and after Calibration**

| Training Negative Set | Uncalibrated Model Precision | Calibrated Model Precision |
|---|---|---|
| Flanking Regions | 0.56 | 0.70 |
| OCRs in Other Tissues | 0.88 | 0.90 |
| Large G/C- and Repeat-Matched | 0.82 | 0.83 |
| Small G/C- and Repeat-Matched | 0.53 | 0.70 |
| Dinucleotide-Shuffled OCRs | 0.36 | 0.44 |
| Non-OCR Orths. of OCRs | 0.78 | 0.77 |

**Supplemental Table 13: Wilcoxon Rank-Sum Test p-Values for Differences in Predicted Open Chromatin in Liver OCR-H3K27ac ChIP-seq Region Orthologs with and without H3K27ac ChIP-seq (All Directions Match Expectations)**

| Species with H3K27ac | Mouse OCRs | Rat OCRs | Macaque OCRs |
|---|---|---|---|
| *Mus musculus* | N/A | $9.45 \times 10^{-101}$ | $4.04 \times 10^{-78}$ |
| *Callithrix jacchus* | $1.51 \times 10^{-46}$ | $6.61 \times 10^{-50}$ | $3.03 \times 10^{-106}$ |
| *Macaca mulatta* | $8.86 \times 10^{-58}$ | $8.05 \times 10^{-55}$ | N/A |
| *Felis catus* | $1.30 \times 10^{-39}$ | $9.36 \times 10^{-32}$ | $4.39 \times 10^{-87}$ |
| *Homo sapiens* | $1.05 \times 10^{-50}$ | $8.40 \times 10^{-47}$ | $3.32 \times 10^{-71}$ |
| *Bos taurus* | $2.00 \times 10^{-49}$ | $3.42 \times 10^{-51}$ | $1.73 \times 10^{-94}$ |
| *Canis lupus familiaris* | $9.10 \times 10^{-39}$ | $5.76 \times 10^{-33}$ | $3.10 \times 10^{-105}$ |
| *Oryctolagus cuniculus* | $2.42 \times 10^{-29}$ | $3.59 \times 10^{-38}$ | $9.75 \times 10^{-59}$ |
| *Heterocephalus glaber* | $2.09 \times 10^{-11}$ | N/A | N/A |
| *Cavia porcellus* | $8.06 \times 10^{-14}$ | N/A | N/A |
| *Sus scrofa* | $3.13 \times 10^{-34}$ | $8.57 \times 10^{-27}$ | $8.15 \times 10^{-64}$ |
| *Rattus norvegicus* | $1.74 \times 10^{-104}$ | N/A | $6.82 \times 10^{-63}$ |

**Supplemental Table 14: GLM Results – Mean Prediction as Function of Divergence from Mouse and log10(Scaffold/Contig N50)**

| Tissue | Genome Quality Metric | Distance from Mouse Coefficient | Distance from Mouse p-Value | Genome Quality Coefficient | Genome Quality p-Value |
|---|---|---|---|---|---|
| Brain | Scaffold N50 | $-2.78 \times 10^{-3}$ | $2.80 \times 10^{-22}$ | $1.88 \times 10^{-10}$ | $5.88 \times 10^{-1}$ |
| Brain | Contig N50 | $-2.82 \times 10^{-3}$ | $6.09 \times 10^{-26}$ | $7.39 \times 10^{-10}$ | $1.74 \times 10^{-1}$ |
| Liver | Scaffold N50 | $-2.22 \times 10^{-3}$ | $4.89 \times 10^{-8}$ | $3.25 \times 10^{-10}$ | $8.44 \times 10^{-1}$ |
| Liver | Contig N50 | $-2.35 \times 10^{-3}$ | $1.92 \times 10^{-9}$ | $1.77 \times 10^{-9}$ | 1.04 |

**Supplemental Table 15: GLM Results – Standard Deviation of Prediction as Function of Divergence from Mouse and log10(Scaffold/Contig N50)**

| Tissue | Genome Quality Metric | Distance from Mouse Coefficient | Distance from Mouse p-Value | Genome Quality Coefficient | Genome Quality p-Value |
|---|---|---|---|---|---|
| Brain | Scaffold N50 | $4.99 \times 10^{-4}$ | $1.03 \times 10^{-14}$ | $-1.60 \times 10^{-10}$ | $1.30 \times 10^{-4}$ |
| Brain | Contig N50 | $5.63 \times 10^{-4}$ | $9.87 \times 10^{-18}$ | $-8.31 \times 10^{-10}$ | $1.08 \times 10^{-3}$ |
| Liver | Scaffold N50 | $4.79 \times 10^{-4}$ | $3.98 \times 10^{-4}$ | $-1.47 \times 10^{-10}$ | $2.53 \times 10^{-1}$ |
| Liver | Contig N50 | $5.27 \times 10^{-4}$ | $4.55 \times 10^{-7}$ | $-8.11 \times 10^{-10}$ | $3.22 \times 10^{-1}$ |

**Supplemental Table 16: OCR Predictions by Multi-Species Models on Other Species' Orthologs versus Conservation Scores**

| Tissue | Species | Conservation Score Type | Brain Open Chromatin Conserved | Brain Open Chromatin Not Conserved |
|---|---|---|---|---|
| Brain | Human | PhastCons | $5.85 \times 10^{-6}$ | $9.30 \times 10^{-2}$ |
| Brain | Human | PhyloP | $6.98 \times 10^{-7}$ | $2.54 \times 10^{-2}$ |
| Brain | Rat | PhastCons | $7.05 \times 10^{-4}$ | $1.62 \times 10^{-9}$ |
| Brain | Rat | PhyloP | $5.95 \times 10^{-4}$ | $3.99 \times 10^{-12}$ |
| Liver | Rat | PhastCons | $5.00 \times 10^{-22}$ | $1.52 \times 10^{-11}$ |
| Liver | Rat | PhyloP | $1.88 \times 10^{-25}$ | $1.65 \times 10^{-17}$ |

**Supplemental Table 17: Significance of Overlap between Mouse Neuron Firing Enhancers (Bic Induces Neuron Firing, TTX Blocks Neuron Firing) and Brain Clusters Active in Mouse**

| Cluster | Bic-Specific Enhancers | Activity-Invariant Enhancers | TTX-Specific Enhancers |
|---|---|---|---|
| cluster 1 | > 1 | $8.54 \times 10^{-19}$ | > 1 |
| cluster 13 | > 1 | > 1 | > 1 |
| cluster 17 | > 1 | > 1 | > 1 |
| cluster 23 | > 1 | $5.27 \times 10^{-1}$ | > 1 |
| cluster 26 | > 1 | $9.47 \times 10^{-1}$ | > 1 |
| cluster 27 | $6.00 \times 10^{-3}$ | > 1 | > 1 |
| cluster 30 | $7.28 \times 10^{-1}$ | > 1 | > 1 |
| cluster 37 | > 1 | $6.97 \times 10^{-3}$ | > 1 |
| cluster 4 | > 1 | > 1 | > 1 |
| cluster 43 | $2.37 \times 10^{-3}$ | > 1 | > 1 |
| cluster 49 | > 1 | > 1 | > 1 |
| cluster 51 | > 1 | > 1 | > 1 |
| cluster 58 | > 1 | > 1 | > 1 |
| cluster 60 | > 1 | $9.10 \times 10^{-1}$ | > 1 |
| cluster 63 | > 1 | > 1 | > 1 |
| cluster 71 | > 1 | > 1 | > 1 |
| cluster 79 | $3.47 \times 10^{-1}$ | $6.60 \times 10^{-2}$ | > 1 |
| cluster 81 | > 1 | $1.92 \times 10^{-8}$ | > 1 |
| cluster 82 | > 1 | $5.94 \times 10^{-2}$ | > 1 |
| cluster 88 | > 1 | $3.83 \times 10^{-2}$ | > 1 |
| cluster 94 | > 1 | > 1 | > 1 |

**Supplemental Table 18: Significance of Overlap between Human Neuron Activity Up and Down Enhancers (Minutes/Hours: Time after KCl Exposure) and Brain Clusters Active in Human**

| Cluster | ↑, 15 Minutes | ↓, 15 Minutes | ↑, 2 Hours | ↓, 2 Hours |
|---|---|---|---|---|
| cluster 1 | 5.04 x $10^{-2}$ | 4.83 x $10^{-3}$ | > 1 | 2.97 x $10^{-3}$ |
| cluster 11 | > 1 | > 1 | 6.18 x $10^{-4}$ | > 1 |
| cluster 13 | > 1 | > 1 | > 1 | > 1 |
| cluster 21 | > 1 | > 1 | > 1 | > 1 |
| cluster 26 | > 1 | > 1 | > 1 | > 1 |
| cluster 29 | > 1 | > 1 | > 1 | > 1 |
| cluster 32 | > 1 | > 1 | > 1 | > 1 |
| cluster 41 | > 1 | > 1 | > 1 | > 1 |
| cluster 42 | > 1 | > 1 | > 1 | > 1 |
| cluster 48 | > 1 | > 1 | 5.84 x $10^{-3}$ | > 1 |
| cluster 5 | > 1 | > 1 | > 1 | > 1 |
| cluster 55 | > 1 | 9.70 x $10^{-1}$ | > 1 | > 1 |
| cluster 61 | > 1 | > 1 | > 1 | > 1 |
| cluster 67 | > 1 | > 1 | > 1 | > 1 |
| cluster 73 | > 1 | > 1 | > 1 | > 1 |
| cluster 74 | > 1 | > 1 | 4.52 x $10^{-2}$ | > 1 |
| cluster 77 | > 1 | > 1 | > 1 | > 1 |
| cluster 78 | > 1 | > 1 | > 1 | > 1 |
| cluster 81 | > 1 | > 1 | > 1 | > 1 |
| cluster 82 | > 1 | > 1 | > 1 | 2.03 x $10^{-2}$ |
| cluster 95 | > 1 | > 1 | > 1 | > 1 |
| cluster 96 | > 1 | > 1 | > 1 | > 1 |
| cluster 99 | > 1 | > 1 | > 1 | > 1 |

**Supplemental Table 19: Significance of Overlap between Mouse Liver Regeneration Enhancers (Wk.: Weeks into Hepatocyte Repopulation) and Liver Clusters Active in Mouse**

| Cluster | Wk. 1 ↑ vs. Ctl. | Wk. 1 ↓ vs. Ctl. | Wk. 4 ↑ vs. Ctl. | Wk. 4 ↓ vs. Ctl. | Wk. 4 ↑ vs. Wk. 1 | Wk. 4 ↓ vs. Wk. 1 |
|---|---|---|---|---|---|---|
| cluster 100 | > 1 | > 1 | 6.36 x $10^{-4}$ | > 1 | > 1 | > 1 |
| cluster 17 | > 1 | > 1 | > 1 | > 1 | > 1 | > 1 |
| cluster 18 | > 1 | > 1 | > 1 | > 1 | > 1 | > 1 |
| cluster 28 | 7.60 x $10^{-1}$ | > 1 | > 1 | > 1 | > 1 | > 1 |
| cluster 29 | > 1 | > 1 | 2.00 x $10^{-3}$ | > 1 | > 1 | > 1 |
| cluster 2 | 6.96 x $10^{-1}$ | > 1 | 2.43 x $10^{-1}$ | > 1 | > 1 | > 1 |
| cluster 31 | > 1 | > 1 | > 1 | > 1 | > 1 | > 1 |
| cluster 34 | > 1 | > 1 | > 1 | > 1 | > 1 | > 1 |
| cluster 36 | > 1 | > 1 | 9.29 x $10^{-3}$ | > 1 | > 1 | > 1 |
| cluster 39 | 3.31 x $10^{-1}$ | > 1 | > 1 | > 1 | > 1 | 6.60 x $10^{-3}$ |
| cluster 51 | > 1 | > 1 | > 1 | > 1 | > 1 | > 1 |

| | | | | | | |
|---|---|---|---|---|---|---|
| cluster 55 | > 1 | > 1 | > 1 | > 1 | > 1 | > 1 |
| cluster 59 | > 1 | > 1 | > 1 | > 1 | > 1 | > 1 |
| cluster 64 | > 1 | > 1 | > 1 | > 1 | > 1 | > 1 |
| cluster 69 | > 1 | > 1 | > 1 | > 1 | > 1 | > 1 |
| cluster 75 | > 1 | > 1 | > 1 | > 1 | > 1 | > 1 |
| cluster 76 | > 1 | > 1 | > 1 | > 1 | > 1 | > 1 |
| cluster 78 | > 1 | > 1 | > 1 | > 1 | > 1 | > 1 |
| cluster 83 | $4.34 \times 10^{-2}$ | > 1 | > 1 | > 1 | > 1 | > 1 |
| cluster 84 | > 1 | > 1 | > 1 | > 1 | > 1 | > 1 |
| cluster 8 | > 1 | > 1 | > 1 | > 1 | > 1 | > 1 |
| cluster 9 | > 1 | > 1 | > 1 | > 1 | > 1 | > 1 |
| cluster 93 | > 1 | > 1 | > 1 | > 1 | > 1 | > 1 |
| cluster 94 | > 1 | > 1 | > 1 | > 1 | > 1 | > 1 |

**Supplemental Table 20: Number of Positives and Negatives Used for Training, Tuning, and Testing Each Model**

| Genomes Used in Training | Tissue | Negative Set | Positives (Training, Validation, Test) | Negatives (Training, Validation, Test) | Negatives:Positives (Training, Validation, Test) |
|---|---|---|---|---|---|
| mm10 | Brain | Flanking Regions | 21594, 2416, 4576 | 35640, 4018, 7440 | 1.65:1, 1.66:1, 1.63:1 |
| mm10 | Brain | OCRs in Other Tissues | 21594, 2416, 4576 | 427174, 70504, 82172 | 19.78:1, 29.18:1, 17.96:1 |
| mm10 | Brain | Large G/C- and Repeat-Matched | 21594, 2416, 4576 | 175912, 23880, 32008 | 8.15:1, 9.88:1, 6.99:1 |
| mm10 | Brain | Small G/C- and Repeat-Matched | 21594, 2416, 4576 | 35358, 4776, 6654 | 1.64:1, 1.98:1, 1.45:1 |
| mm10 | Brain | Dinucleotide-Shuffled Enhs. | 21594, 2416, 4576 | 215940, 24160, 45760 | 10:1, 10:1, 10:1 |
| mm10 | Brain | Non-OCR Orths. of OCRs | 21594, 2416, 4576 | 25086, 3456, 4694 | 1.16:1, 1.43:1, 1.03:1 |
| mm10 | Liver | Non-OCR Orths. of OCRs | 32498, 4032, 7752 | 22890, 2994, 4434 | 1:1.42, 1:1.35, 1:1.75 |
| mm10, hg38, rheMac8, rn6 | Brain | Non-OCR Orths. Of OCRs | 74688, 9036, 15266 | 111206, 14650, 19688 | 1.49:1, 1.62:1, 1.29:1 |
| mm10, rheMac8, rn6 | Liver | Non-OCR Orths. of OCRs | 81886, 10428, 17688 | 67278, 8680, 14544 | 1:1.22, 1:1.20, 1:1.22 |
