## Supplementary figures and images for "Inferring mammalian tissue-specific regulatory conservation by predicting tissue-specific differences in open chromatin"

### SupplementalFigure1

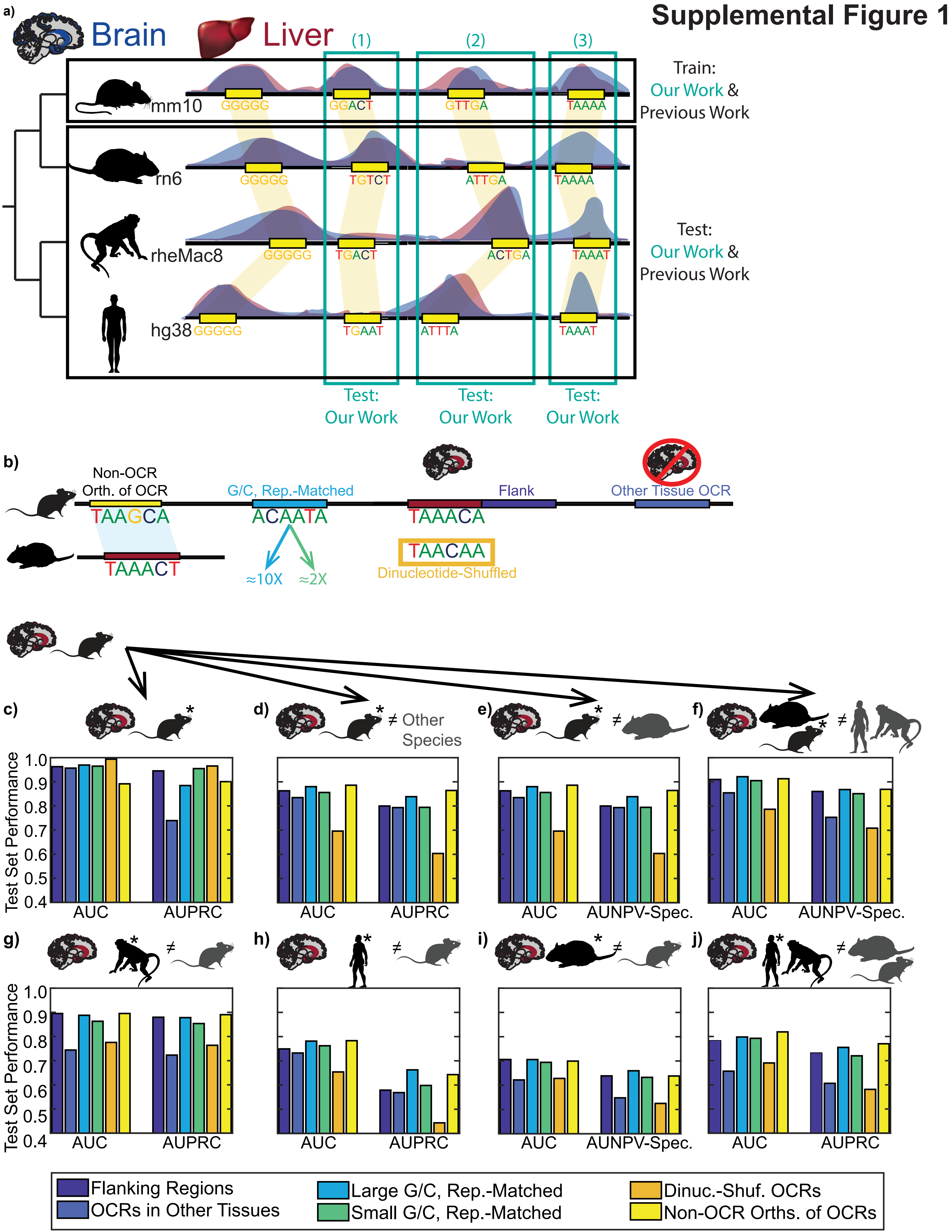

### SupplementalFigure2

#### Supplemental Figure 2

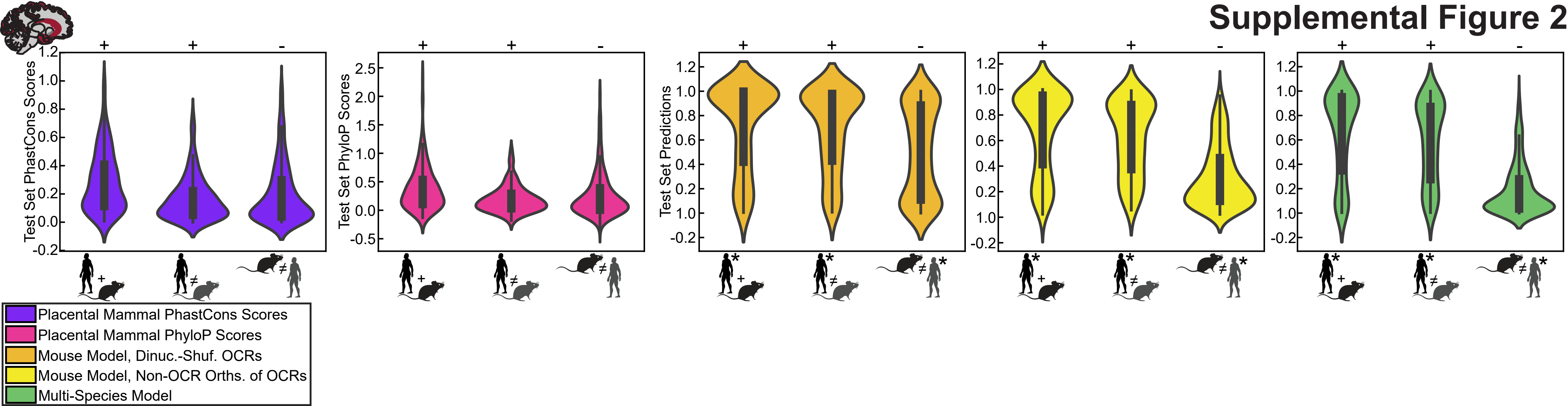

### SupplementalFigure3

#### Supplemental Figure 3

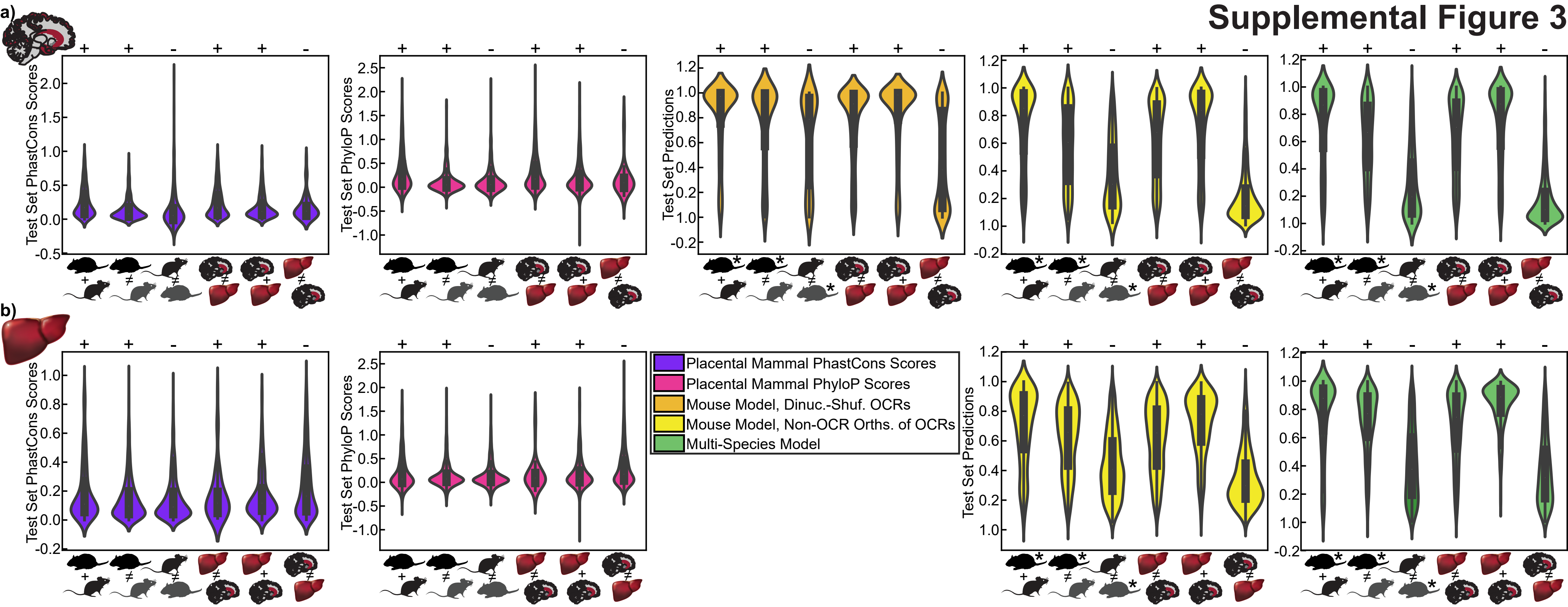

### SupplementalFigure4

##### Supplemental Figure 4

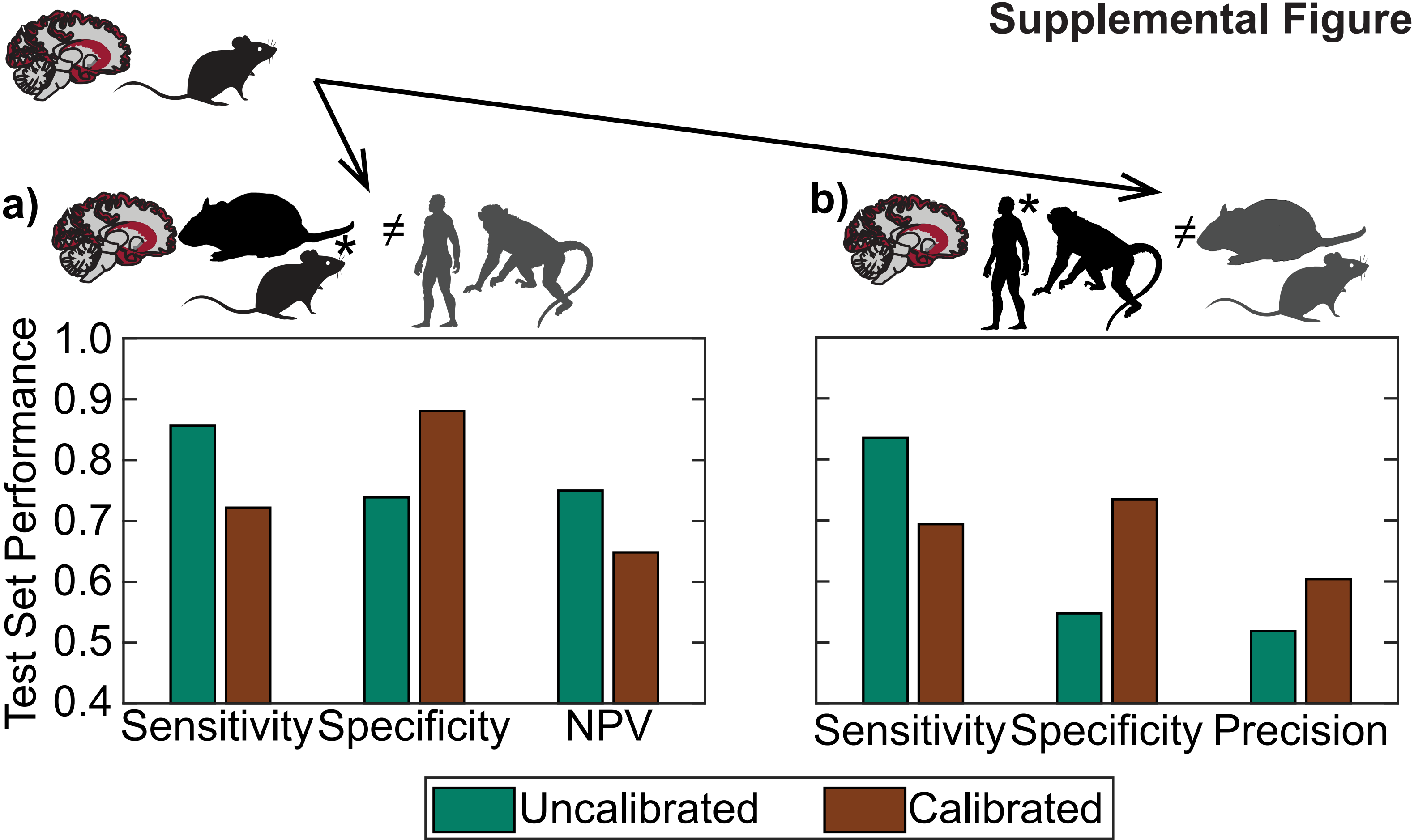

### SupplementalFigure5

#### Supplemental Figure 5

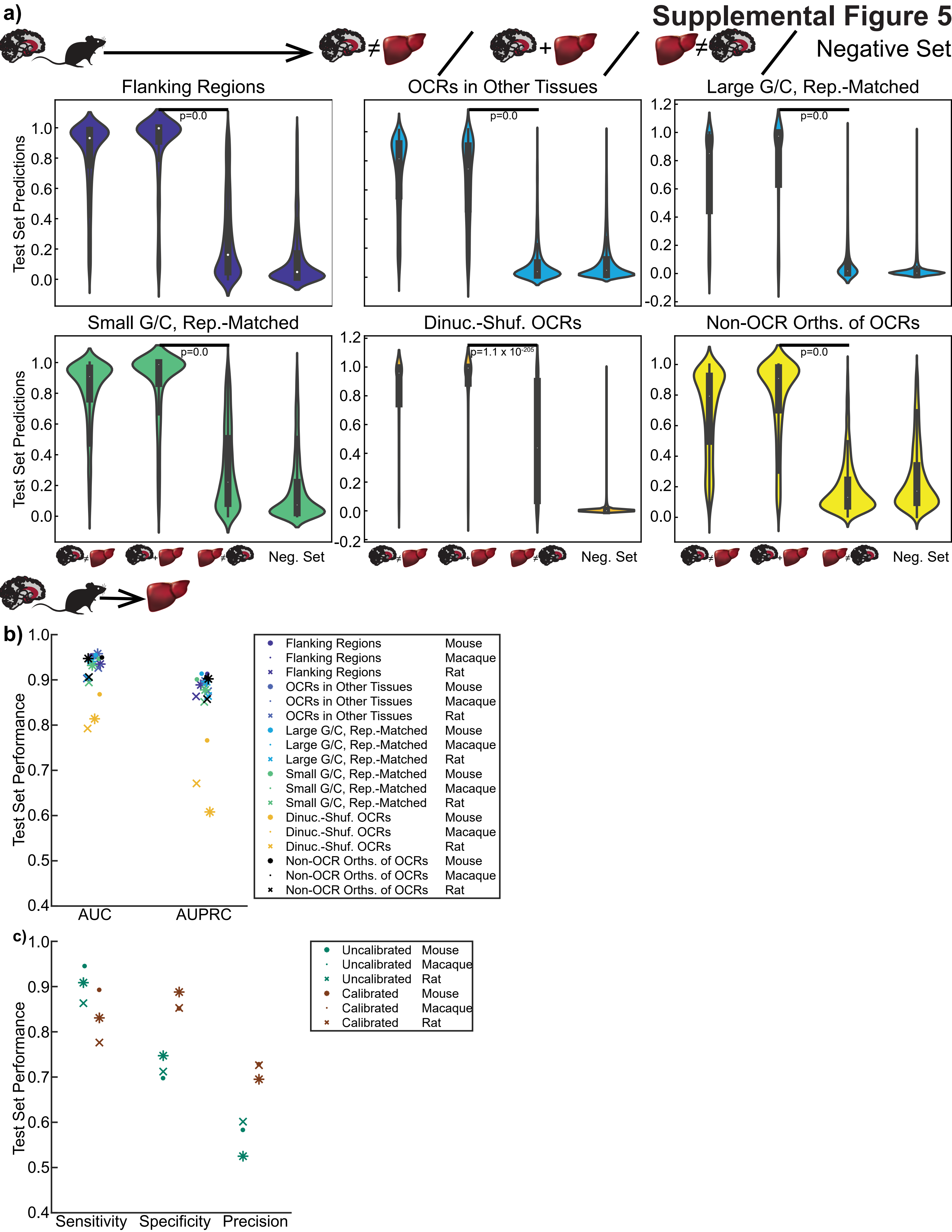

### SupplementalFigure7

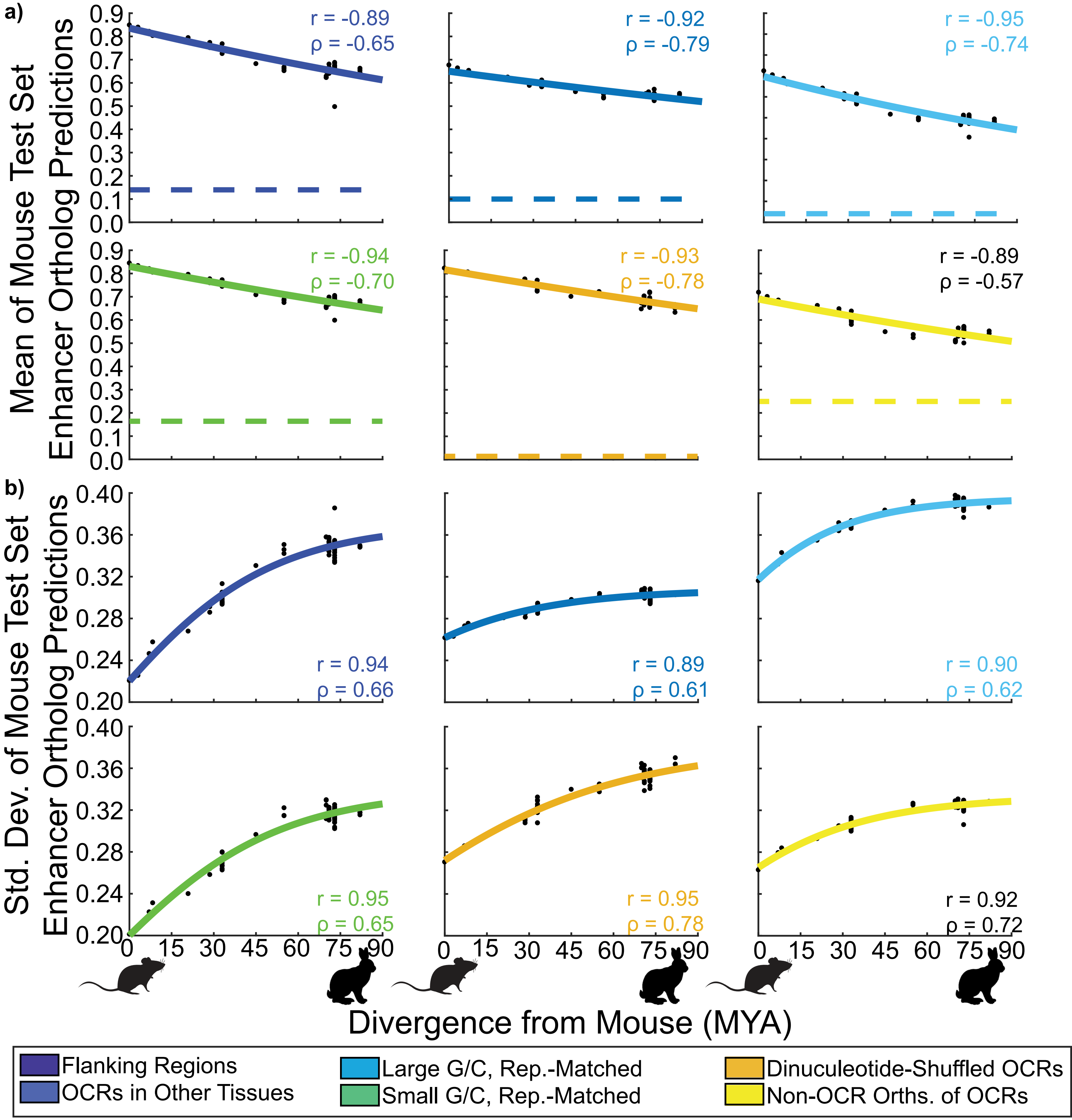


**Supplemental Figure 7**

### SupplementalFigure10

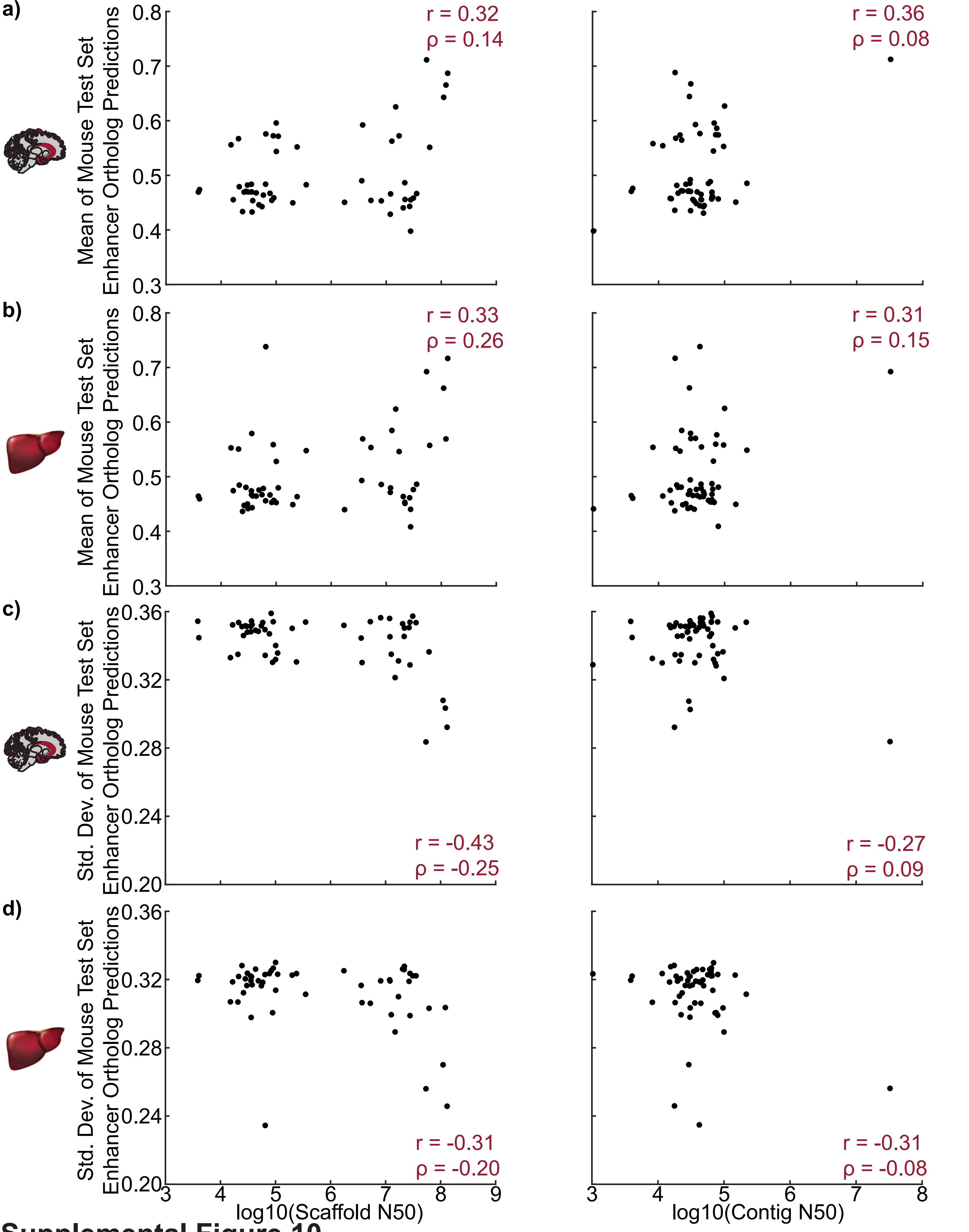


**Supplemental Figure 10**

### SupplementalFigure11

**Supplemetnal Figure 11**

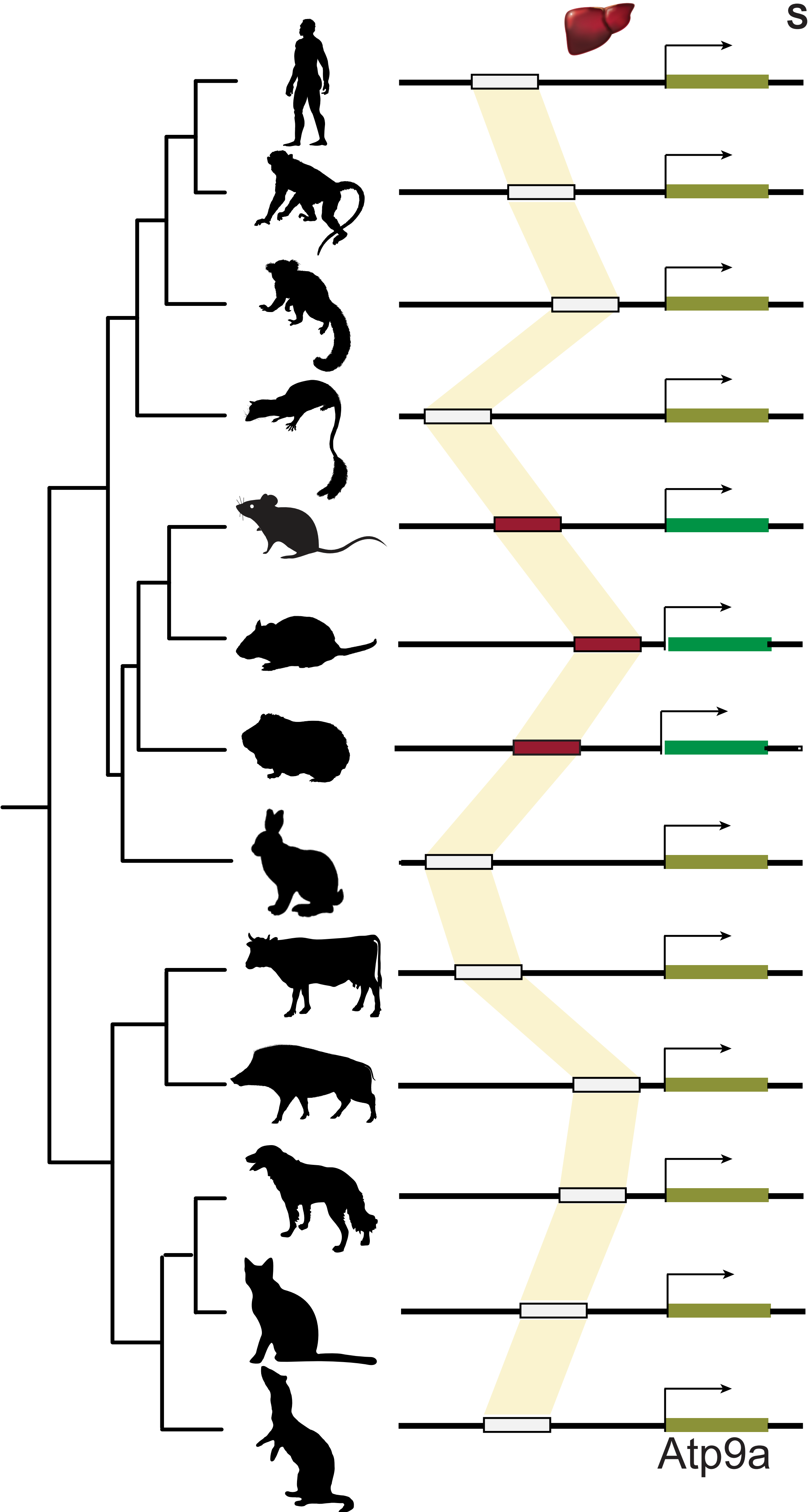

### SupplementalFigure12

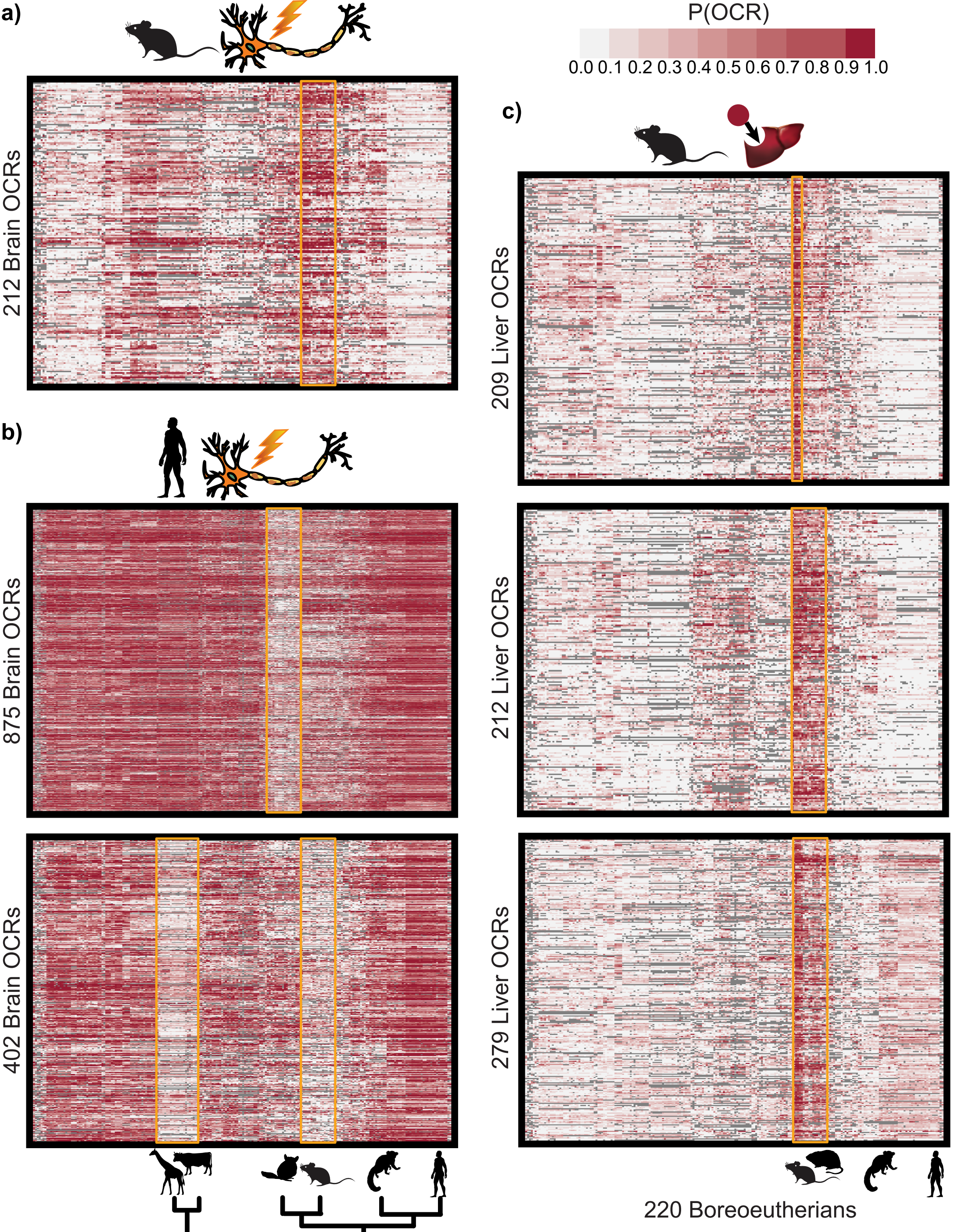


#### Supplemental Figure 12
