## SupplementalFigure6 for "Inferring mammalian tissue-specific regulatory conservation by predicting tissue-specific differences in open chromatin"

a)

| TF-MoDISco Motif | TFs with Similar Motif | Seqlets |
|---|---|---|
| 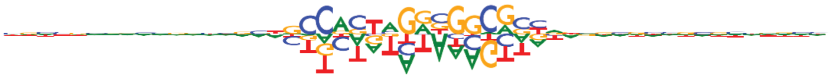 | Ctcf, Ctcfl | 267 |
| 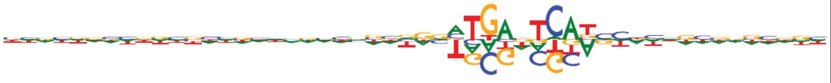 | Fos, Smarcc1, Fosb, Jund | 143 |
| 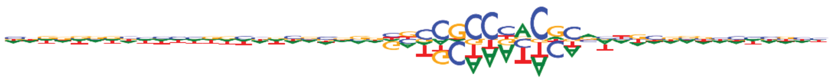 | Egr2 | 137 |
| 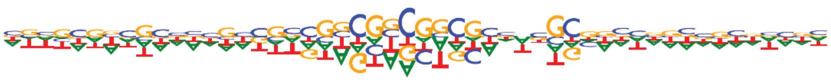 | | 97 |
| 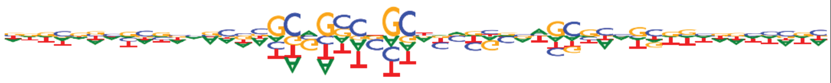 | | 77 |
| 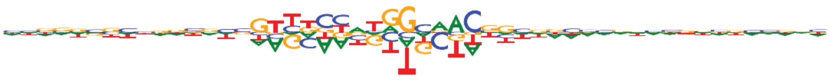 | Rfx1, Arid2, Rfx2, Rfx4, Rfx7, Rfx5, Rfx3 | 35 |
| 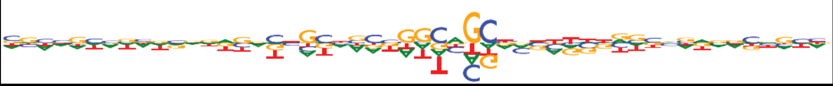 | | 31 |
| 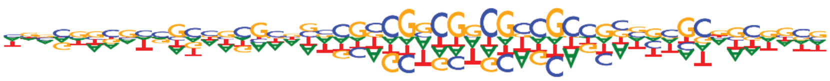 | | 28 |

b)

| TF-MoDISco Motif | TFs with Similar Motif | Seqlets |
|---|---|---|
| 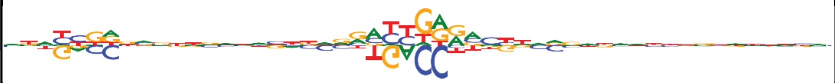 | Rxra, Hnf4g, Nr4a3, Ppara, Nr1h2 | 340 |
| 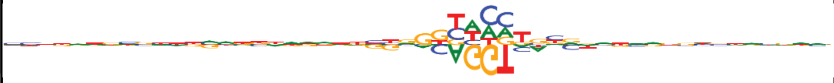 | | 71 |
| 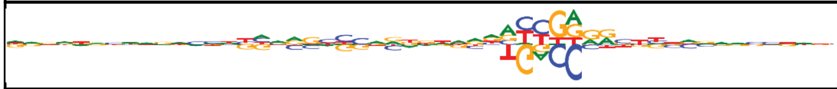 | Nr4a3, Rxra, Nr4a2, Nr4a1, Ppara, Nr5a1 | 69 |
| 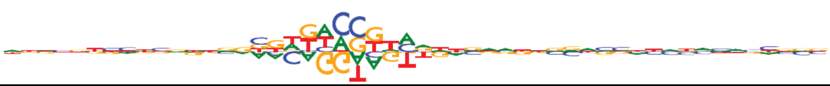 | Erg, Fev, Bcl11a, Elk3, Elk1, Fli1, Erf, Etv3, Elk4, Etv6, Etv5, Elf4, Gm4881, Etv1, Etv4, Gm5454, Elf2, Etv2, Ets1, Spfi1, Zkscan5, Elf3, Ehf | 39 |
| 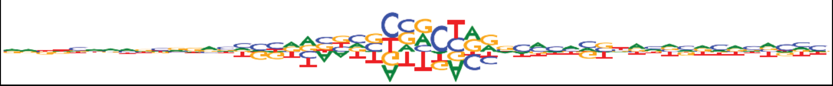 | Foxc1, Foxc2, Foxl1, Foxp1, Foxb1, Foxp2, Foxj2, Foxo3, Foxj3, Foxd1, Foxk1, Foxg1, Foxf2, Foxo4, Foxd2, ENSMUSG00000090020, Foxn3, Foxo6, Foxa2, Gm5294, Foxp4, Tbpl2 | 44 |
| 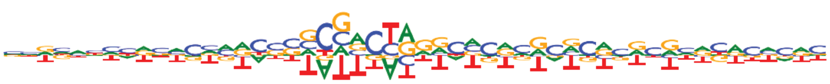 | | 35 |
| 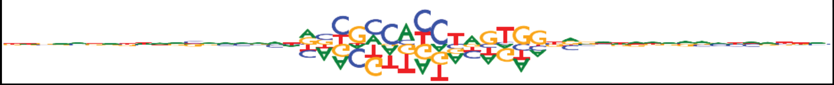 | Ctcf, Ctcfl | 169 |
| 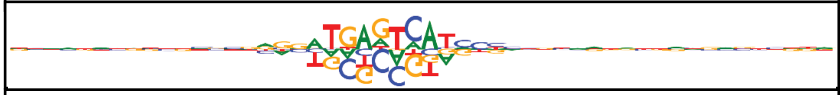 | Fos, Jund, Smarcc1, Fosb | 143 |
| 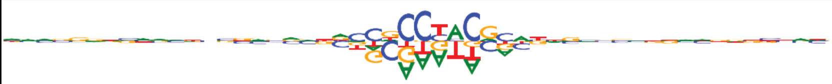 | | 101 |
| 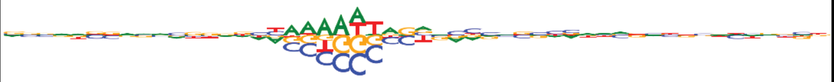 | Mef2c, Mef2a, Mef2d, Mef2b | 36 |
| 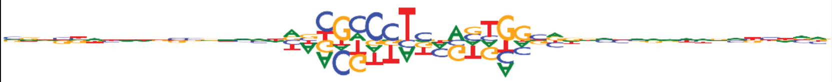 | Ctcf, Ctcfl, Twist1 | 29 |
| 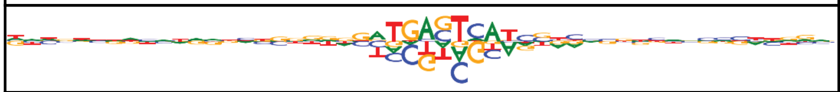 | | 24 |
| 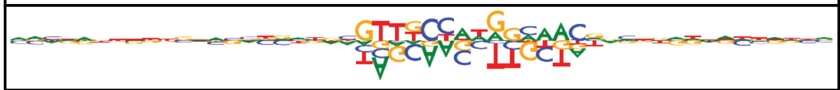 | Rfx1, Rfx2, Rfx4, Arid2, Rfx7, Rfx5, Rfx3 | 38 |

**Supplemental Figure 6**

c)

| TF-MoDISco Motif | TFs with Similar Motif | Seqlets |
|---|---|---|
| 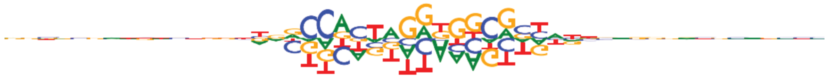 | Ctcf, Ctcfl | 275 |
| 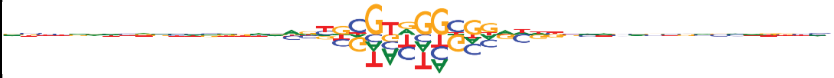 | Egr3, Egr2, Bcl6 | 190 |
| 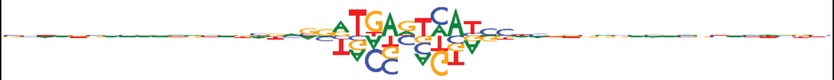 | Fos, Jund, Fosb, Smarcc1 | 160 |
| 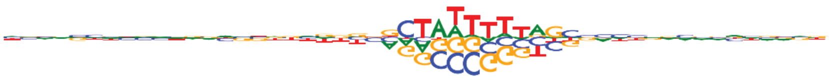 | Mef2a, Mef2c, Mef2d, Mef2b | 109 |
| 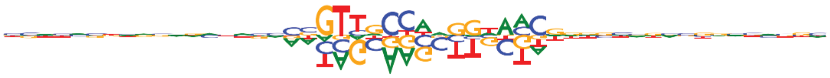 | Rfx1, Rfx5, Rfx4, Arid2, Rfx2, Rfx7, Rfx3 | 43 |
| 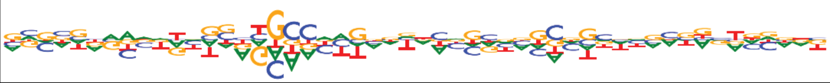 | | 32 |
| 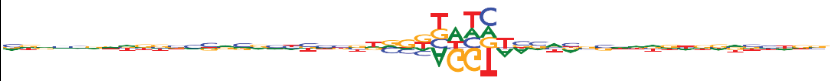 | | 38 |
| 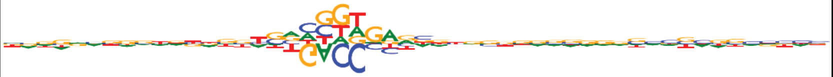 | Ppara, Rxrb, Nr2f2, Rara, Nr2f1, Nr2c1, Rxrg, Esrra, Nr2f6 | 24 |

d)

| TF-MoDISco Motif | TFs with Similar Motif | Seqlets |
|---|---|---|
| 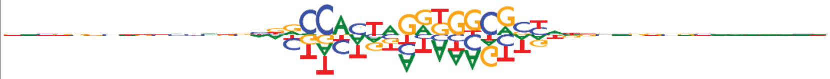 | Ctcf, Ctcfl | 301 |
| 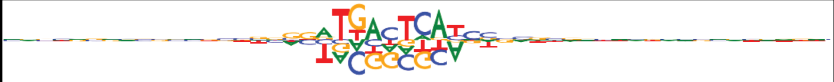 | Fos, Jund, Fosb, Smarcc1 | 175 |
| 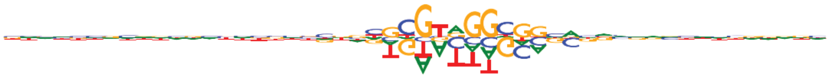 | Egr2, Hif3a, Egr3, Bcl6, E2f3, Ets1 | 167 |
| 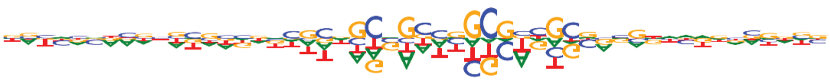 | | 53 |
| 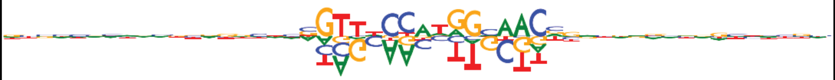 | Rfx1, Rfx2, Arid2, Rfx4, Rfx7, Rfx5, Rfx3 | 47 |
| 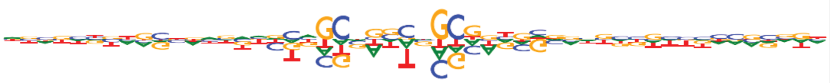 | | 31 |
| 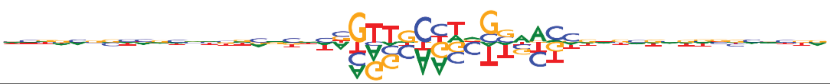 | Rfx5, Rfx4, Rfx7, Rfx8, Rfx3, Rfx1, Stat1 | 29 |
| 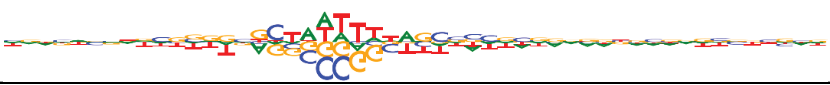 | Mef2c, Mef2a, Mef2d, Mef2b | 55 |

**e)**

| TF-MoDISco Motif | TFs with Similar Motif | Seqlets |
|---|---|---|
| 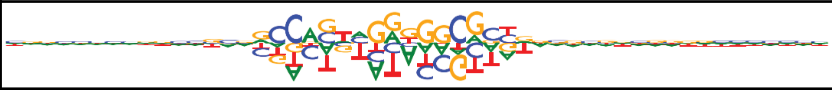 | Ctcf, Ctcfl | 250 |
| 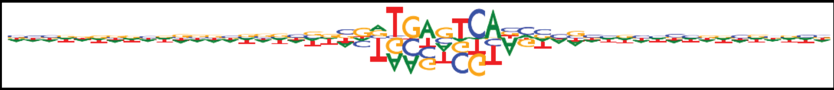 | Fos, Fosb, Jund, Smarcc1 | 110 |
| 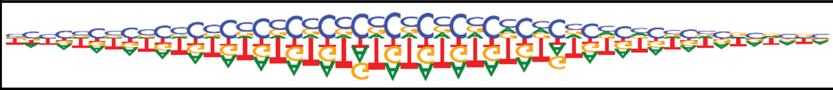 | | 80 |
| 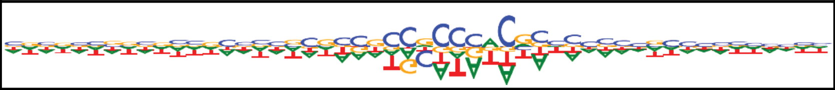 | Egr2, Egr3, Zfp148, Maz, E2f3, Zfp281 | 58 |
| 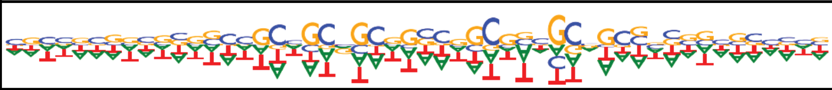 | | 75 |
| 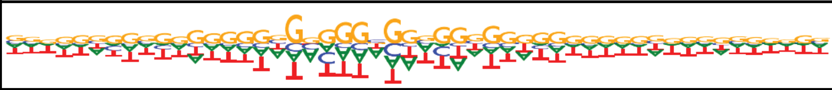 | Maz, Bcl6, Sp3, Zfp148, Zfp281, Sp2, Wt1, E2f1, E2f3, Rreb1, Zbtb7a, Zfp219, E2f6, Klf15, Plag1 | 54 |
| 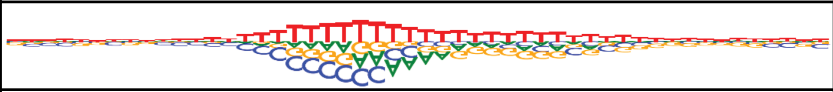 | | 28 |
| 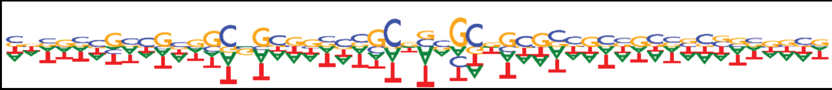 | | 35 |
| 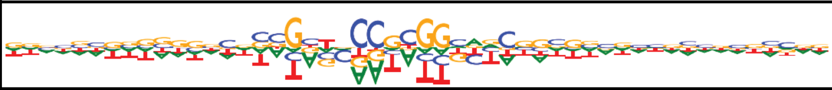 | Rfx1, Arid2, Rfx2, Rfx4, Rfx5, Rfx7 | 27 |
| 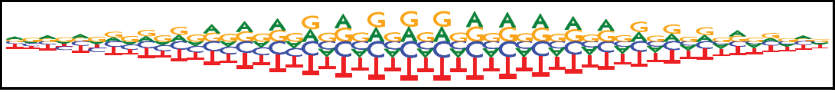 | Bcl6 | 22 |

**f)**

| TF-MoDISco Motif | TFs with Similar Motif | Seqlets |
|---|---|---|
| 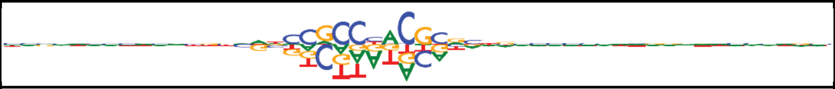 | Egr2, Bcl6, Maz, Hif3a, Egr3, Rreb1, Zpf148, Zfp281, Ets1, Egr1, E2f3, Sp3, Sp1, Sp4, Zpf740 | 250 |
| 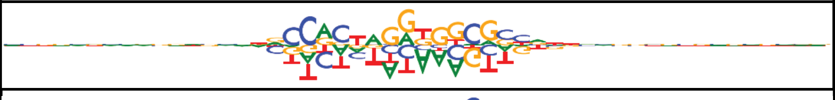 | Ctcf, Ctcfl | 110 |
| 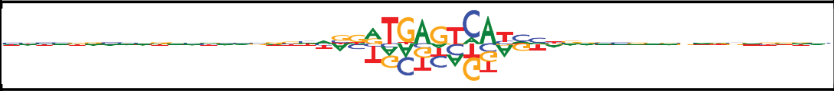 | Fos, Fosb, Jund, Smarcc1, Junb, Bach1 | 80 |
| 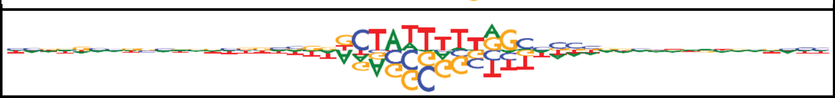 | Mef2a, Mef2c, Mef2d, Mef2b | 58 |
| 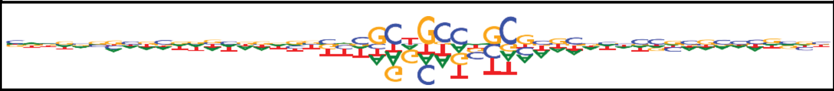 | | 75 |
| 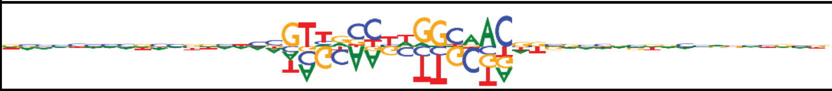 | Rfx1, Rfx2, Arid2, Rfx4, Rfx7, Rfx5, Rfx3 | 54 |
| 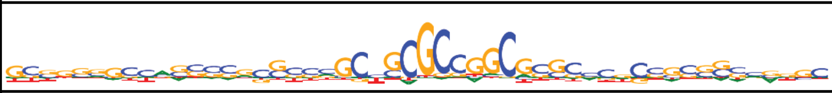 | | 28 |
| 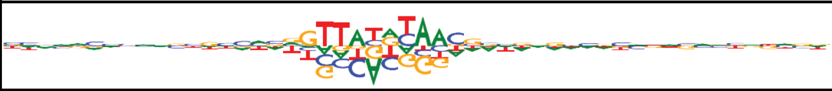 | Dbp, Atf4, Tef, Nfil3 | 35 |
| 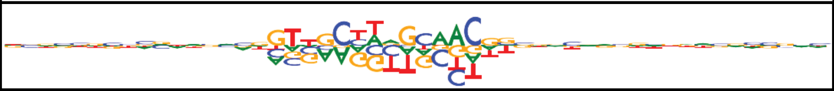 | Rfx5, Rfx8, Rfx4, Rfx7 | 27 |
| 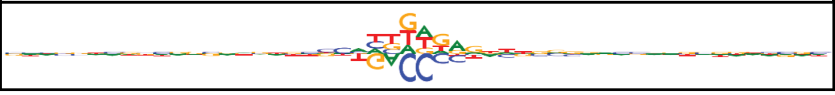 | Thra | 22 |
