## SupplementalFigure8 for "Inferring mammalian tissue-specific regulatory conservation by predicting tissue-specific differences in open chromatin"

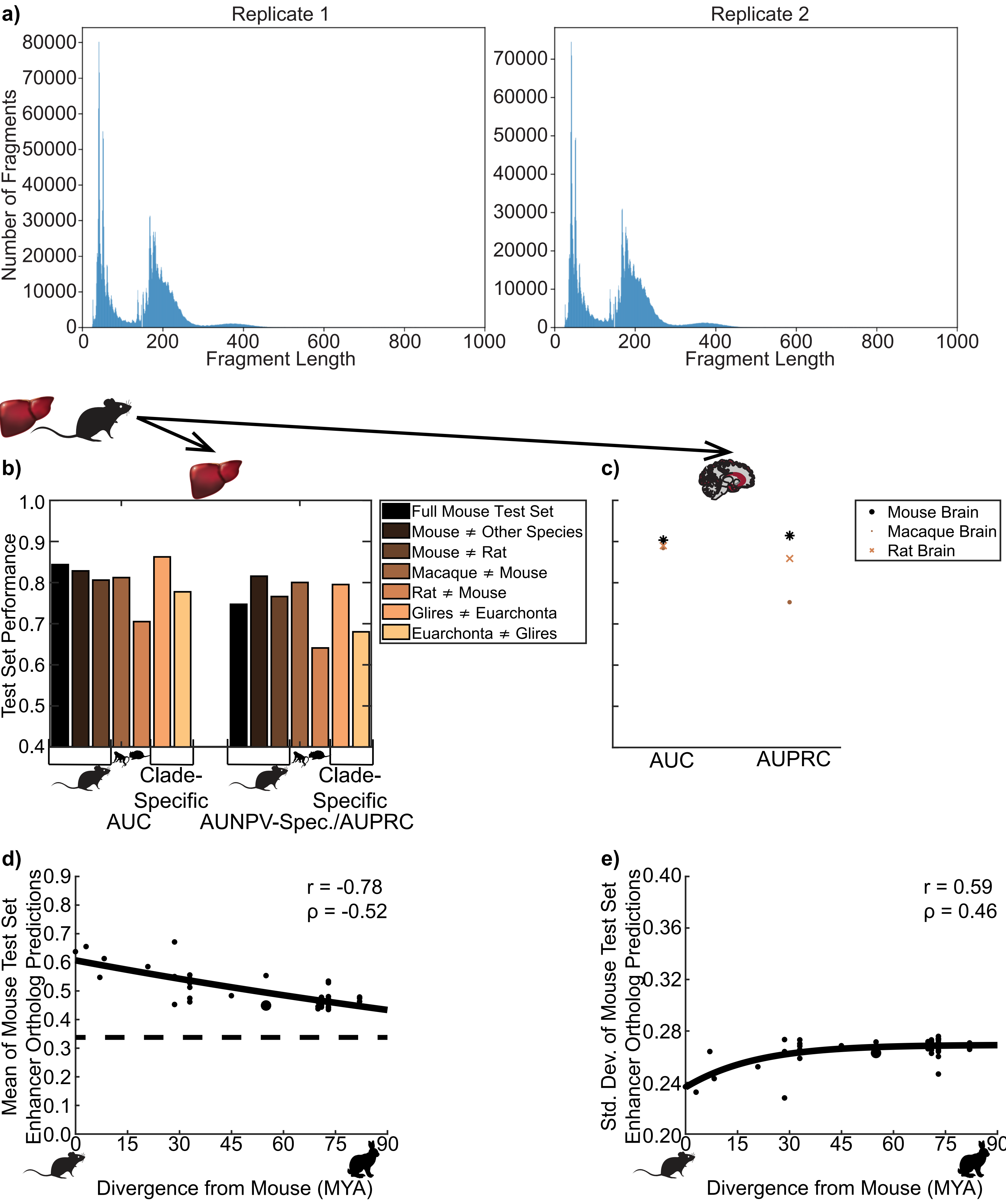


### Supplemental Figure 8

f)

| TF-MoDISco Motif | TFs with Similar Motif | Seqlets |
|---|---|---|
| 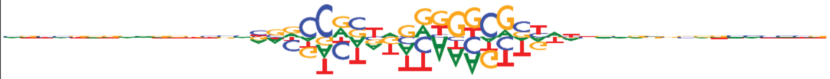 | Ctcf, Ctcfl | 291 |
| 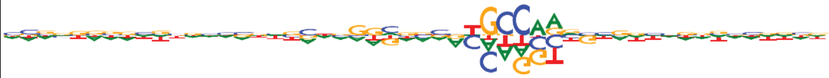 |  | 123 |
| 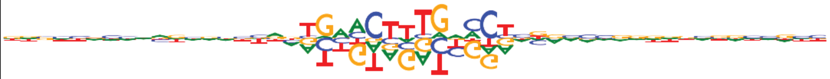 | Nr4a3, Rxra, Hnf4g, Nr1h2, Nr4a2, Ppard, Nr4a1, Ppara | 63 |
| 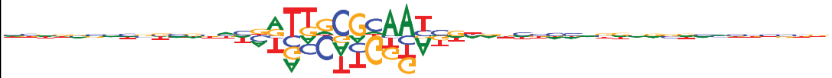 | Cebpb, Cebpg, Cebpa, Cebpd, Tef, Dbp, Nfil3, Hlf, Cebpe, Jdp2 | 56 |
| 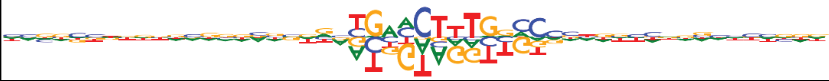 |  | 36 |
| 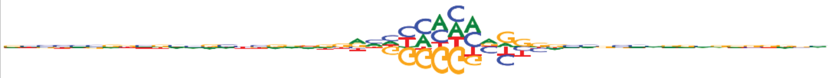 | Zbtb7a, Wt1, Plagl1, Zfp148, Sp2, Klf15, Plag1 | 51 |
