## SupplementalFigure9 for "Inferring mammalian tissue-specific regulatory conservation by predicting tissue-specific differences in open chromatin"

### Supplemental Figure 9

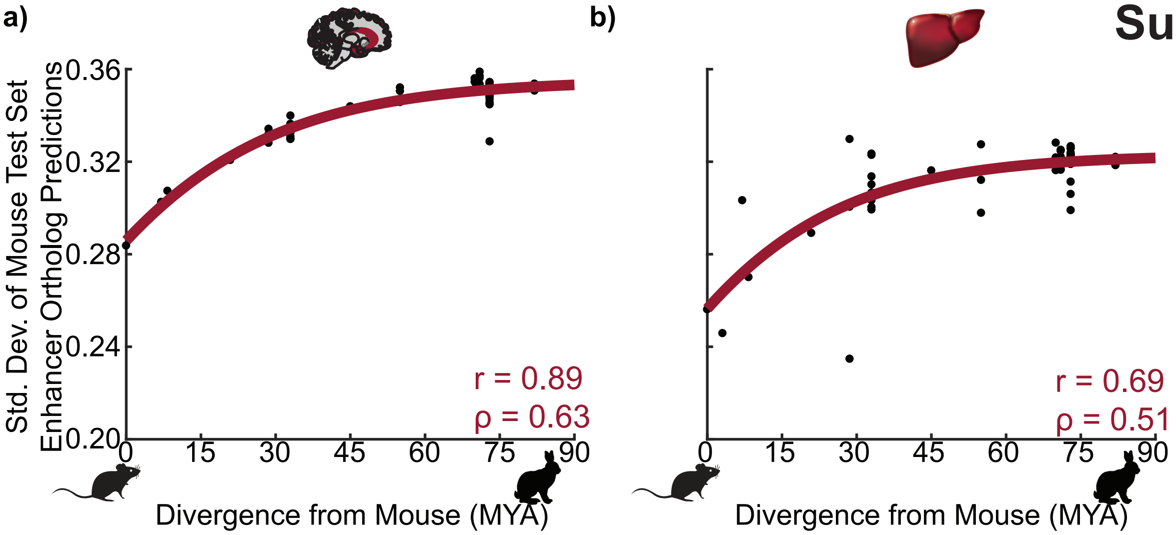


c)

| Brain TF-MoDISco Motif | TFs with Similar Motif | Seqlets |
|---|---|---|
| 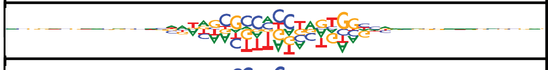 | Ctcf, Ctcfl | 1052 |
| 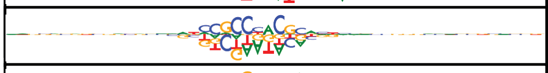 | Egr2, Egr3, Bcl6, Hif3a | 668 |
| 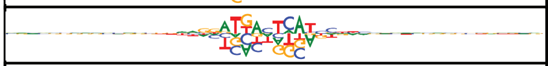 | Fos, Smarcc1, Fosb | 515 |
| 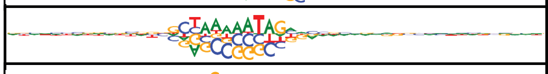 | Mef2a, Mef2c, Mef2d, Mef2b, Tead4 | 220 |
| 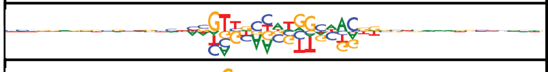 | Rfx1, Rfx2, Arid2, Rfx5, Rfx4, Rfx7 | 156 |
| 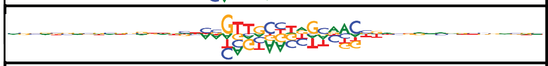 | Rfx8, Rfx5, Rfx7 | 72 |
| 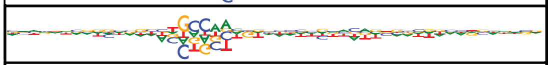 | | 57 |
| 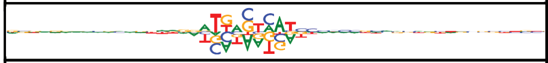 | | 51 |
| 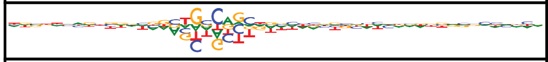 | | 48 |
| 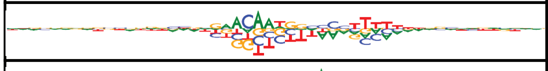 | | 34 |
| 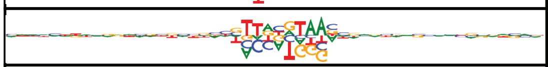 | Hlf, Tef, Dbp, Nfil3, Cebpb | 44 |
| 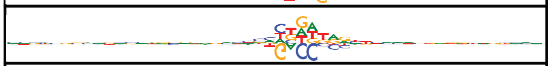 | Gli3, Glis3, Rxrb, Gli1, Rxrg, Nr1i3, Zic5, Gli2 | 93 |
| 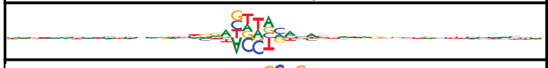 | | 82 |
| 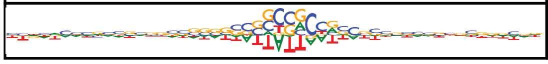 | Onecut1, Onecut3, Hmg20b, Foxl1, Pit1 | 52 |

**d)**

| Liver TF-MoDISco Motif | TFs with Similar Motif | Seqlets |
|---|---|---|
| 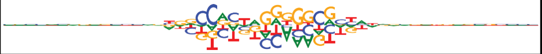 | Ctcf, Ctcfl | 1161 |
| 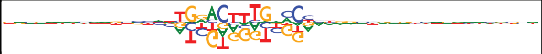 | Hnf4g, Nr1h2, Rxra, Hnf4a, Nr4a3, Pparg | 839 |
| 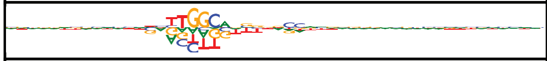 | | 665 |
| 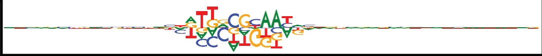 | Cebpg, Cebpb, Cepbd, Cebpa, Cebpe, Dbp, Tef, Hlf, Nfil3 | 663 |
| 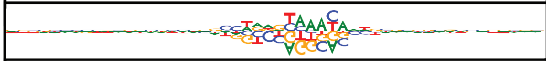 | Foxa2, Foxf2, Foxp4, Foxc1, Foxc2, Foxb1, Foxa3, Foxa1, Foxj3, Foxd1, Foxp2, Foxo3, Foxn3 Foxl1, Foxp1, Foxk1, Foxj2, ENSMUSG00000090020, Foxg1, Foxd2, Foxd3, Gm5294, Foxj1 | 496 |
| 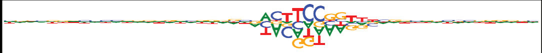 | Ets1, Erg, Sfpi1, Bcl6, Spib, Fli1, Bcl11a, Elk1, Etv6, Etv1, Elk3, Etv4, Erf, Etv3, Gm5454, Gm4881, Elk4, Etv5, Elf4, Fev, Etv2, Elf2, Gabpa, Ets2, Spic, Elf3, Ehf, Maz, Prdm1, Elf5 | 356 |
| 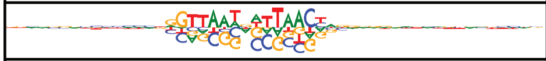 | | 278 |
| 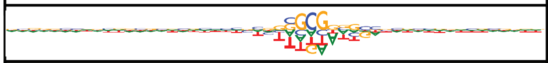 | Sp3, Sp2, Zbtb7a, Zfp148, Wt1, Maz, Zfp281, Klf16, Klf5, Egr4, Sp1, Sp5, Zfx, E2f1, Klf6, Sp8, Klf2, Klf7, Klf14, Klf4, Klf15, Zfp219, Klf8, Klf12, Sp4, Egr2, E2f3, Tcfap2c, Mbd2, Rreb1 | 266 |
| 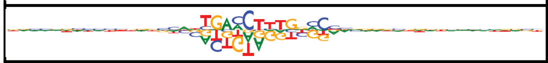 | Hnf4g, Nr4a3, Ppara, Nr4a2, Rxra, Nr1h2, Nr4a1, Ppard, Pparg | 215 |
| 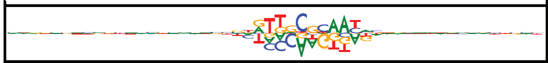 | | 181 |
| 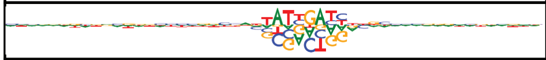 | Onecut3, Onecut1 | 178 |
| 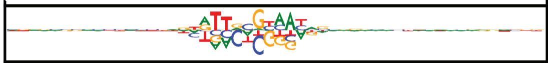 | Bcl6 | 138 |
| 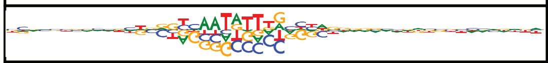 | | 119 |
| 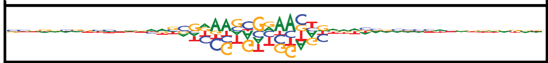 | Irf1, Stat2, Prdm1, Bcl11a, Bcl6, Sfpi1, Irf2 | 95 |
| 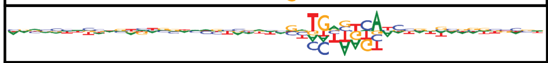 | Bach1, Fosb, Nfe2l2 | 35 |
| 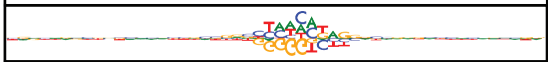 | | 132 |
| 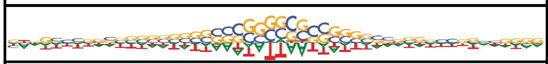 | Phf21a, Arid3b, Dbx1, Tlx2, Lhx3, Pou3f4, Pit1, Lhx5, Onecut1, Pou3f1, Lmx1b, Lhx1, Pouf43, Pou1f1, Hmg20b, Onecut3, Pou2f1, Lmx1a, Pou4f1 | 90 |
| 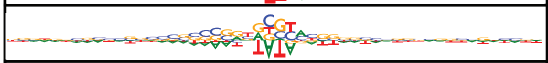 | Zfp637 | 38 |
| 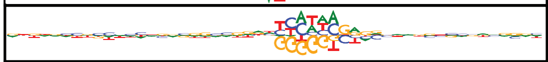 | | 35 |
| 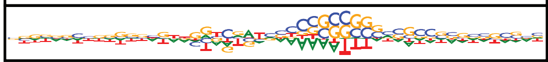 | | 28 |
